# Partitioning the genetic architecture of amyotrophic lateral sclerosis

**DOI:** 10.1101/505693

**Authors:** Iris J. Broce, Chun C. Fan, Nicholas T. Olney, Catherine Lomen-Hoerth, Steve Finkbeiner, Nazem Atassi, Merit E. Cudkowicz, Sabrina Paganoni, Jennifer S. Yokoyama, Aimee Kao, William P. Dillon, Christine M. Glastonbury, Christopher P. Hess, Wouter van Rheenen, Jan H. Veldink, Ammar Al-Chalabi, Ole A. Andreassen, Anders M. Dale, William W. Seeley, Leo P. Sugrue, Aaron Ofori-Kuragu, Celeste M. Karch, Bruce L. Miller, Rahul S. Desikan

## Abstract

The genetic basis of sporadic amyotrophic lateral sclerosis (ALS) is not well understood. Using large genome-wide association studies and validated tools to quantify genetic overlap, we systematically identified single nucleotide polymorphisms (SNPs) associated with ALS conditional on genetic data from 65 different traits and diseases from >3 million people. We found strong genetic enrichment between ALS and a number of disparate traits including frontotemporal dementia, coronary artery disease, C-reactive protein, celiac disease and memory function. Beyond *C9ORF72*, we detected novel genetic signal within numerous loci including *GIPC1, ELMO1* and *COL16A* and confirmed previously reported variants, such as *ATXN2, KIF5A, UNC13A* and *MOBP.* We found that ALS variants form a small-world co-expression network characterized by highly inter-connected ‘hub’ genes. This network clustered into smaller sub-networks, each associated with a unique function. Altered gene expression of several sub-networks and hubs was over-represented in neuropathological samples from ALS patients and SOD1 G93A mice. Our collective findings indicate that the genetic architecture of ALS can be partitioned into distinct components where some genes are highly important for developing disease. These findings have implications for stratification and enrichment strategies for ALS clinical trials.

## Introduction

Sporadic amyotrophic lateral sclerosis (ALS) is a fatal neurodegenerative disease characterized by progressive muscle paralysis from selective loss of upper and lower motor neurons. Spreading rapidly, ALS can lead to respiratory failure and death in 3-5 years^1^. Given the paucity of disease modifying treatments, elucidating the genetic basis of ALS can delineate putative pharmacological targets and highlight molecular mechanisms underlying disease. Importantly, refining the genetic landscape of ALS can inform cohort stratification and enrichment strategies for clinical trials.

ALS is increasingly recognized as a complex disorder with an incompletely understood genetic architecture. Prior work suggests that sporadic ALS is genetically characterized by a few rare variants, each explaining a substantial portion of the inherited risk (oligogenic)^2–4^. However, more recent evidence indicates that low-risk, common variants underlie ALS (polygenic) ^5^. Importantly, several ALS-associated variants have been implicated in other diseases suggesting genetic pleiotropy^6–8^. Furthermore, it is not known whether certain ALS genes are more important than others for influencing disease etiology.

Here, our goal was to elucidate the genetic architecture of ALS by leveraging statistical power from large GWAS from 65 distinct traits and diseases. Using these methods, we have discovered novel genetic risk loci and shown abundant genetic pleiotropy between several neurodegenerative diseases including ALS, FTD, progressive supranuclear palsy (PSP), corticobasal degeneration (CBD), Parkinson’s disease (PD), and Alzheimer’s disease (AD) ^8–12^.

## Results

### Selective shared genetic risk between ALS and 65 distinct traits

Using previously published stratified FDR methods (see Methods), we assessed genetic overlap between ALS and 65 distinct traits and diseases. We identified genetic enrichment in ALS SNPs across different levels of significance with 65 distinct traits and diseases (Fig. 1). Consistent with prior work^8^, we found that the highest level of pleiotropic enrichment was between ALS and FTD (700-fold enrichment). Surprisingly, we also found robust genetic enrichment in ALS SNPs as a function of coronary artery disease (CAD; 300-fold enrichment), memory (225-fold enrichment), C-reactive protein (CRP; 50-fold enrichment), and PSP (50-fold enrichment). We found weaker genetic enrichment with celiac disease (CeD), CBD, body mass index (BMI), rheumatoid arthritis (RA), schizophrenia (SCZ), verbal numeric reasoning (VNR), and putamen volume (PUT). We found no enrichment between ALS and the other phenotypes. We note that these analyses reflect genetic enrichment after removing all SNPs within chromosome 9.

**Fig. 1:**
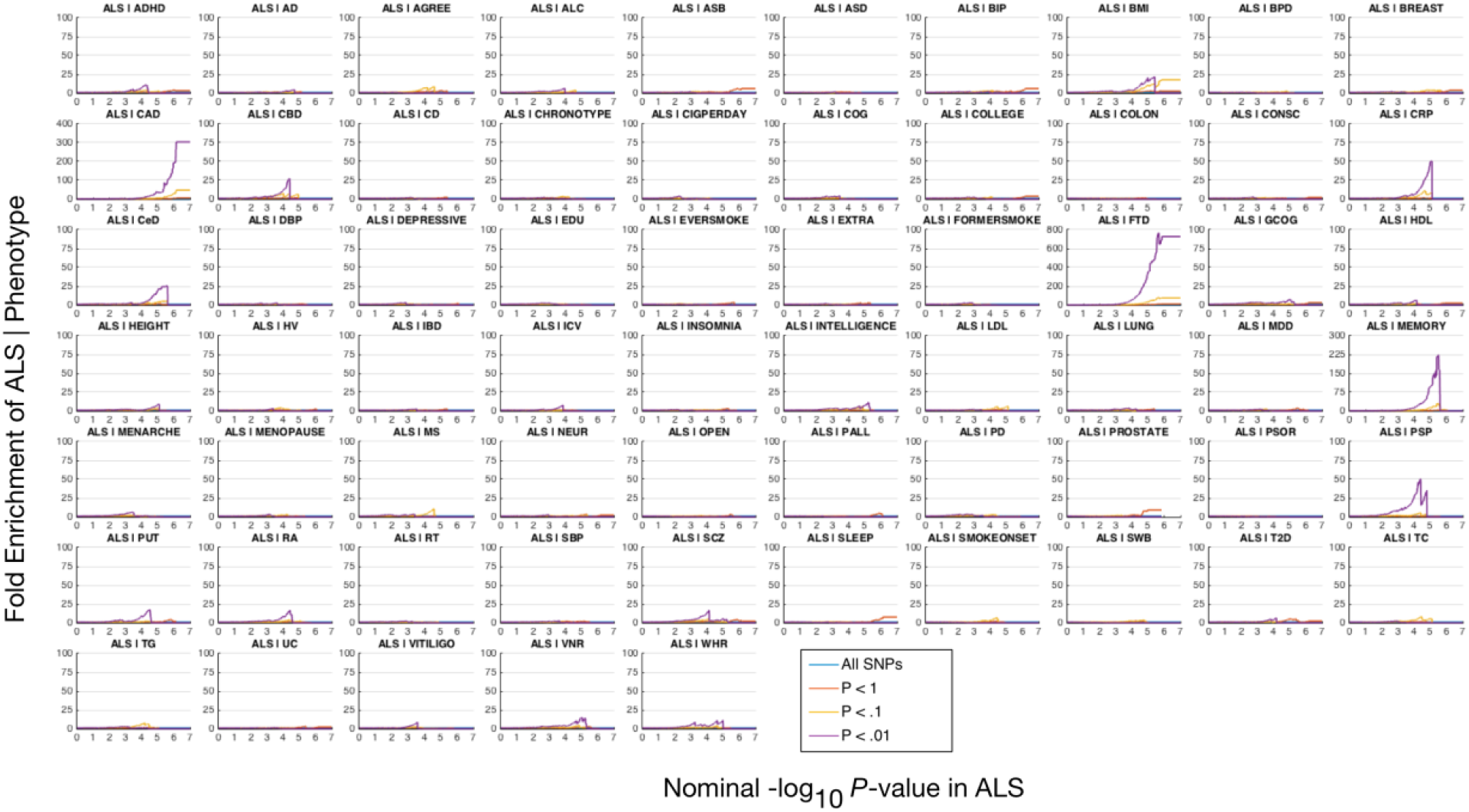
Fold enrichment plots of enrichment versus nominal -log_10_ p-values (corrected for inflation) in Amyotrophic lateral sclerosis (ALS). Fold enrichment plots of enrichment versus nominal -log_10_ p-values (corrected for inflation) in ALS below the standard GWAS threshold of p-value < 5×10^−8^ as a function of significance of association with 65 distinct traits and diseases and at the level of p-value ≤ 1, p-value ≤ 0.1, p-value ≤ 0.01, respectively. Blue line indicates all SNPs.

To identify novel ALS risk loci, we used a stratified approach. First, we computed conditional FDR, a statistical framework that is well suited for gene detection^9,13^. Conditional FDR analysis at a FDR p-value < 0.05 revealed 180 SNPs across 21 chromosomes (Fig. 2, Supplementary Table 1). Next, we performed extensive LD analyses to identify the variants underlying the genetic signals (see Supplemental Information). After accounting for LD, we identified 89 risk loci and annotated each ALS risk SNP with the closest gene(s), resulting in a total of 92 closest genes (Fig. 2, Table 1). Of these, 30 SNPs were either previously reported or were in LD with a previously reported SNP (Table 1, Supplementary Fig. 1). An additional 59 SNPs were novel or were not in LD with SNPs within previously reported loci (Table 1, Supplementary Fig. 1). We found independent hits from SNPs previously reported within *CRIM1, NAF1, TNIPI, PARKIN, ELP3, MRSA, C9ORF72, PCDH9, A2BP1*, and *CPNE7* (Table 1).

**Fig. 2:**
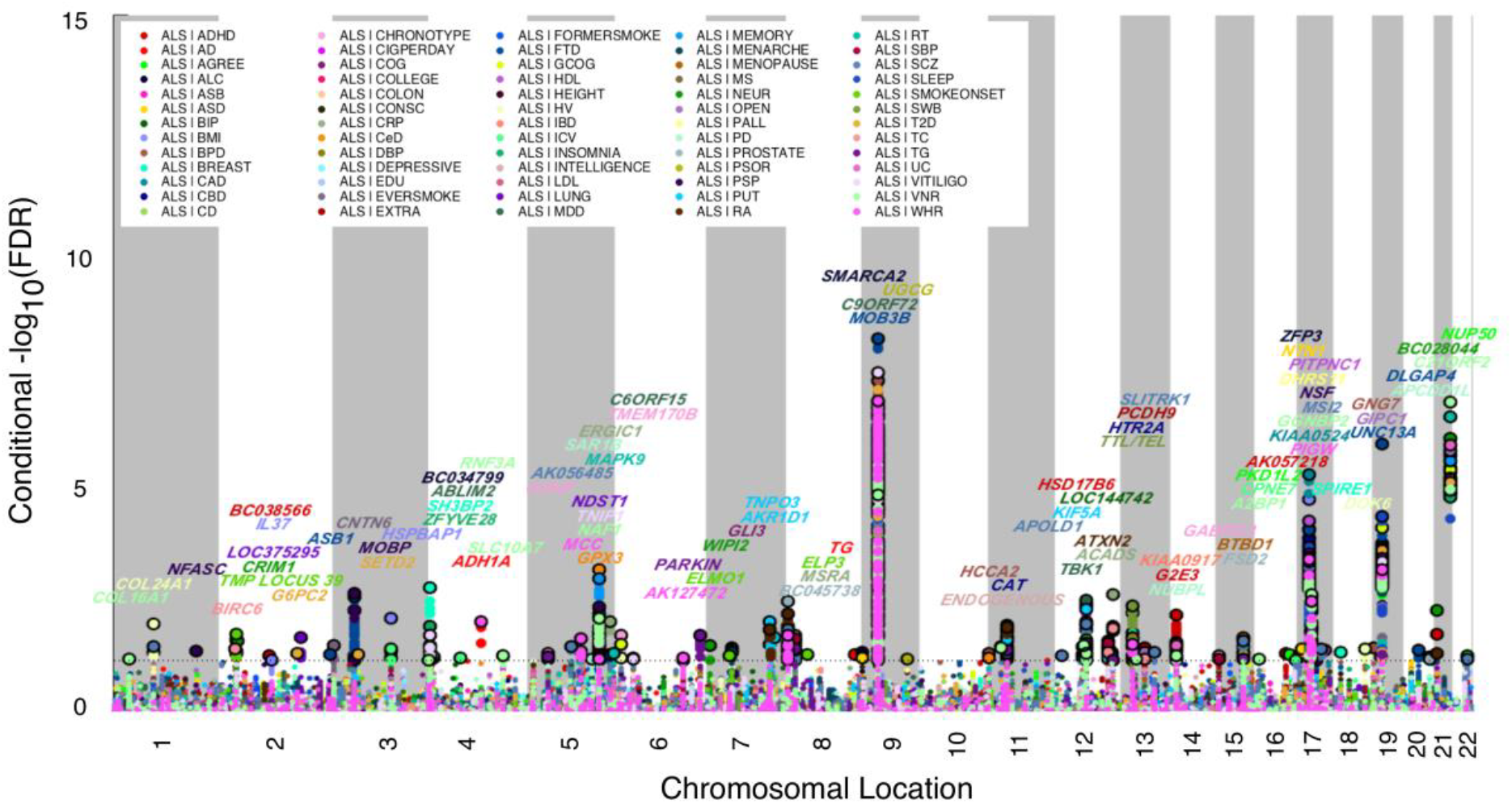
‘Conditional’ Manhattan plot of conditional –log_10_ (FDR) values for Amyotrophic lateral sclerosis (ALS) as a function of 65 distinct traits and diseases. SNPs with conditional –log_10_ FDR > 1.3 (i.e. FDR < 0.05) are shown with large points. A black line around the large points indicates the most significant SNP in each LD block and this SNP was annotated with the closest gene, which is listed above the symbols in each locus.

**Table 1.**
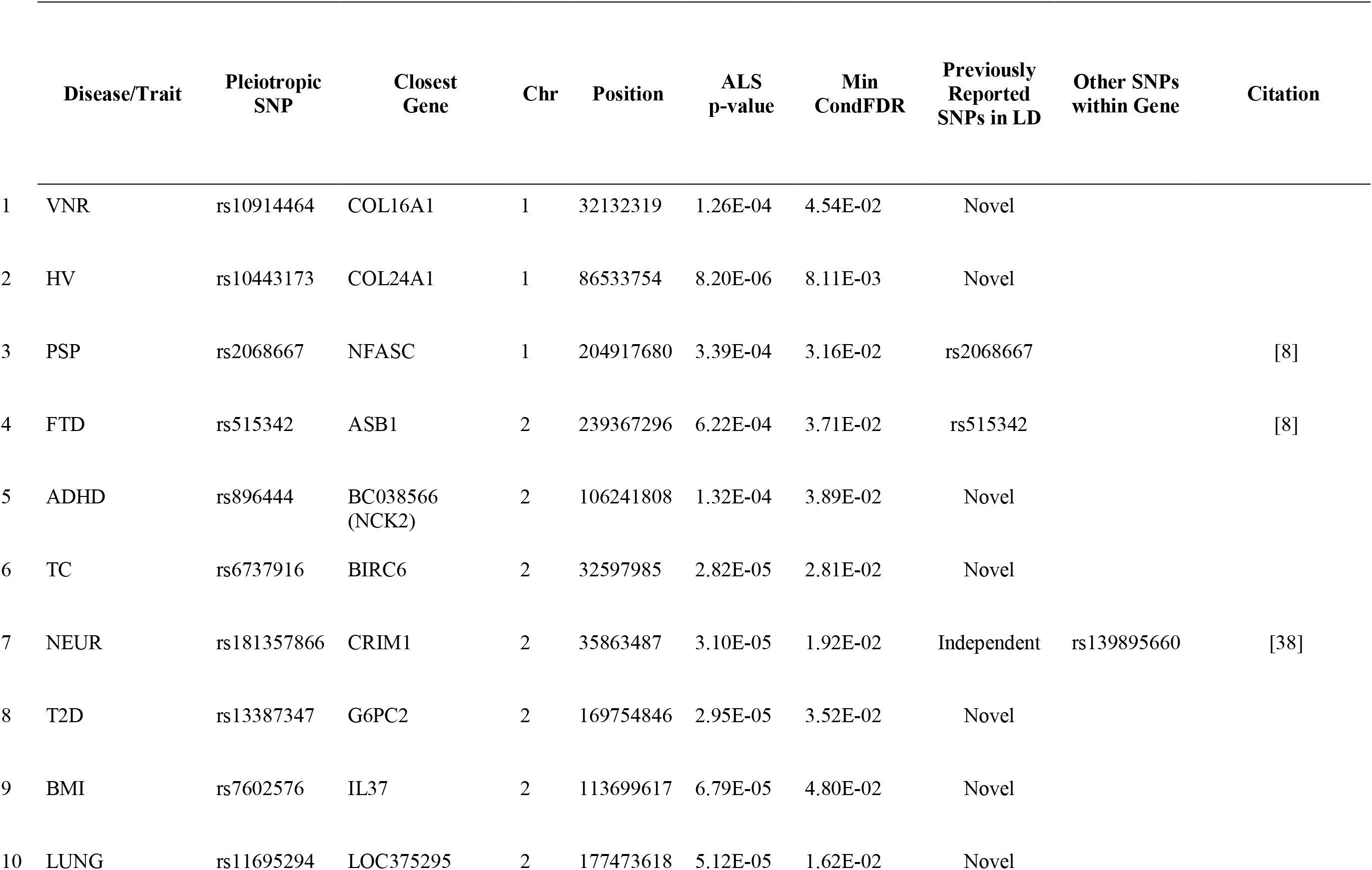
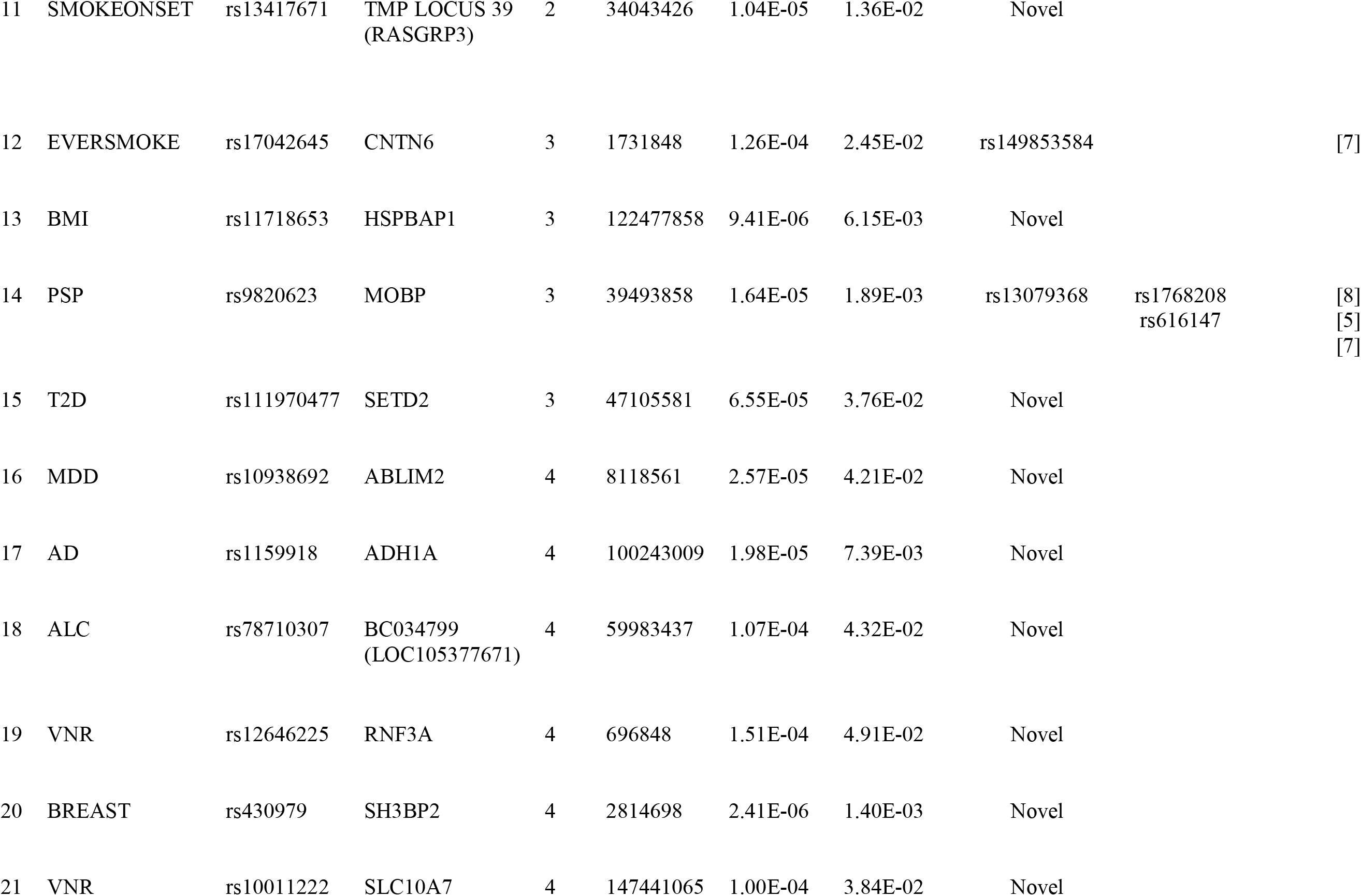
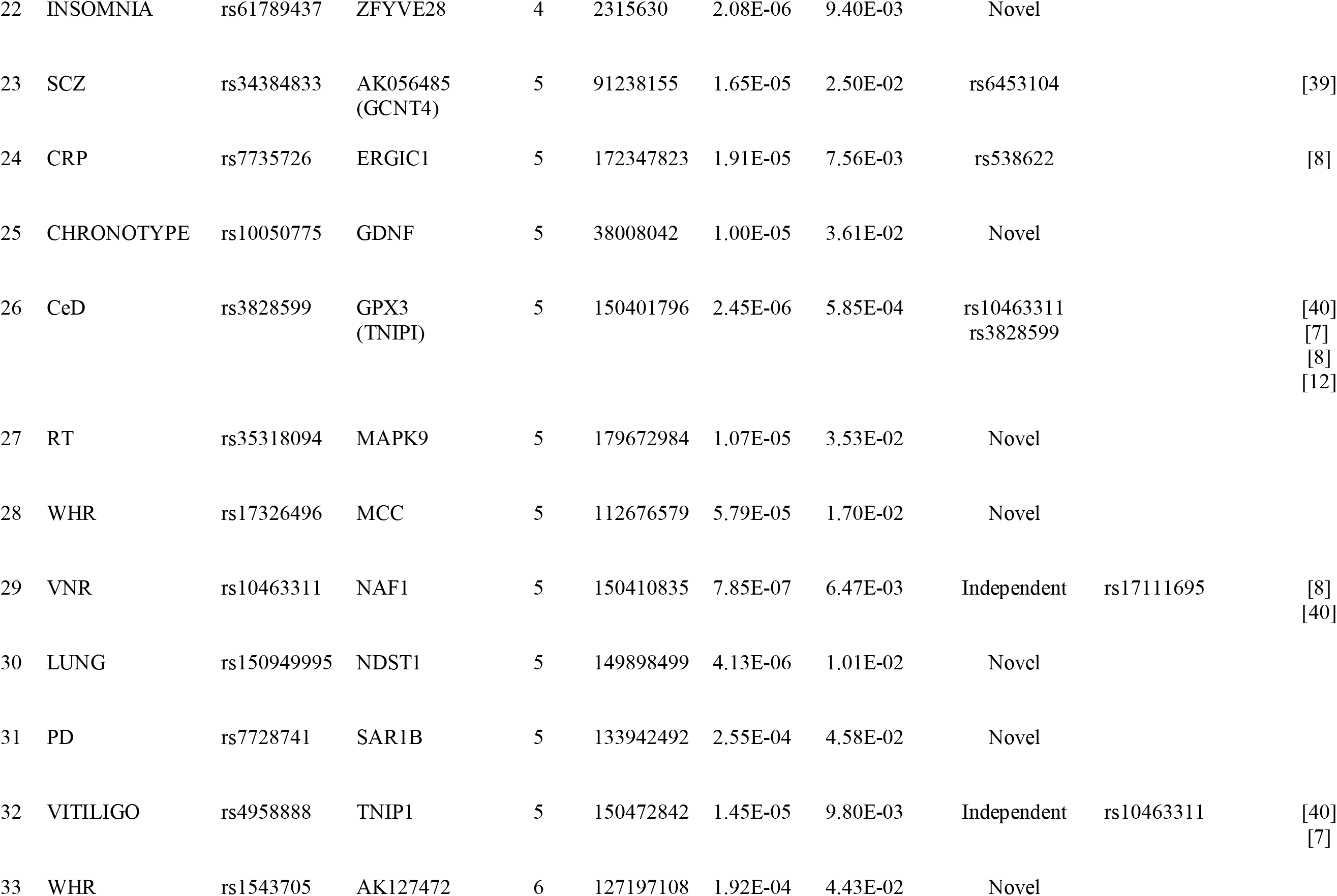
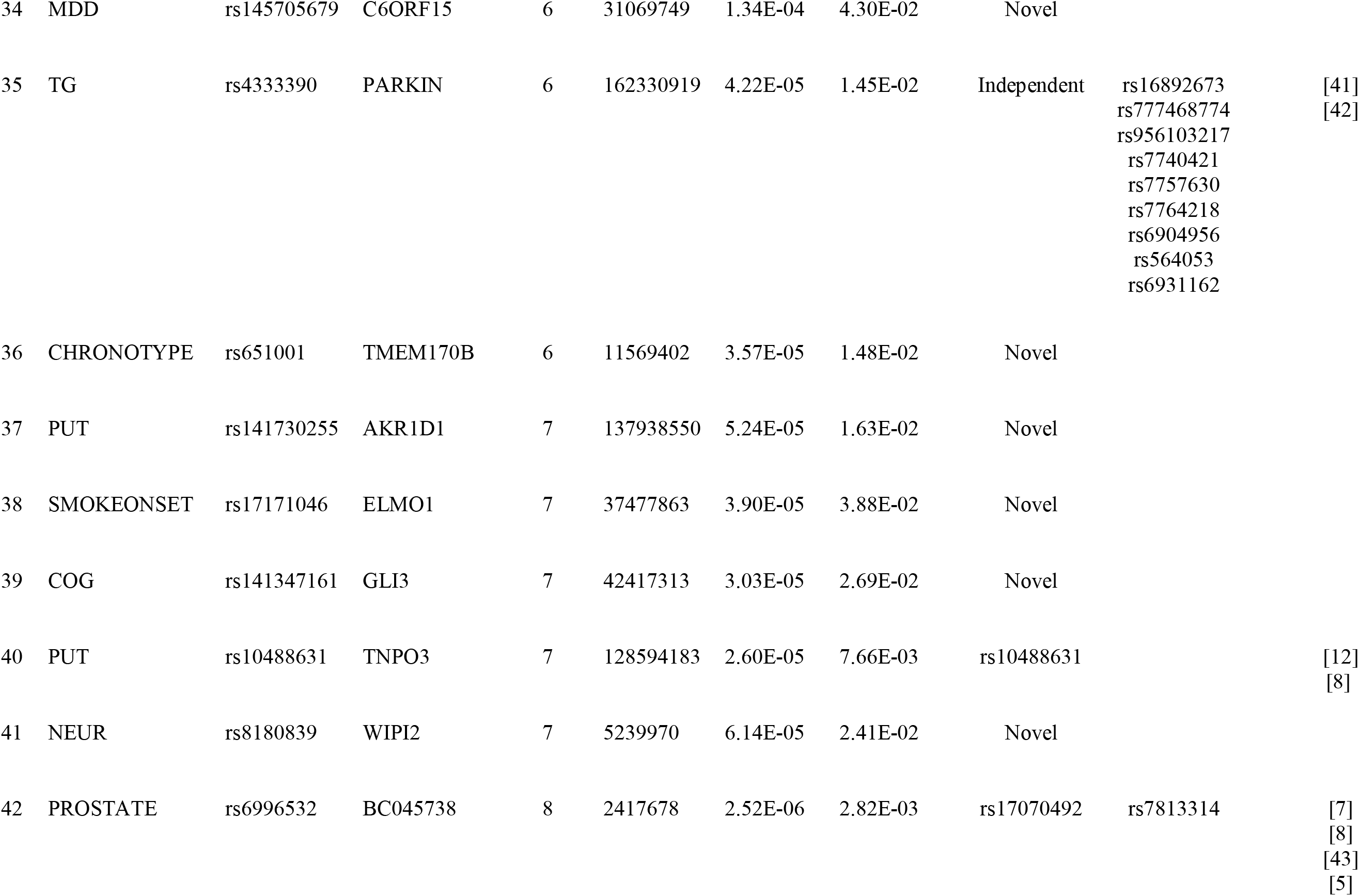
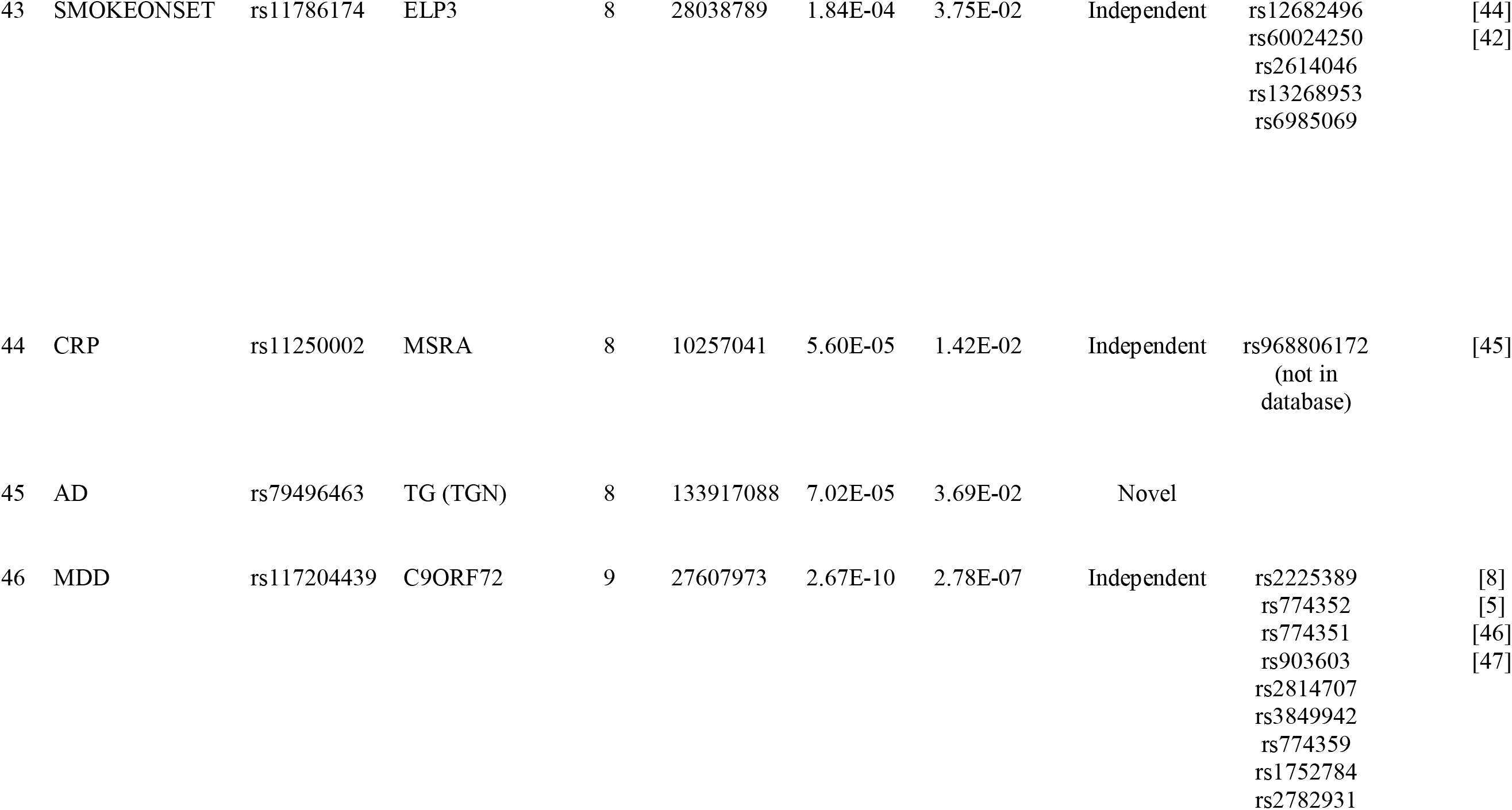
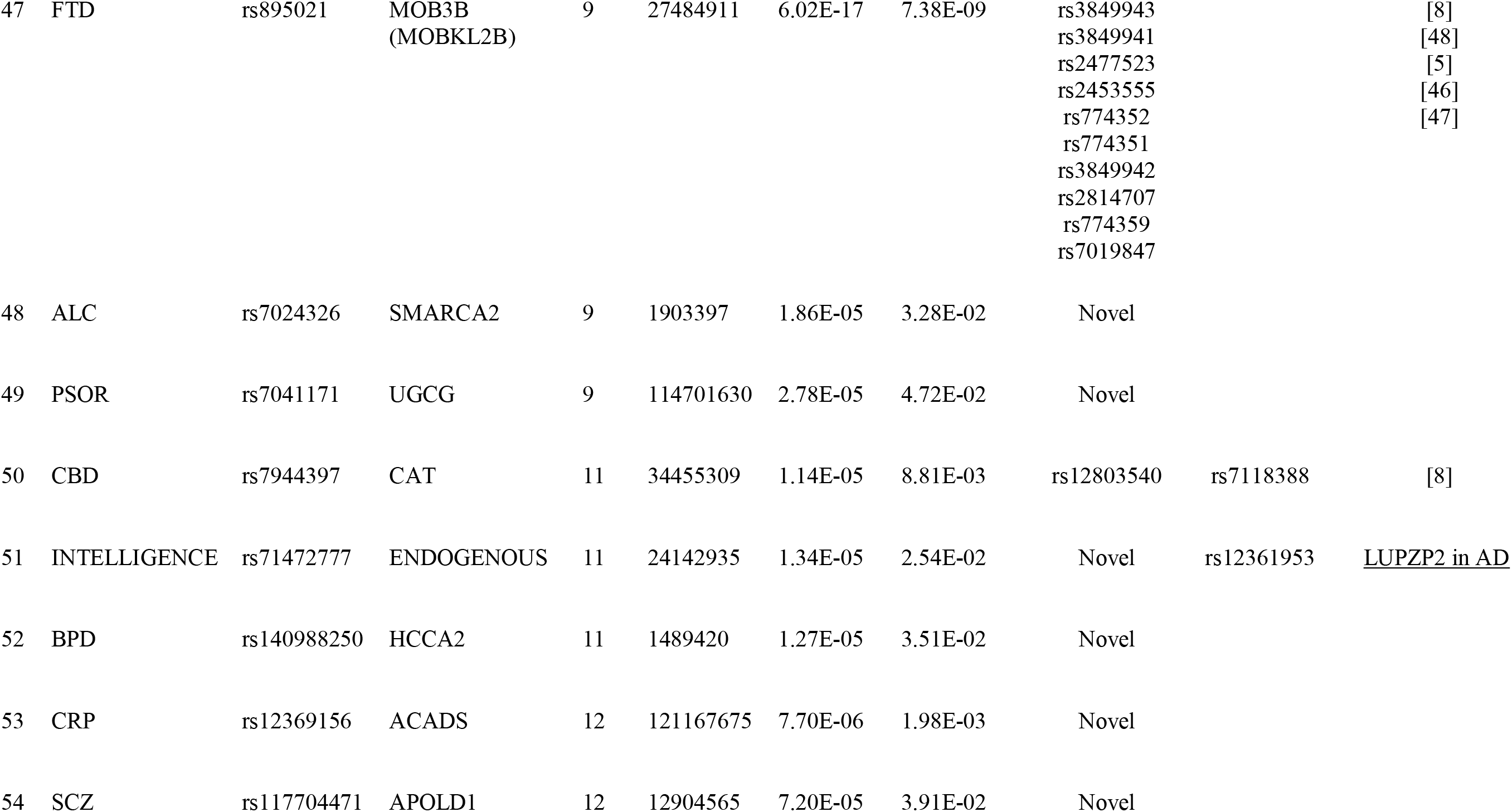
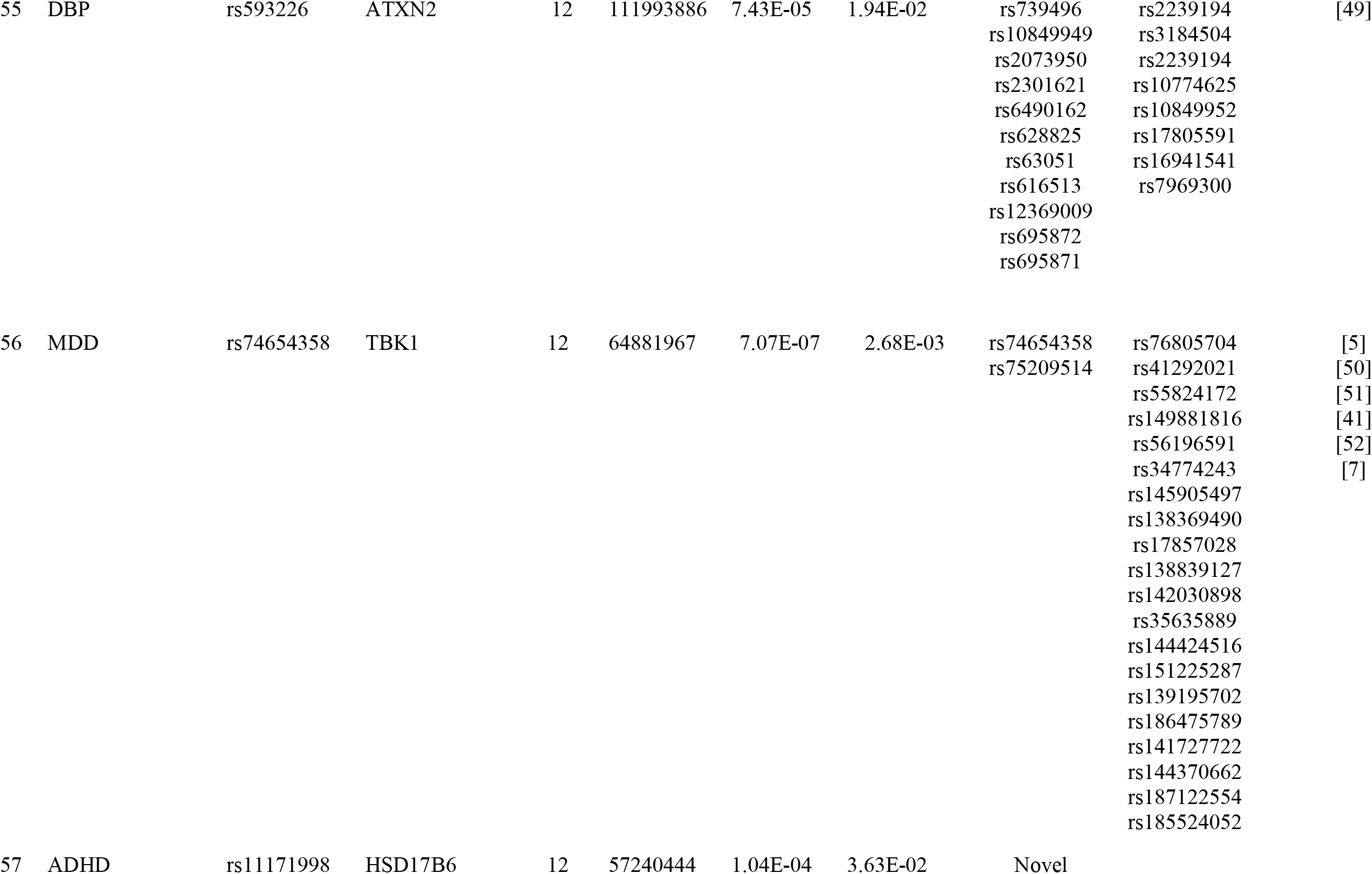
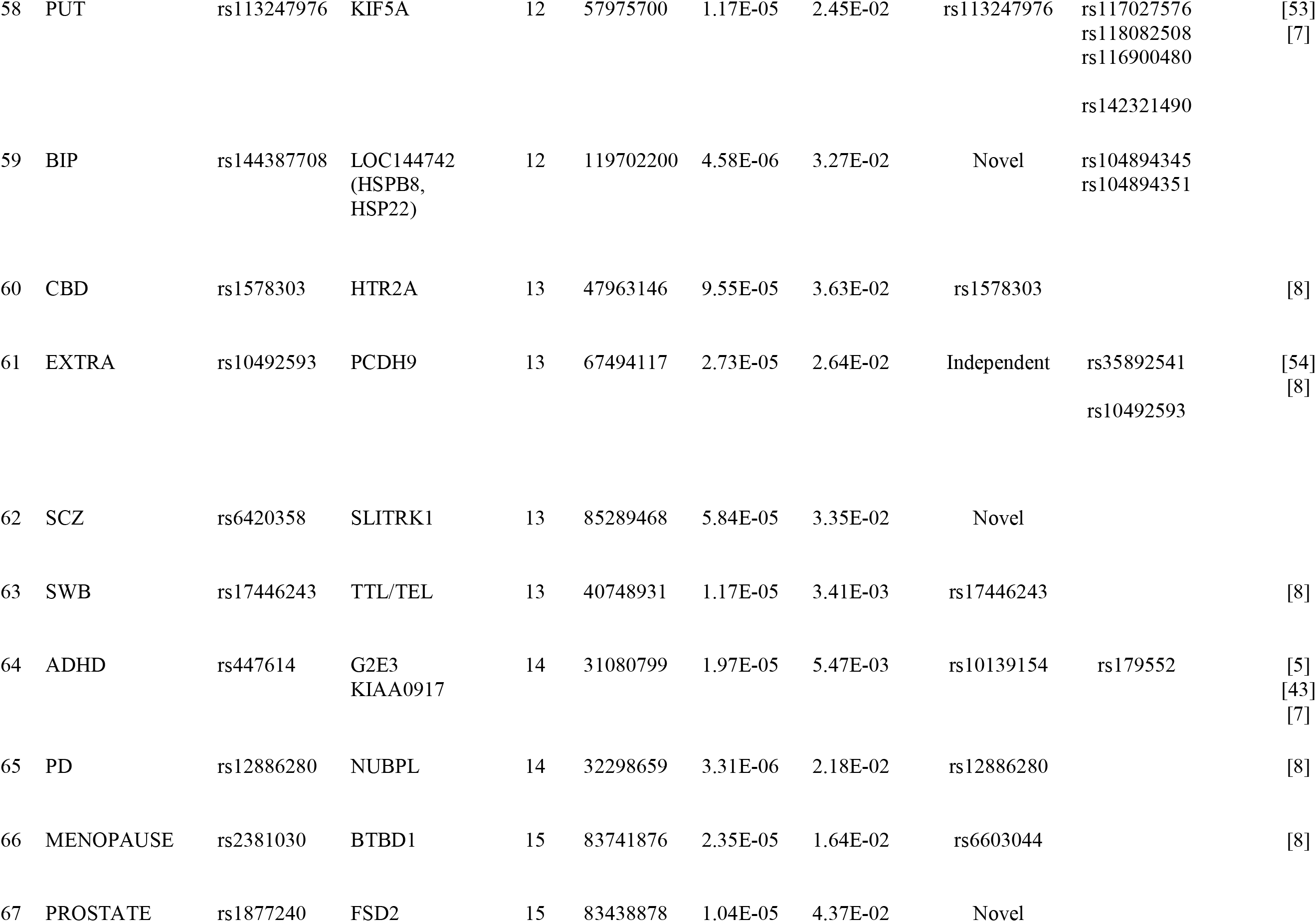
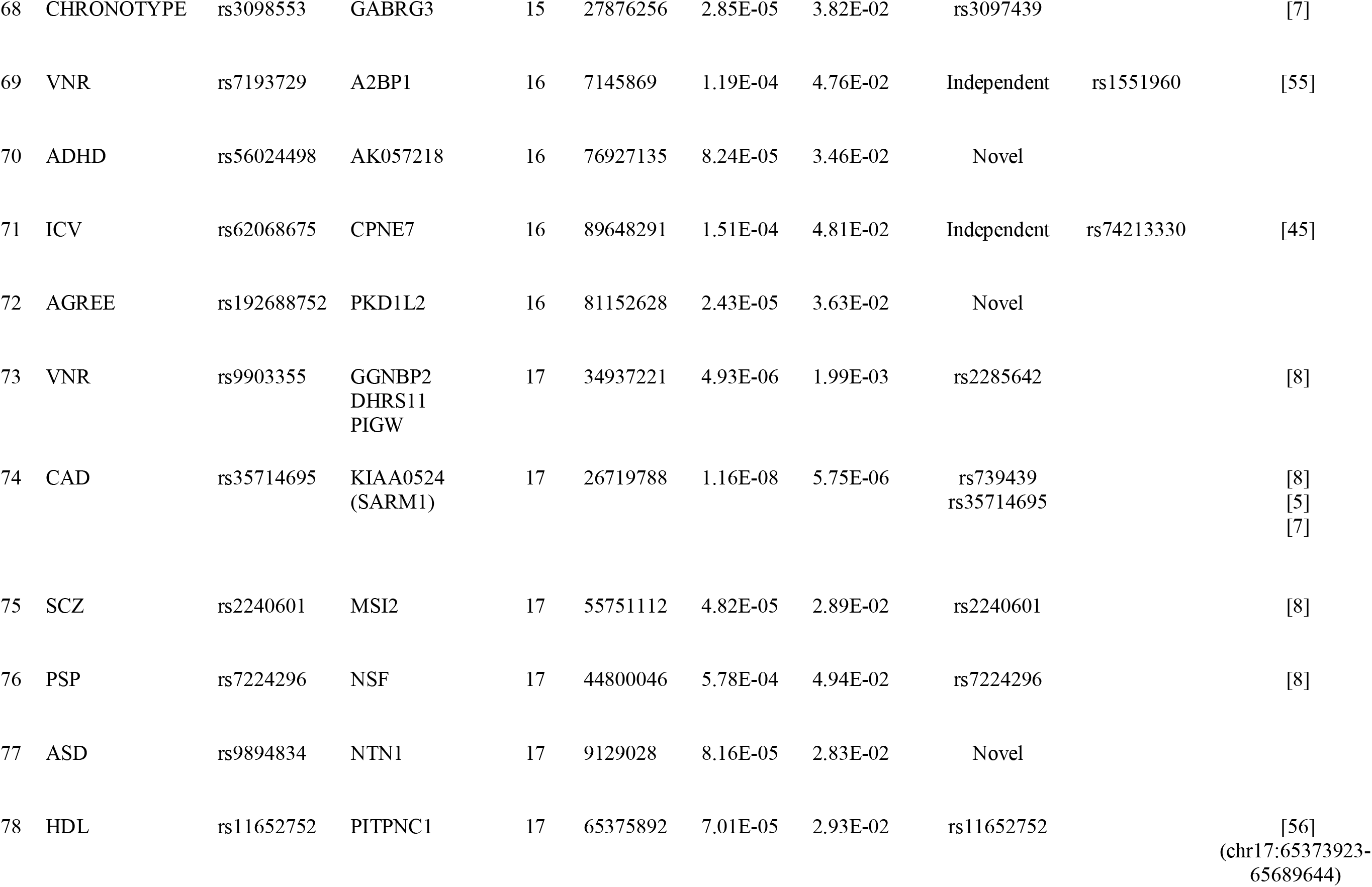
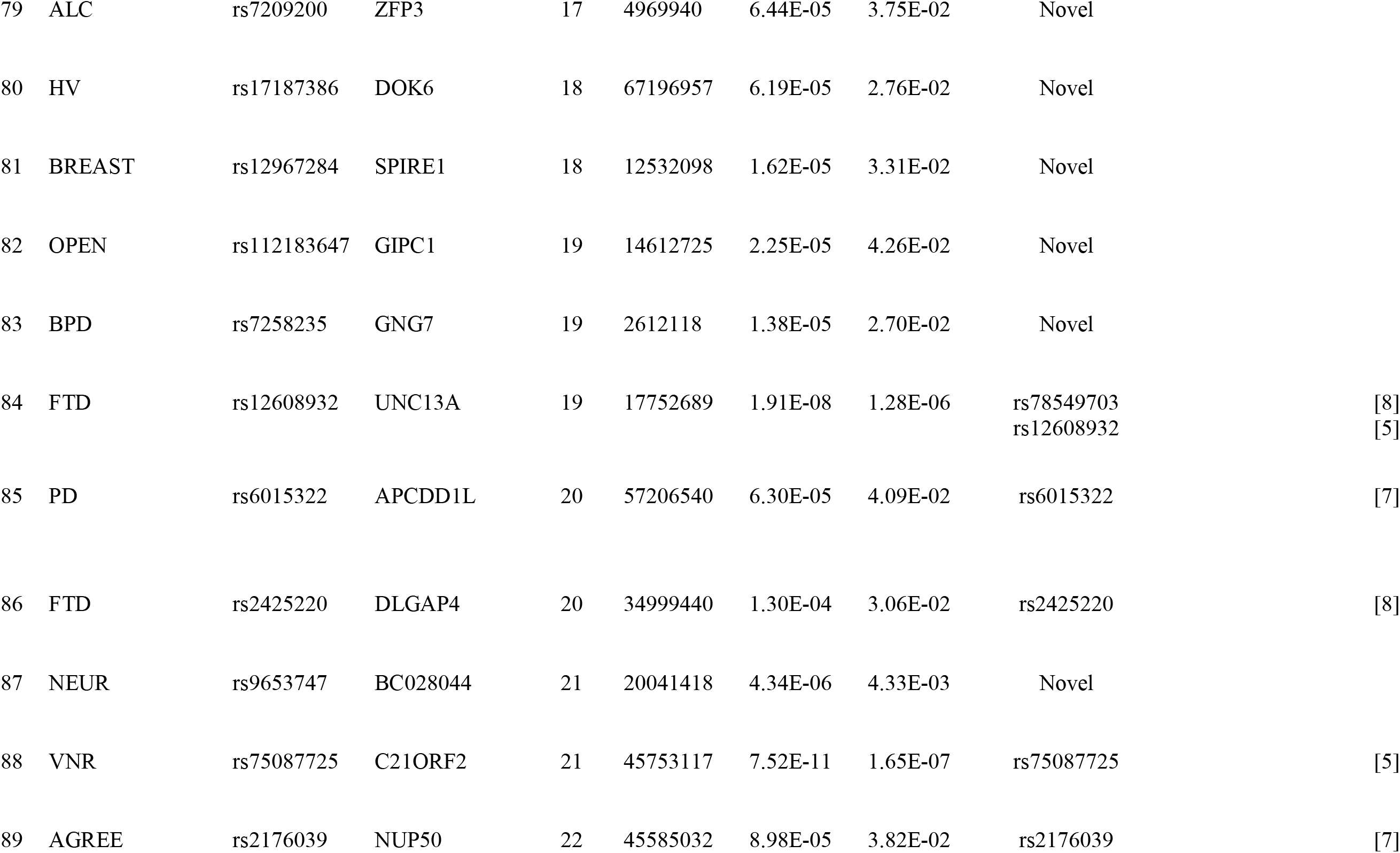

To determine whether the ALS risk genes were associated with a single trait or multiple traits, we plotted the minimum conditional FDR associated with all traits and closest genes. As shown in Supplementary Fig. 2, across all 65 traits and diseases, we found that the ALS risk variants are associated with multiple traits and diseases. Genetic variants within *C9ORF72* were identified on 46 traits. Genetic variants within *KIAA0524* (also known as *SARM1*) were identified on 64 traits, *UNC13A* on 62 traits, *MOB3B* on 58 traits, *TBK1* on 34 traits, and *CAT* and *C21ORF2* on 27 traits.

These findings suggest that ALS has a polygenic component where several genes potentially contribute to disease risk. Genetic pleiotropy with traits like FTD and CAD can be leveraged for ALS gene detection. Importantly, there are several ALS susceptibility loci that are also associated with numerous other traits and diseases.

### cis-eQTL expression

To determine the functional effects of the ALS pleiotropic risk SNPs, we evaluated *cis*-expression quantitative loci (*cis-*eQTL) in human brains free of neuropathology (Supplementary Table 2). In total, the ALS risk SNPs produced significant *cis*-eQTLs (below 1.5 x 10^−3^) within 41 genes. Of these, SNPs within *SMARCA2, GGNBP2, NUP50*, and *TNIP1* showed overlapping annotation between the eQTL and the closest genes. Thirteen SNPs showed significant *cis*-eQTLs with multiple genes.

### Biological networks associated with ALS genetic risk genes

Using GeneMANIA (www.genemania.org), an online web-portal for bioinformatic assessment of gene networks^14^, we conducted a network analysis to explore the interaction and co-expression patterns associated with the ALS risk genes defined as the combination of the 1) closest genes to the SNP and 2) functional genes (i.e., SNPs with significant *cis-*eQTLs). We found that a large number of these genes showed physical protein-protein interactions (42.93%), were co-expressed (29.33%), and showed genetic interactions (13.28%) (Fig. 3). Few ALS genetic risk genes shared pathways (9.36%), were co-localized (2.60%), or predicted functional interactions between genes based on orthology (2.50%) (Supplementary Table 3).

**Fig. 3:**
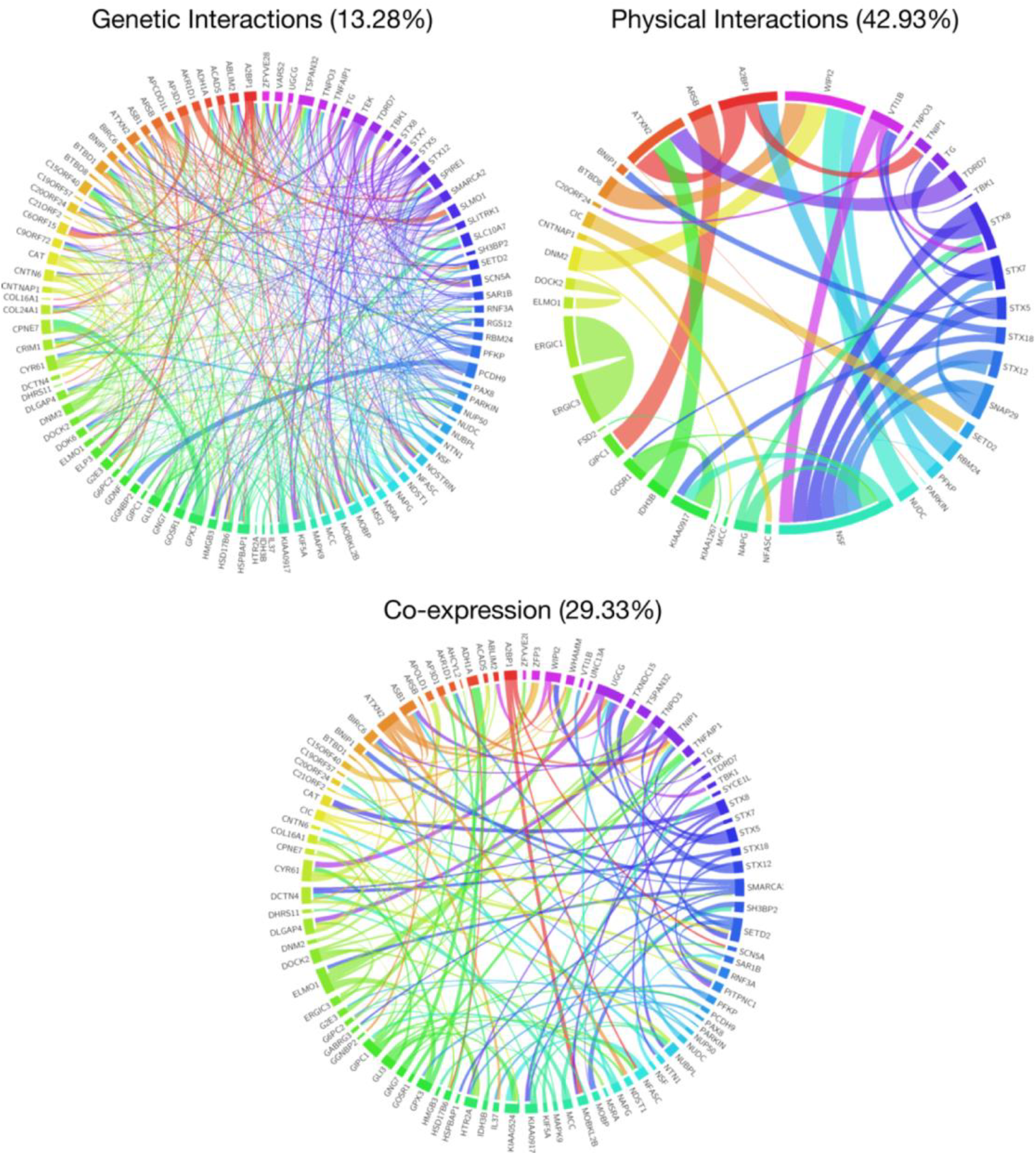
Network interaction graph illustrating genetic interactions, physical interactions, and co-expression patterns associated with the ALS risk genes.

### Properties of the ALS biological networks

We assessed the network structure of the physical protein-protein interaction network, co-expression network, and genetic interactions network. Specifically, we asked whether some genes play a more influential role than others. Most complex networks have a small-world property characterized by relatively short paths between any pair of nodes (genes) ^15^. In small-world networks, perturbing any given node is thought to also perturb neighboring nodes and the entire network in general. Quantitatively, a network is considered small-world if its “small-worldness” index is higher than one (a stricter rule is small-worldness >=3)^16^. Further, the clustering coefficient for the target small-world network should be higher than the clustering coefficient of a comparable random network. Also, the average shortest path length of the target network should be similar or higher (but not substantially higher) than a comparable random network.

First, we evaluated the degree to which each network assumed a small-work network structure. The co-expression interaction network consisted of 95 nodes and 132 edges, had a small-world index of 6.06, a diameter of 13, and average shortest path length of 5.12. The clustering coefficient was 0.102, which is higher than the clustering coefficient of a random graph with the same number of indices (0.031). The physical protein-protein interaction network consisted of 85 nodes and 41 edges, had a small-world index of 4.43, a diameter of 5, and average shortest path length of 82.03. The clustering coefficient was 0.326, which is also higher than the clustering coefficient of a random network with the same number of indices (0.06). Lastly, the genetic interaction network consists of 98 nodes and 472 edges, had a low small-world network index of 1.11, a diameter of 5, and average shortest path length of 2.33. The clustering coefficient for this network (0.197) was similar to the clustering coefficient of a random network with the same number of indices (0.192). Of the three networks, the co-expression network showed robust small-world network properties - the physical protein-protein interactions network had a substantially large shortest path length and the genetic-interactions network clustering coefficient did not differ from random. Therefore, in subsequent network analysis we focused on the co-expression network.

To further assess the structure of the co-expression network, we evaluated various network centrality measures, including degree centrality, eigenvector centrality, and edge-betweenness centrality. Centrality network measures define how important each node is within a given network. The degree centrality is the number of edges connected to a node. Eigenvector centrality is the extent to which a node is connected to other highly influential “hub” nodes. Edge-betweenness centrality is the extent to which a node lies on the shortest path between other nodes (see Methods). Fig. 4a shows the co-expression network. The size of each node is defined by its eigenvector centrality (EC) value. Genes with a high EC and high degree centrality can be characterized as hubs^15,16^. Within the co-expression network, genes with high degree centrality (Supplementary Fig. 3) and high EC (Supplementary Fig. 4) were *GIPC1* (n=8, EC=1.00), *ELMO1* (n=6, EC = 0.994), *UGCG* (n=7, EC = 0.978), *SMARCA2* (n=6, EC = 0.924), *ATXN2* (n=6, EC = 0.765), and *SETD2* (n=6, EC = 0.729). In addition to these, *DOCK2* (n= 5, EC = 0.809), *KIAA0917*/*SCFD1* (n=5, EC = 0.752), *HTR2A* (n=4, EC = 0.695), and *COL16A1* (n=5, EC = 0.645) were also highly influential based on EC values.

**Fig. 4:**
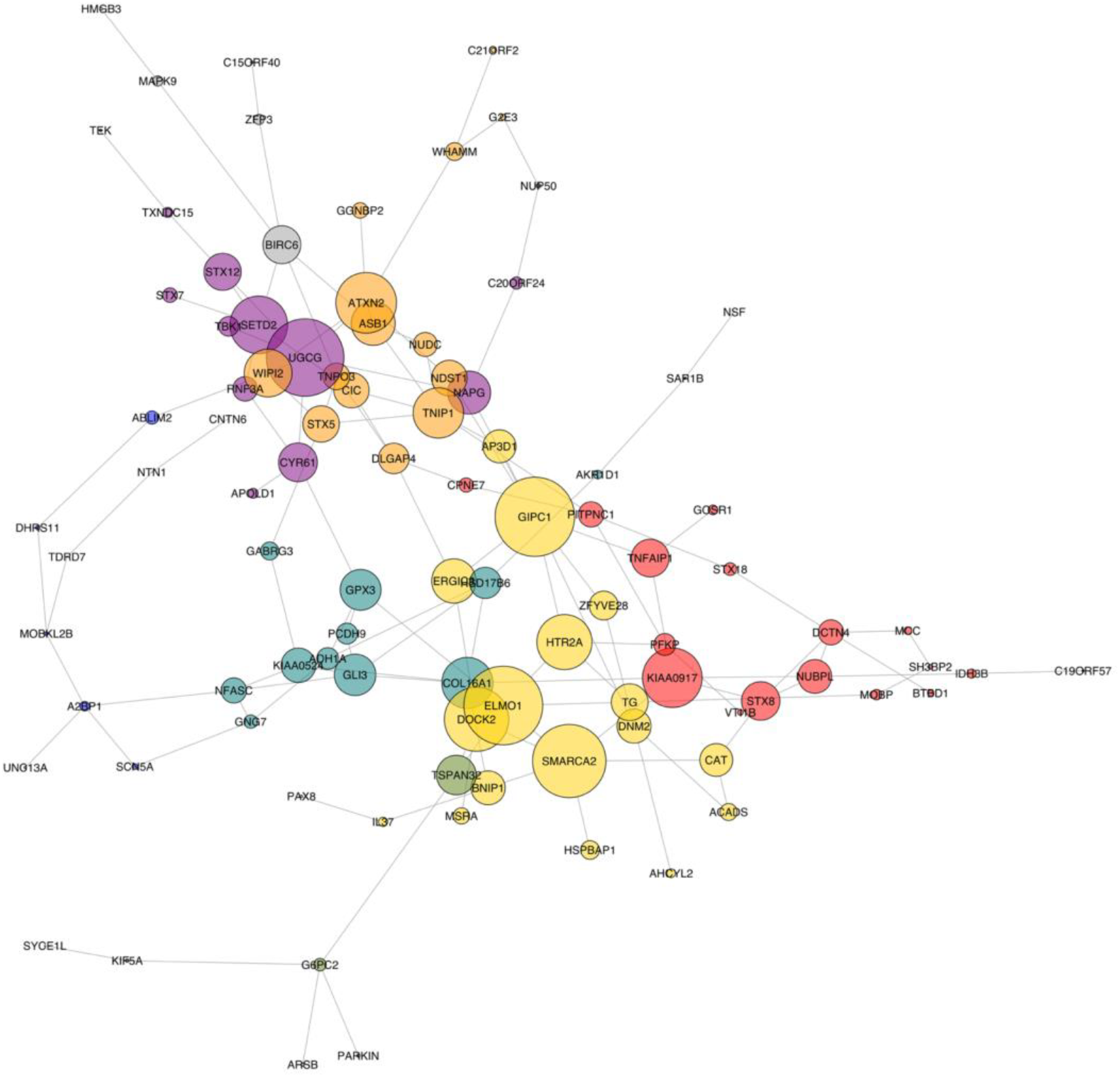
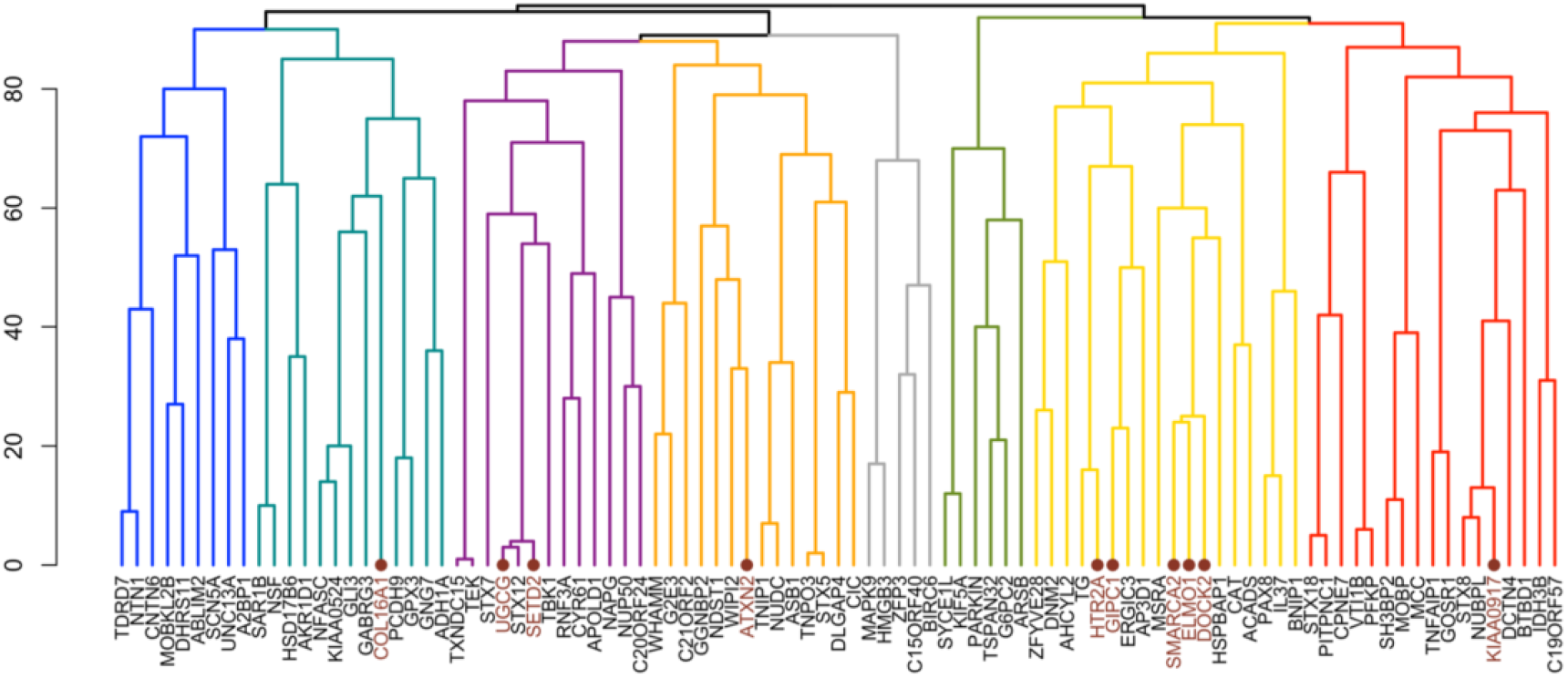
Co-expression interaction network plot and dendrogram. **a,** Co-expression interaction network plot. Each node represents a single gene and the edges (lines between genes) represent co-expression interactions between genes. The size of each node corresponds to the eigenvector centrality score for that gene. The color of each node represents membership to a distinct subnetwork. **b,** Co-expression interaction dendrogram plot. The co-expression network was partitioned into 8 subnetworks by removing edges with high edge-betweenness centrality. Hub genes are annotated in brown and marked by a large point.

Next, we explored whether the co-expression network could be partitioned into potentially meaningful subnetworks (Fig. 4a,b). The network was partitioned into subnetworks by removing edges with high edge-betweenness centrality (see Methods). The result was a hierarchical map, called a dendrogram (Fig. 4b). We identified a total of 8 clusters. Five clusters contained the most influential hub genes. The yellow cluster contained 5 hub genes (*HTR2A, GIPC1, SMARCA2, ELMO1*, and *DOCK2*). The purple cluster contained two hub genes (*UGCG* and *SETD2*), and the teal, orange, and red clusters contained a single hub gene (*COL16A1, ATXN2, KIAA0917*, respectively).

Collectively, these findings suggest that ALS genes that are co-expressed form networks with small-world properties and can be further partitioned into different clusters. Importantly, specific genes, such as *GIPC1* and *ATXN2*, act like hubs, characterized by a high degree of connectivity with other genes; perturbing these hub genes may disrupt the entire network.

### Biological pathways associated with ALS risk genes within co-expression network

We used bioinformatics approaches to identify distinct and common biological pathways associated with the 8 subnetworks within the ALS co-expression network (see Methods). The top 5 functional annotations for each subnetwork are shown in Fig. 5a. The blue subnetwork was enriched for developmental processes, including neuron projection development and regulation of cell growth. The gray subnetwork was enriched for apoptotic signaling pathway, cell death, and protein phosphorylation. The green subnetwork was enriched for cell-cell signaling, response to external stimuli, and metabolic processes. The purple cluster was enriched for vascular morphology and angiogenesis. The orange, red, teal, and yellow subnetworks were enriched for vesicle-mediated processes. Lastly, the orange and red networks were also enriched for organelle membrane fusion and disassembly processes, and the teal subnetwork was also enriched for oxidative reduction and androgen metabolic processes.

**Fig. 5:**
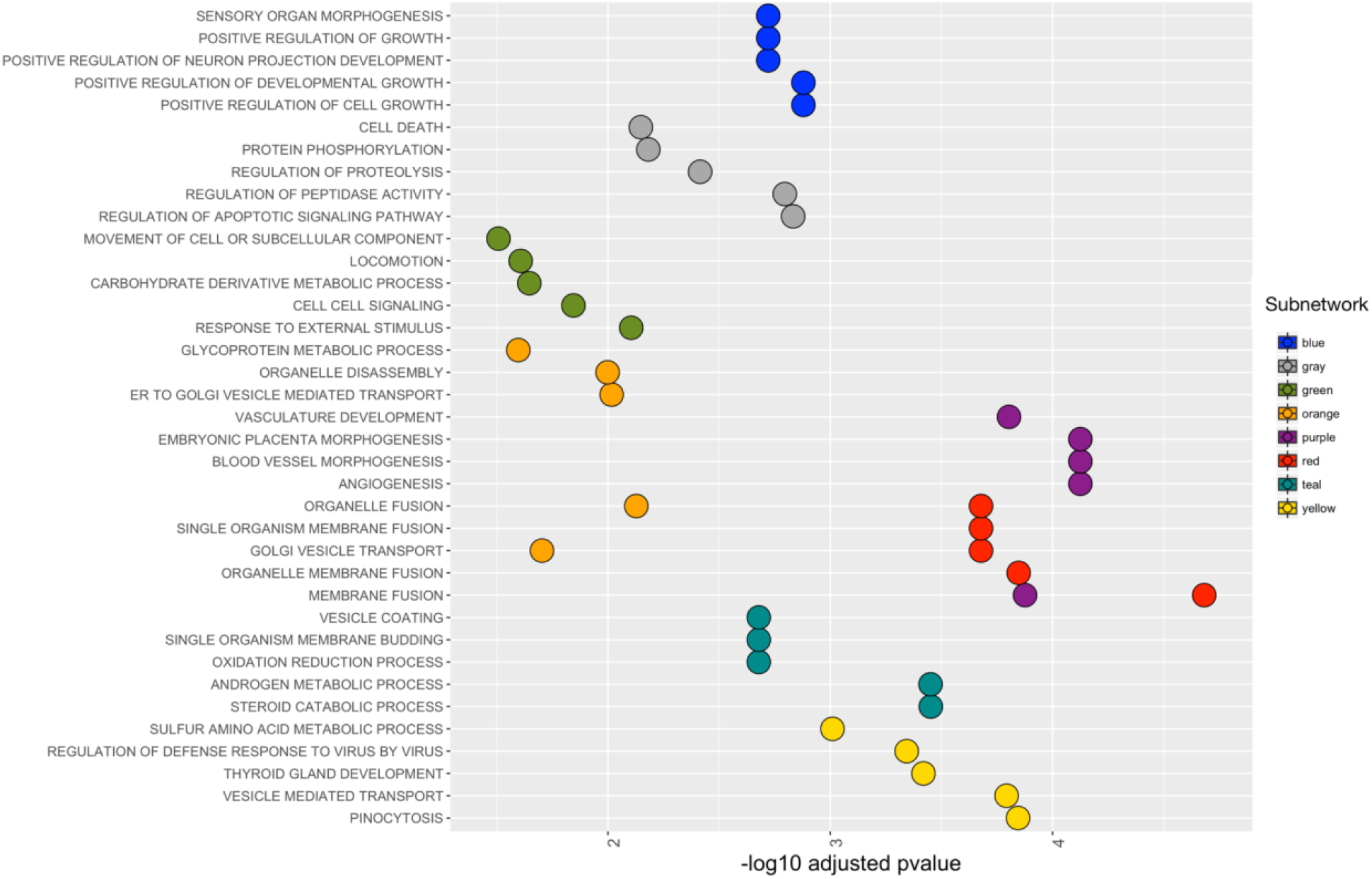
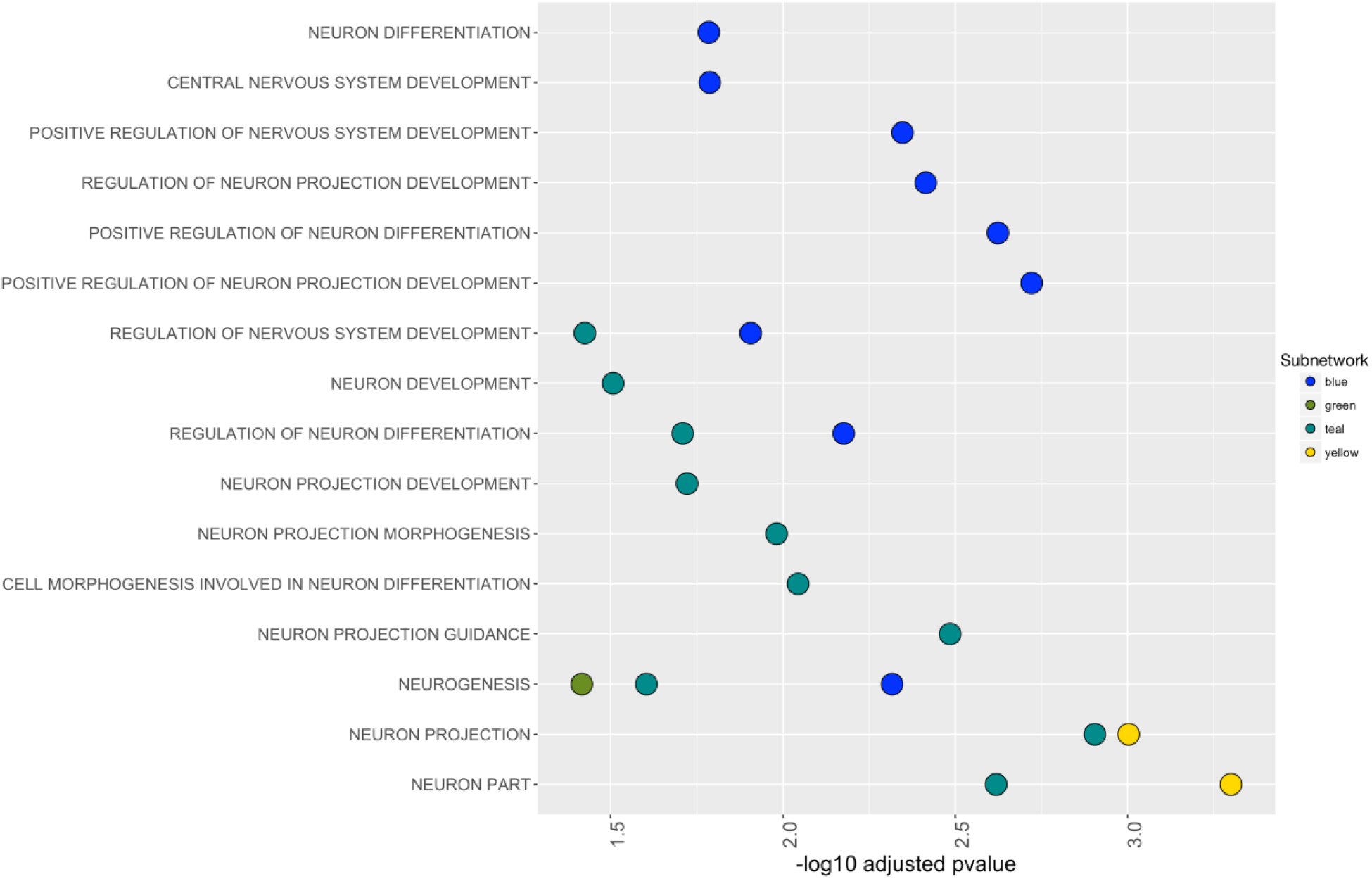
Biological pathways associated with the ALS co-expression subnetworks. **a,** Biological pathways associated with each subnetwork of the ALS co-expression network classified using FUMA (http://fuma.ctglab.nl/). **b,** Neuron-specific biological pathways associated with each subnetwork also classified using FUMA.

Further, we explored whether genes within the distinct subnetworks were selectively enriched for neuron-specific biological, chemical, or molecular processes. As shown in Fig. 5b, the blue, green, teal, and yellow subnetworks were selectively enriched for neuronal processes, particularly involved in neuron differentiation, projection, guidance, and development. We also found that several hub genes within the yellow subnetwork (*GIPC1, SMARC2, HTR2A*) were over-represented in these processes (Supplementary Fig. 5). Taken together, these findings highlight that the ALS co-expression subnetworks are involved in distinct and overlapping biological pathways.

### Differential expression of ALS risk genes in tissue from ALS patients and SOD1 G93A transgenic mice

We investigated whether particular clusters or hub genes were enriched in pathological samples from ALS patients. To do this, we assessed the differential expression of genes within the ALS co-expression network in the gray matter of motor neurons isolated from spinal cords of patients with familial and sporadic ALS and controls (Fig. 6a, see Methods). We found that 2 genes within the teal subnetwork (*COL16A1* and *GPX3)* and one gene within the purple subnetwork *(UGCG)* were differentially expressed in tissue from controls compared to familial ALS patients. *COL16A1* was also differentially expressed in tissue from controls compared to sporadic ALS patients. Importantly, *COL16A1* and *UGCG* are hub genes.

**Fig. 6:**
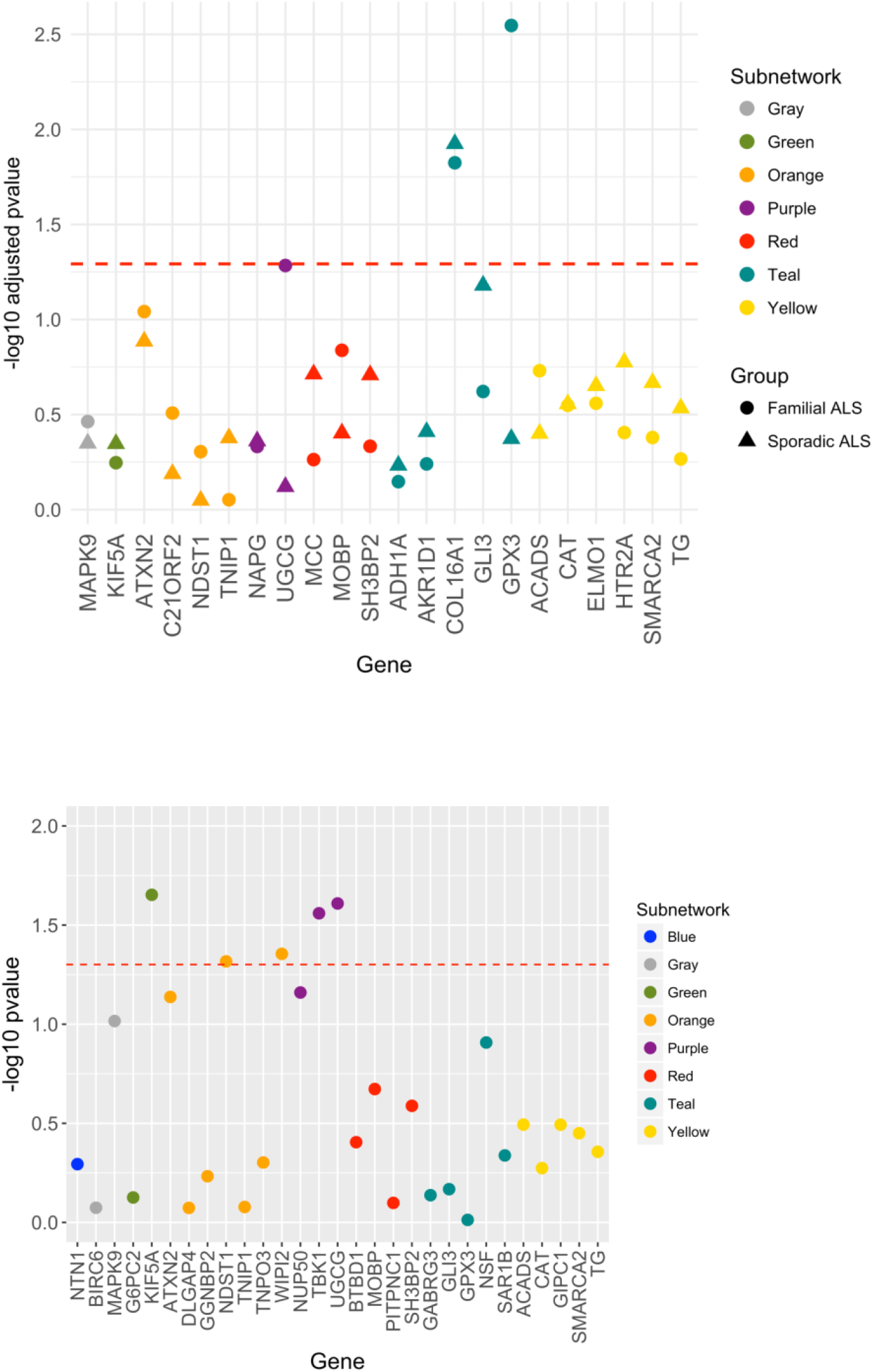
Differential expression of ALS risk genes in diseased tissue. **a,** Differential expression of ALS risk genes in tissues of patients with ALS. **b,** Differential expression of ALS risk genes in SOD1 G93A transgenic mouse.

To validate the genes identified in ALS human tissue, we evaluated expression data from a well-characterized mouse model^17^. The RNA expression data were analyzed in non-transgenic SOD1 WT and SOD1 G93A mice at 75 and 110 days (GEO accession number GSE4390). Differential expression of *UGCG* was independently replicated in the SOD1 G93A mice (Fig. 6b). Additionally, we found that *TBK1* within the purple cluster, *NDSTI* and *WIPI2* within the orange cluster, and *KIF5A* within the green cluster were enriched in SOD1 G93A mice (Fig. 6b).

## Discussion

We sought to elucidate the genetic basis of sporadic ALS. By exploiting statistical power from several large GWAS of >3 million people, we identified novel susceptibility loci, each associated with a small increase in ALS risk. We found that ALS variants form a small-world co-expression network characterized by highly inter-connected ‘hub’ genes. This network clustered into smaller sub-networks, each with a unique function. Altered gene expression of several sub-networks and hubs was over-represented in neuropathological samples from ALS patients and SOD1 G93A mice. Our collective findings indicate that the genetic architecture of ALS can be partitioned into distinct components where some genes are highly important for developing disease.

We found that ALS has a robust polygenic component. By leveraging genetic studies from 65 different traits and diseases, we identified 89 ALS risk loci across 21 chromosomes of which 59 are novel. Beyond *C9ORF72*, our pleiotropy analyses detected novel genetic signal within numerous loci including *GIPC1, ELMO1* and *COL16A* and confirmed previously reported variants, such as *ATXN2, KIF5A, UNC13A* and *MOBP* ^4–6^. Neither as polygenic as schizophrenia^18^ or Alzheimer’s disease^19^ nor purely oligogenic^5^, it is likely that the genetic architecture of ALS is a continuum of common low-risk variants and rare high-risk variants. Although each of the ALS susceptibility loci we detected was associated with a small effect, when aggregated together into a polygenic score they may explain a substantial portion of the inherited risk underlying ALS^20^.

We found genetic pleiotropy between ALS and a number of diseases and traits. In line with previous reports, we show strong genetic enrichment in ALS conditional on FTD and PSP^8,21,22^. Building on prior work showing a relationship between inflammation/immune dysfunction and motor neuron disease^23–25^, we found enrichment in ALS SNPs as a function of CRP and celiac disease. Surprisingly, we also identified strong genetic overlap between ALS and CAD, and memory function. Clinically, these findings suggest that a subset of ALS patients are at elevated genetic risk for FTD whereas another (potentially overlapping) group of ALS individuals may be at high risk for CAD or immune dysfunction. Therefore, development of multiple pathway specific polygenic scores may identify individuals at risk for developing ALS who are ‘enriched’ for FTD, cardiovascular or immune mediated processes.

Our findings inform cohort stratification and enrichment strategies for ALS clinical trials. We found that the ALS pleiotropic and functional risk genes form a small-world co-expression network. This network can be partitioned into 8 subnetworks, each enriched for distinct biological pathways. Similar to previous research, we found functional enrichment of ALS risk genes for oxidative-mediated (teal subnetwork), neuronal (teal, blue, yellow, and green subnetworks), and endoplasmic reticulum processes (orange subnetwork)^26,27^. Additionally, genes within the blue subnetwork were enriched for developmental and growth pathways while genes within the gray subnetwork were enriched for cell-death and apoptotic processes. Altered gene expression within the teal subnetwork was over-represented in postmortem spinal cord samples from familial and sporadic ALS patients, whereas abnormal expression of genes in the orange and purple clusters was present in SOD1 G93A mice. Clinically, these results suggest that partitioning genetic susceptibility may help identify individuals who have a higher likelihood of responding to therapies with a specific mechanism of action and support including DNA collection and sequencing in ALS clinical trials. For example, ALS patients who are enriched for genetic abnormalities within the teal subnetwork may respond to therapies targeting oxidation reduction or steroid catabolism (Fig. 5). On the other hand, vascular treatments may be effective in ALS individuals with an overabundance of altered purple cluster genes (Fig. 5).

Not all ALS genes are created equal. We show that particular genes within the ALS co-expression network are characteristic of hubs. Playing a central role within a biological system, perturbation of a hub can cause rapid degeneration of the whole network^28^. We found that these hub genes were key drivers of biological enrichment. Specifically, *HTR2A, SMARCA2*, and *GIPC1* were enriched for neuronal processes (Supplementary Fig. 5), and *ELMO1* and *DOCK2* were enriched for vesicle-mediated transport (Supplementary Fig. 6). Therapeutically targeting hub genes may be most effective for altering an entire biological pathway. However, given abundant pleiotropy with other traits (Supplementary Fig. 2), comprehensive biological and experimental evaluation of the entire network of a hub gene will be necessary prior to therapeutic evaluation.

This study should be interpreted within the context of its limitations. First, the ALS GWAS used contained people predominantly of European descent, while the other GWAS included people of both European and non-European descent. Therefore, these results may not be generalizable to ALS patients from other populations. Second, like most GWAS, a major limitation of our study is that we could not determine with certainty the causal genes underlying our genetic signal. Although we performed extensive LD and *cis-*eQTL analyses and included the combination of closest and functional genes in our network analyses, it is likely that genetic fine mapping and experimental approaches, such as CRISPR/Cas9 gene editing, will be needed to isolate the causal variants. Finally, given evidence that a substantial proportion of coronary disease is associated with inflammation^29^, future work should evaluate whether CAD influences ALS risk through inflammation or other mediator variables.

In summary, we show that the genetic architecture of ALS has a robust polygenic component that can be partitioned into distinct subnetworks, each enriched for divergent biological pathways. We also identify several hub genes that may be key drivers of ALS pathobiology. Our findings are compatible with the hypothesis that ALS is a multi-step, non-uniform disease process. These results have implications for cohort stratification and enrichment strategies for ALS clinical trials.

## Methods

### Participant samples

We conducted a meta-analysis of summary data obtained from published data. We evaluated complete GWAS results in the form of summary statistics (p-values and odds ratios) for ALS and 65 distinct traits and diseases (see Supplementary Table 4). We obtained ALS GWAS summary statistic data from 12,577 ALS cases and 23,475 controls at 18,741,501 SNPs (see Supplementary Table 4 for additional details). The ALS GWAS summary statistics and sequenced variants are publicly available through the Project MinE data browser: http://databrowser.projectmine.com. We also obtained GWAS summary statistic data for the 65 distinct traits and diseases (for additional details, please see Supplementary Table 4). The relevant institutional review boards or ethics committees approved the research protocol of the individual GWASs used in the current analysis, and all participants gave written informed consent.

### Genetic Enrichment Statistical Analyses

The pleiotropic enrichment strategies implemented here were derived from previously published stratified FDR methods^13,30^. For given phenotypes A and B, pleiotropic ‘enrichment’ of phenotype A with phenotype B exists if the proportion of SNPs or genes associated with phenotype A increases as a function of increased association with phenotype B. To assess for enrichment, we constructed fold-enrichment plots of nominal –log_10_(p) values for all ALS SNPs and for subsets of SNPs determined by the significance of their association with the 65 distinct traits and diseases. In fold-enrichment plots, the presence of enrichment is reflected as an upward deflection of the curve for phenotype A with increasing strength of association with phenotype B. To assess for polygenic effects below the standard GWAS significance threshold, we focused the fold-enrichment plots on SNPs with nominal –log_10_(p) < 7.3 (corresponding to p > 5×10^−8^). The enrichment seen can be directly interpreted in terms of true discovery rate (TDR = 1 – False Discovery Rate (FDR)). Given prior evidence that several genetic variants within chromosome 9 are associated with increased ALS risk, one concern is that random pruning may not sufficiently account for these large LD blocks, resulting in artificially inflated genetic enrichment^12^. To better account for these large LD blocks, in our genetic enrichment analyses, we removed all SNPs within chromosome 9.

To identify novel ALS risk loci as a function of genetic variants associated with the 65 traits and diseases, we computed conditional FDRs^13,30^, a statistical framework that is well suited for gene detection. The standard FDR framework is based on Bayesian statistics and follows the assumption that SNPs are either associated with the phenotype (non-null) or are not associated with the phenotype (null SNPs). Within a Bayesian statistical framework, the FDR is then the posterior probability of the SNP being null given its p-value is as small as or smaller than the observed one. The conditional FDR is an extension of the standard FDR, which incorporates information from GWAS summary statistics of a second phenotype to adjust its significance level. The conditional FDR is defined as the probability that a SNP is null in the first phenotype given that the p-values in the first and second phenotypes are as small as or smaller than the observed ones. Ranking SNPs by the standard FDR or by p-values gives the same ordering of SNPs. In contrast, if the primary and secondary phenotypes are related genetically, the conditional FDR reorders SNPs and results in a different ranking than that based on p-values alone. We used an overall FDR threshold of p < .05 to indicate statistical significance, meaning 5 expected false discoveries per 100 reported. In addition, we constructed Manhattan plots based on the ranking of the conditional FDR to illustrate the genomic location. In all analyses, we controlled for the effects of genomic inflation. Detailed information on the conditional FDR can be found in prior reports^13,30^.

### Functional evaluation of shared risk loci

To assess whether the SNPs associated with ALS and the 65 traits and diseases modify gene expression, we identified *cis*-expression quantitative loci (eQTLs, defined as variants within 1 Mb of a gene’s transcription start site) associated with the identified ALS pleiotropic SNPs and measured their regional brain expression in a publicly available dataset of normal control brains (UK Brain Expression Consortium, http://braineac.org/)^31^. To minimize multiple comparisons, we analyzed *cis-*eQTL for the mean p-value obtained from the following brain regions: the cerebellum, frontal cortex, hippocampus, medulla, occipital cortex, putamen, substantia nigra, temporal cortex, thalamus, and white matter. To minimize false positives, we applied a Bonferroni-corrected p-value of 1.5 x 10^−3^.

### Biological networks associated with ALS genetic risk genes

To evaluate potential protein and genetic interactions, co-expression, co-localization, and protein domain similarity for the combined pleiotropic (i.e. closest genes from the pleiotropy analyses) and functionally expressed ALS genes (i.e., with significant *cis*-eQTLs), we used GeneMANIA (www.genemania.org), an online web-portal for bioinformatic assessment of gene networks^14^. To visualize the composite gene network, we also assessed the weights of individual components within the network^32^. Further, we evaluated whether the biological networks fell into the class of a small-world network using several diagnostic criteria. First, we computed the “small-worldness” index, using the R package ‘qgraph’. The function computes the global transitivity of the target network and its average shortest path length^16,33^. It then computes the same indices on 1000 random networks. The small-worldness index is then equal to the transitivity of the target network (normalized by the random transitivity) over the average shortest path of the target network (normalized by the random average shortest path length). A network was considered small-world if the “small-worldness” index was >= 3; ^16^. In addition to the small-worldness index, we inspected whether the network had a transitivity substantially higher than comparable random networks and that its average shortest path length was similar or higher (but not substantially higher) than that computed on random networks.

Further, to determine whether some genes play a more influential role than others, we evaluated various network centrality measures, including degree centrality, eigenvector centrality, and edge-betweenness centrality. We used the R package ‘igraph’ for all network centrality analysis and visualization^34^. The degree of a node corresponds to the sum of its adjacent edges (i.e., connections). The eigenvector centrality of a node corresponds to the values of the first eigenvector of the graph adjacency matrix. In general, nodes with high eigenvector centralities are also connected to many other nodes which are, in turn, connected to many other nodes. Consequently, eigenvector centrality corresponds to the degree to which a node is connected to other highly influential nodes. Lastly, we partitioned the co-expression network, in particular, into subnetworks based on edge-betweenness centrality. Edge-betweenness centrality is defined by the number of shortest-paths going through an edge. Here, a subnetwork is analogous to modules within a network. Nodes within a module are densely connected to themselves (e.g., cluster) but sparsely connected to other modules. To create modules, we gradually remove the edge with the highest edge-betweenness score, since all the shortest paths from one module to another typically pass through them. The result is a hierarchical map, called a dendrogram. The leaves of the tree are the individual nodes and the root of the tree represents the whole graph.

Lastly, to evaluate biological pathways of the ALS pleiotropic genes (i.e. closest genes from the pleiotropy analyses) and functionally expressed ALS genes (i.e., with significant *cis*-eQTLs), we used FUMA (http://fuma.ctglab.nl/), a web-based platform that integrates information from multiple biological resources to facilitate functional annotation of GWAS results^35^.

### Gene expression alterations in tissue from ALS patients and SOD1 G93A transgenic mice

To determine whether the ALS genetic risk genes were differentially expressed in tissue from patients with ALS, we analyzed the gene expression of the target genes from postmortem spinal cord gray matter from 11 individuals (2 patients with familial ALS, 5 patients with sporadic ALS, and 4 controls; Gene Expression Omnibus [GEO] accession number GDS412^36^). Details about this dataset and analysis–including the human brain samples used, RNA extraction and hybridization methods, microarray quality control, and microarray data analysis– are described in the original manuscript^36^. To validate the genes identified in ALS human tissue, we also analyzed RNA expression data in non-transgenic SOD1 WT (n=2) and SOD1 G93A (n=2) mice at 75 and 110 days (GEO accession number GSE4390). SOD1 G93A mice are pre-symptomatic at 75 days and exhibit hindlimb paralysis at 110 days^17,37^.

### Code availability

Code and scripts available by request from authors.

### Data availability

Summary statistics from secondary GWAS of single disorders and traits are available upon request from the corresponding author. Cis-eqtl data from the UK Brain Expression Consortium are publicly available (http://braineac.org/). Findings from biological networks were obtained using GeneMANIA (www.genemania.org), an online web-portal for bioinformatic assessment of gene networks. Biological pathways were evaluated using FUMA (http://fuma.ctglab.nl/), a web-based platform that integrates information from multiple biological resources to facilitate functional annotation of GWAS results. Expression data from sporadic and familial ALS patients and controls postmortem spinal cord gray matter are available in GEO with the accession number GDS412. RNA expression data in non-transgenic SOD1 WT and SOD1 G93A mice at 75 and 110 days are also available in GEO with the accession number GSE4390.

## ACKNOWLEDGEMENTS

We salute the millions of people who battle with ALS every single day - your courage is our strength. This research is in part an EU Joint Programme - Neurodegenerative Disease Research (JPND) project. The project is supported through the following funding organizations under the aegis of NL (ZONMW) and JPND - www.jpnd.eu (United Kingdom, Medical Research Council (MR/L501529/1 (STRENGTH); MR/R024804/1 (BRAIN-MEND)) and Economic and Social Research Council ((ES/L008238/1) ALS-CarE) and through the Motor Neurone Disease Association. This study represents independent research part funded by the National Institute for Health Research (NIHR) Biomedical Research Centre at South London and Maudsley NHS Foundation Trust and King’s College London. This project has also received funding from the European Research Council (ERC) under the European Union’s Horizon 2020 research and innovation programme (grant agreement n° 772376 – EScORIAL).

