## Supplemental for "Partitioning the genetic architecture of amyotrophic lateral sclerosis" (from another version of this preprint: version 1, 2018-12-23)

**Figure S1.** LD plots for SNPs within **A-F)** chromosomes 2-7, **G-O)** chromosomes 8, 9, 11-17.

**Figure S2.** Degree centrality scores for genes within the ALS co-expression network.

**Figure S3.** Eigenvector centrality scores for genes within the ALS co-expression network.

**Figure S4.** Genes enriched for biological pathways for each subnetwork.

**Figure S5.** Genes enriched for neuron-specific biological pathways.

Figure S1

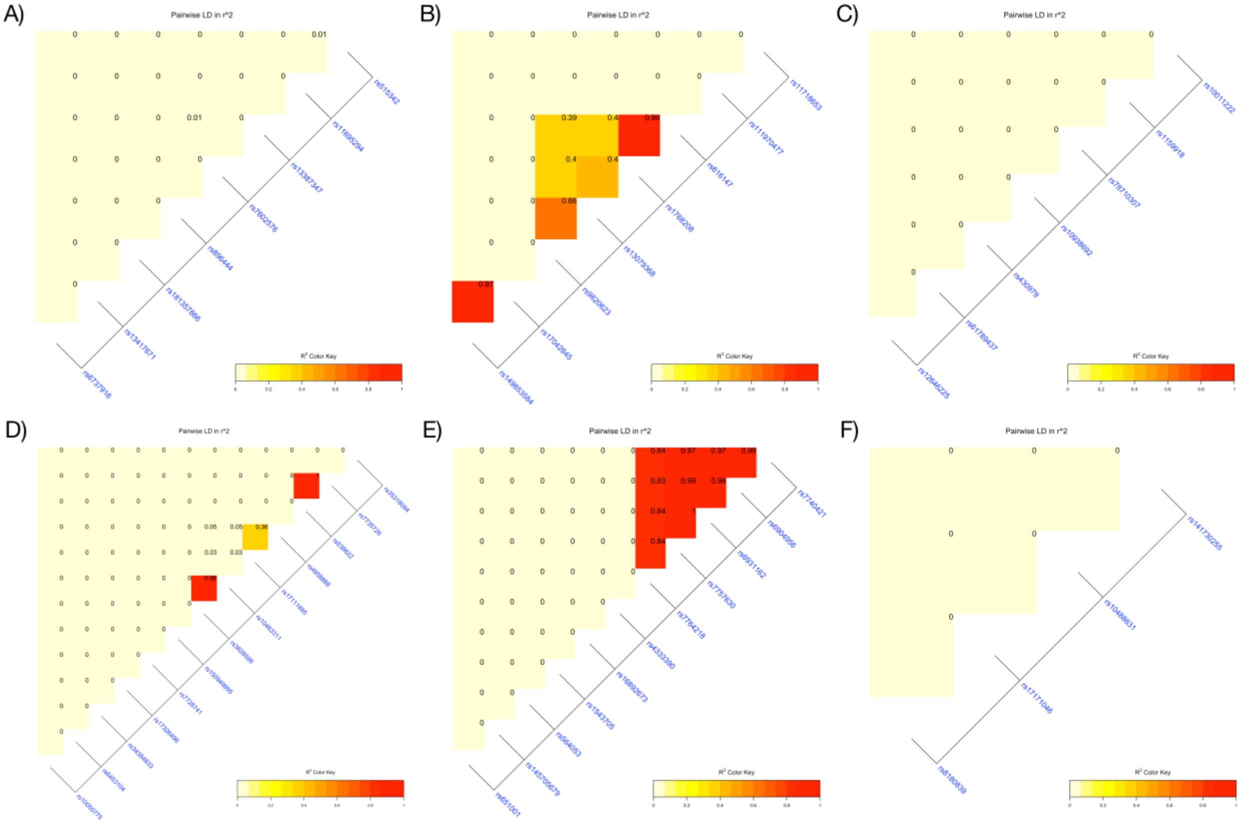

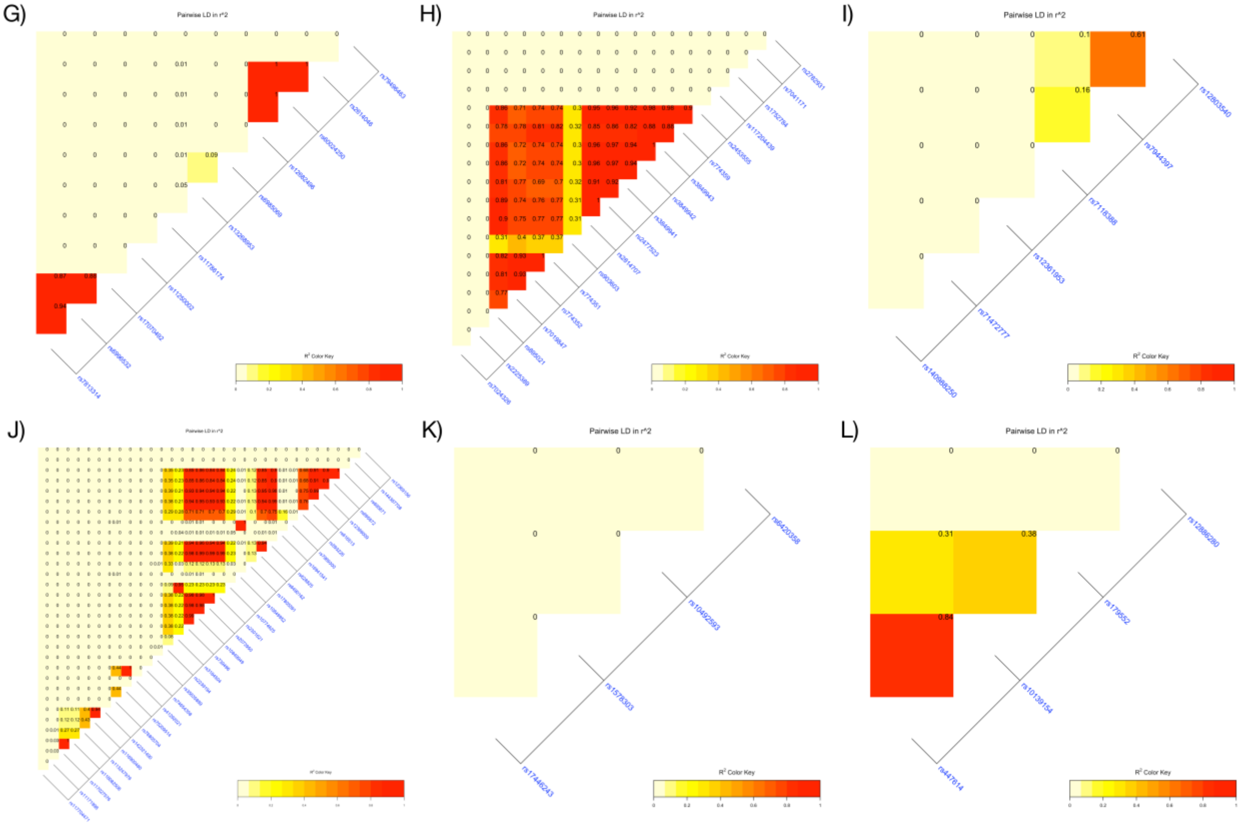
G)
Pairwise LD in r^2
H)
Pairwise LD in r^2
I)
Pairwise LD in r^2
J)
K)
Pairwise LD in r^2
L)
Pairwise LD in r^2
R² Color Key

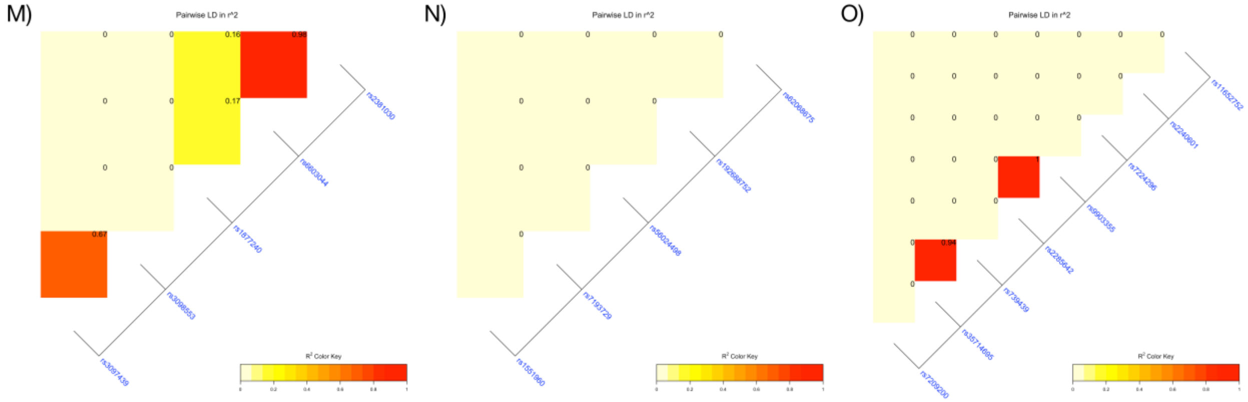
M)
Pairwise LD in r^2
0
0
0.16
0.98
0.17
0.67
rs2381030
rs6603044
rs1877240
rs3098553
rs3097439
R2 Color Key
N)
Pairwise LD in r^2
rs62068875
rs192688752
rs56024498
rs7193729
rs1551960
R2 Color Key
O)
Pairwise LD in r^2
1
0.94
rs11652752
rs2240601
rs7224296
rs9903355
rs2285642
rs739439
rs35714695
rs7209200
R2 Color Key

Figure S2

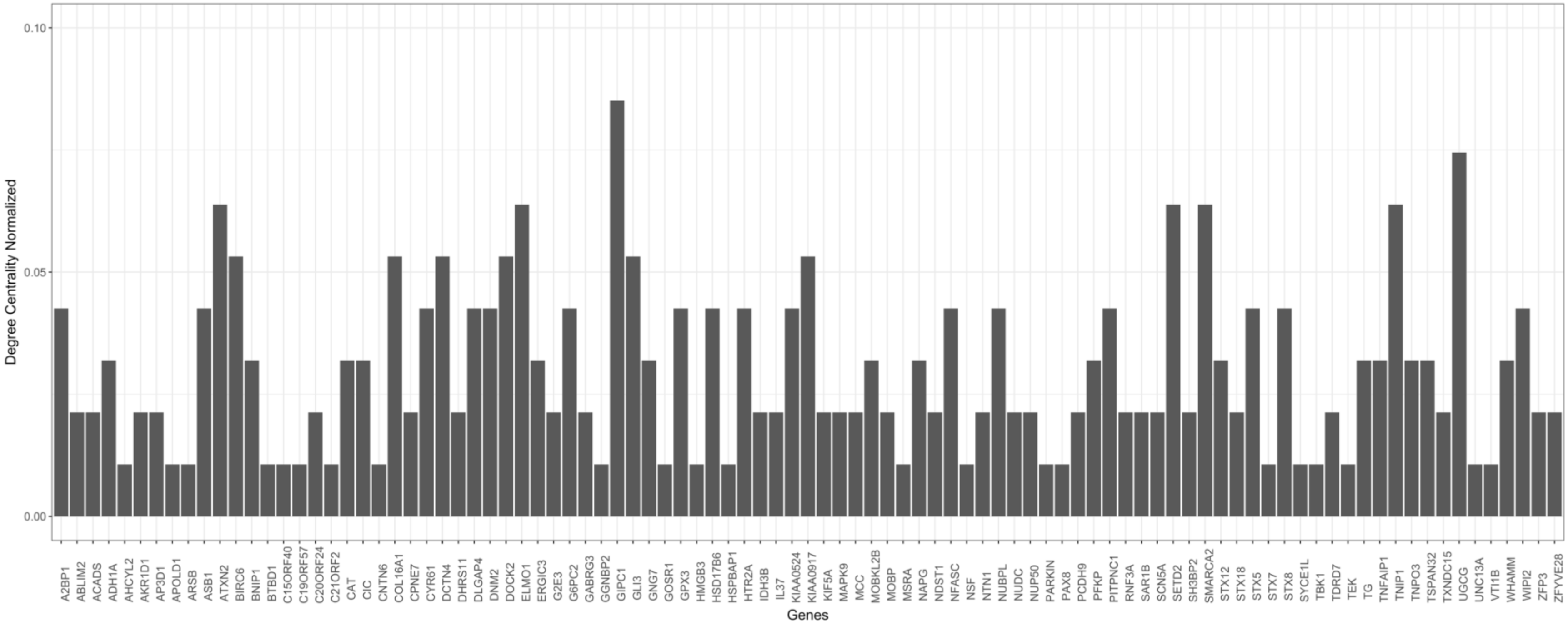
Degree Centrality Normalized
0.10
0.05
0.00
Genes
A2BP1
ABLIM2
ACADS
ADH1A
AHCYL2
AKR1D1
AP3D1
APOLD1
ARSB
ASB1
ATXN2
BIRC6
BNIP1
BTBD1
C15ORF40
C19ORF57
C20ORF24
C21ORF2
CAT
CIC
CNTN6
COL16A1
CPNE7
CYR61
DCTN4
DHRS11
DLGAP4
DNM2
DOCK2
ELMO1
ERGIC3
G2E3
G6PC2
GABRG3
GGNBP2
GIPC1
GLI3
GNG7
GOSR1
GPX3
HMGB3
HSD17B6
HSPBAP1
HTR2A
IDH3B
IL37
KIAA0524
KIAA0917
KIF5A
MAPK9
MCC
MOBKL2B
MOBP
MSRA
NAPG
NDST1
NFASC
NSF
NTN1
NUBPL
NUDC
NUP50
PARKIN
PAX8
PCDH9
PFKP
PITPNC1
RNF3A
SAR1B
SCN5A
SETD2
SH3BP2
SMARCA2
STX12
STX18
STX5
STX7
STX8
SYCE1L
TBK1
TDRD7
TEK
TG
TNFAIP1
TNIP1
TNPO3
TSPAN32
TXNDC15
UGCG
UNC13A
VTI1B
WHAMM
WIPI2
ZFP3
ZFYVE28

Figure S3

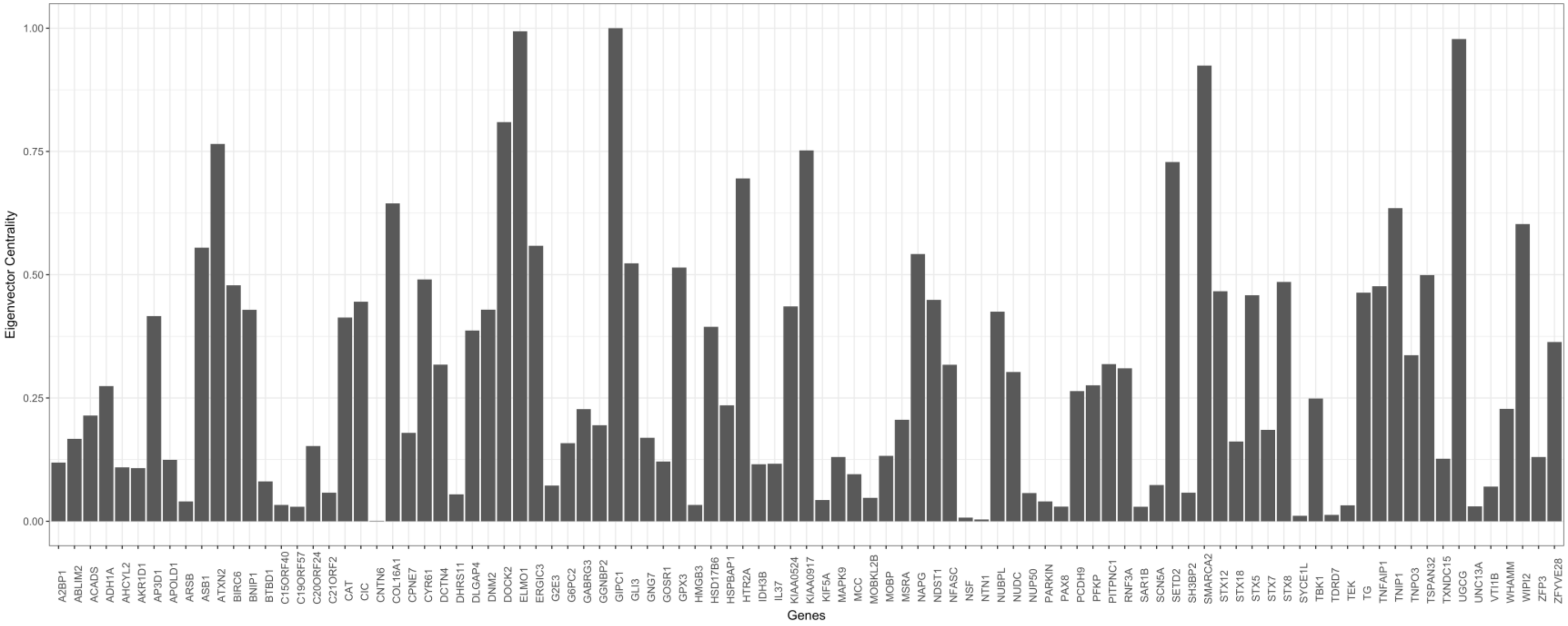

Eigenvector Centrality
1.00
0.75
0.50
0.25
0.00
Genes
A2BP1
ABLIM2
ACADS
ADH1A
AHCYL2
AKR1D1
AP3D1
APOLD1
ARSB
ASB1
ATXN2
BIRC6
BNIP1
BTBD1
C15ORF40
C19ORF57
C20ORF24
C21ORF2
CAT
CIC
CNTN6
COL16A1
CPNE7
CYR61
DCTN4
DHRS11
DLGAP4
DNM2
DOCK2
ELMO1
ERGIC3
G2E3
G6PC2
GABRG3
GGNBP2
GIPC1
GLI3
GNG7
GOSR1
GPX3
HMGB3
HSD17B6
HSPBAP1
HTR2A
IDH3B
IL37
KIAA0524
KIAA0917
KIF5A
MAPK9
MCC
MOBKL2B
MOBP
MSRA
NAPG
NDST1
NFASC
NSF
NTN1
NUBPL
NUDC
NUP50
PARKIN
PAX8
PCDH9
PFKP
PITPNC1
RNF3A
SAR1B
SCN5A
SETD2
SH3BP2
SMARCA2
STX12
STX18
STX5
STX7
STX8
SYCE1L
TBK1
TDRD7
TEK
TG
TNFAIP1
TNIP1
TNPO3
TSPAN32
TXNDC15
UGCG
UNC13A
VTI1B
WHAMM
WIPI2
ZFP3
ZFYVE28

Figure S4

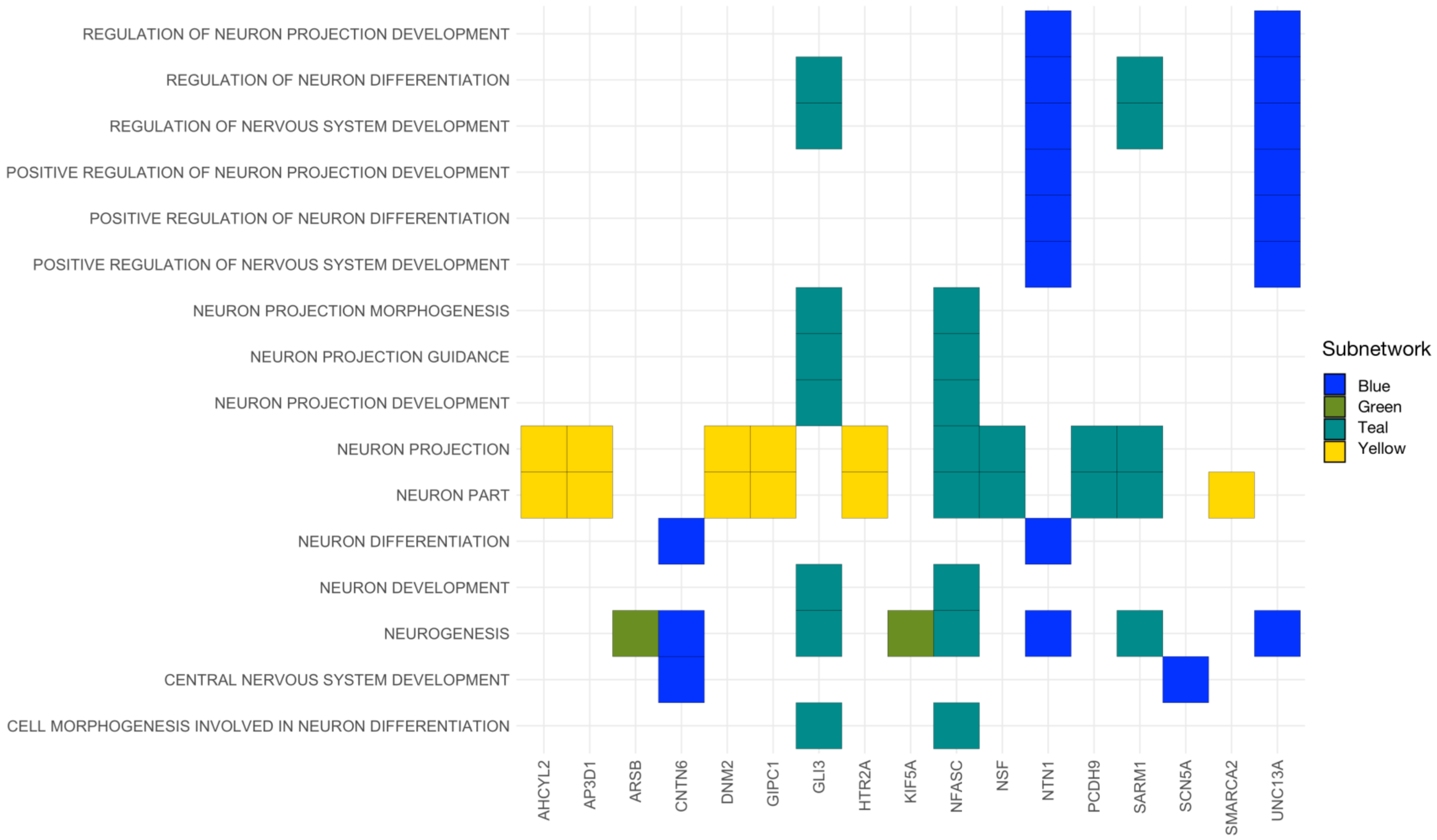

REGULATION OF NEURON PROJECTION DEVELOPMENT
REGULATION OF NEURON DIFFERENTIATION
REGULATION OF NERVOUS SYSTEM DEVELOPMENT
POSITIVE REGULATION OF NEURON PROJECTION DEVELOPMENT
POSITIVE REGULATION OF NEURON DIFFERENTIATION
POSITIVE REGULATION OF NERVOUS SYSTEM DEVELOPMENT
NEURON PROJECTION MORPHOGENESIS
NEURON PROJECTION GUIDANCE
NEURON PROJECTION DEVELOPMENT
NEURON PROJECTION
NEURON PART
NEURON DIFFERENTIATION
NEURON DEVELOPMENT
NEUROGENESIS
CENTRAL NERVOUS SYSTEM DEVELOPMENT
CELL MORPHOGENESIS INVOLVED IN NEURON DIFFERENTIATION
AHCYL2
AP3D1
ARSB
CNTN6
DNM2
GIPC1
GLI3
HTR2A
KIF5A
NFASC
NSF
NTN1
PCDH9
SARM1
SCN5A
SMARCA2
UNC13A
Subnetwork
Blue
Green
Teal
Yellow

Figure S5

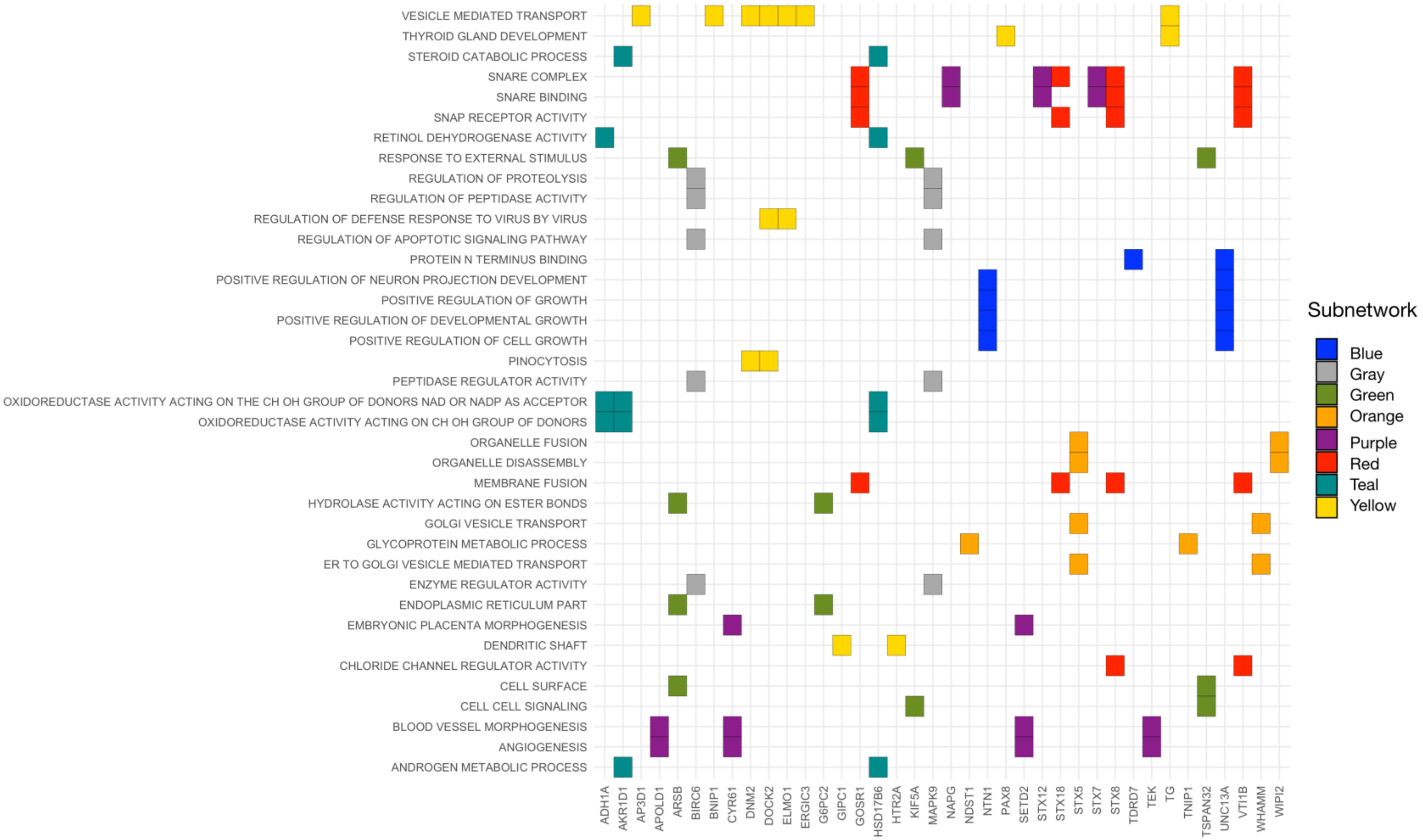
VESICLE MEDIATED TRANSPORT
THYROID GLAND DEVELOPMENT
STEROID CATABOLIC PROCESS
SNARE COMPLEX
SNARE BINDING
SNAP RECEPTOR ACTIVITY
RETINOL DEHYDROGENASE ACTIVITY
RESPONSE TO EXTERNAL STIMULUS
REGULATION OF PROTEOLYSIS
REGULATION OF PEPTIDASE ACTIVITY
REGULATION OF DEFENSE RESPONSE TO VIRUS BY VIRUS
REGULATION OF APOPTOTIC SIGNALING PATHWAY
PROTEIN N TERMINUS BINDING
POSITIVE REGULATION OF NEURON PROJECTION DEVELOPMENT
POSITIVE REGULATION OF GROWTH
POSITIVE REGULATION OF DEVELOPMENTAL GROWTH
POSITIVE REGULATION OF CELL GROWTH
PINOCYTOSIS
PEPTIDASE REGULATOR ACTIVITY
OXIDOREDUCTASE ACTIVITY ACTING ON THE CH OH GROUP OF DONORS NAD OR NADP AS ACCEPTOR
OXIDOREDUCTASE ACTIVITY ACTING ON CH OH GROUP OF DONORS
ORGANELLE FUSION
ORGANELLE DISASSEMBLY
MEMBRANE FUSION
HYDROLASE ACTIVITY ACTING ON ESTER BONDS
GOLGI VESICLE TRANSPORT
GLYCOPROTEIN METABOLIC PROCESS
ER TO GOLGI VESICLE MEDIATED TRANSPORT
ENZYME REGULATOR ACTIVITY
ENDOPLASMIC RETICULUM PART
EMBRYONIC PLACENTA MORPHOGENESIS
DENDRITIC SHAFT
CHLORIDE CHANNEL REGULATOR ACTIVITY
CELL SURFACE
CELL CELL SIGNALING
BLOOD VESSEL MORPHOGENESIS
ANGIOGENESIS
ANDROGEN METABOLIC PROCESS
ADH1A
AKR1D1
AP3D1
APOLD1
ARSB
BIRC6
BNIP1
CYR61
DNM2
DOCK2
ELMO1
ERGIC3
G6PC2
GIPC1
GOSR1
HSD17B6
HTR2A
KIF5A
MAPK9
NAPG
NDST1
NTN1
PAX8
SETD2
STX12
STX18
STX5
STX7
STX8
TDRD7
TEK
TG
TNIP1
TSPAN32
UNC13A
VTI1B
WHAMM
WIPI2
Subnetwork
Blue
Gray
Green
Orange
Purple
Red
Teal
Yellow

Table S1. All SNPs associated with ALS pleiotropic risk genes at a conditional FDR < 0.05

| | **Disease/Trait** | **SNP** | **Nearest Gene** | **Chr** | **Position** | **ALS p-value** | **Min CondFDR** |
|---|---|---|---|---|---|---|---|
| 1 | VNR | rs10914464 | COL16A1 | 1 | 32132319 | 1.26E-04 | 4.54E-02 |
| 2 | HV | rs10443173 | COL24A1 | 1 | 86533754 | 8.20E-06 | 8.11E-03 |
| 3 | PSP | rs2068667 | NFASC | 1 | 204917680 | 3.39E-04 | 3.16E-02 |
| 4 | FTD | rs515342 | ASB1 | 2 | 239367296 | 6.22E-04 | 3.71E-02 |
| 5 | ADHD | rs896444 | BC038566 | 2 | 106241808 | 1.32E-04 | 3.89E-02 |
| 6 | TC | rs6737916 | BIRC6 | 2 | 32597985 | 2.82E-05 | 2.81E-02 |
| 7 | NEUR | rs181357866 | CRIM1 | 2 | 35863487 | 3.10E-05 | 1.92E-02 |
| 8 | T2D | rs13387347 | G6PC2 | 2 | 169754846 | 2.95E-05 | 3.52E-02 |
| 9 | BMI | rs7602576 | IL37 | 2 | 113699617 | 6.79E-05 | 4.80E-02 |
| 10 | LUNG | rs11695294 | LOC375295 | 2 | 177473618 | 5.12E-05 | 1.62E-02 |
| 11 | SMOKEONSET | rs13417671 | TMP_LOCUS_39 | 2 | 34043426 | 1.04E-05 | 1.36E-02 |
| 12 | AD | rs28407220 | TMP_LOCUS_39 | 2 | 34041173 | 6.91E-06 | 1.77E-02 |
| 13 | EVERSMOKE | rs17042645 | CNTN6 | 3 | 1731848 | 1.26E-04 | 2.45E-02 |
| 14 | BMI | rs11718653 | HSPBAP1 | 3 | 122477858 | 9.41E-06 | 6.15E-03 |
| 15 | PSP | rs9820623 | MOBP | 3 | 39493858 | 1.64E-05 | 1.89E-03 |
| 16 | FTD | rs1464047 | MOBP | 3 | 39526874 | 3.96E-05 | 2.25E-03 |
| 17 | CBD | rs1768208 | MOBP | 3 | 39523003 | 3.90E-05 | 4.35E-03 |
| 18 | CAD | rs6765697 | MOBP | 3 | 39493239 | 1.60E-05 | 3.99E-02 |
| 19 | PSP | rs562545 | MOBP | 3 | 39536524 | 5.64E-04 | 4.85E-02 |
| 20 | T2D | rs111970477 | SETD2 | 3 | 47105581 | 6.55E-05 | 3.76E-02 |
| 21 | MDD | rs10938692 | ABLIM2 | 4 | 8118561 | 2.57E-05 | 4.21E-02 |
| 22 | AD | rs1159918 | ADH1A | 4 | 100243009 | 1.98E-05 | 7.39E-03 |
| 23 | ALC | rs78710307 | BC034799 | 4 | 59983437 | 1.07E-04 | 4.32E-02 |
| 24 | ICV | rs72630987 | BC034799 | 4 | 59508966 | 1.39E-04 | 4.44E-02 |
| 25 | VNR | rs12646225 | RNF3A | 4 | 696848 | 1.51E-04 | 4.91E-02 |

| | | | | | | | |
|---|---|---|---|---|---|---|---|
| 26 | BREAST | rs430979 | SH3BP2 | 4 | 2814698 | 2.41E-06 | 1.40E-03 |
| 27 | PUT | rs62637887 | SH3BP2 | 4 | 2828139 | 5.79E-06 | 4.48E-02 |
| 28 | VNR | rs10011222 | SLC10A7 | 4 | 147441065 | 1.00E-04 | 3.84E-02 |
| 29 | INSOMNIA | rs61789437 | ZFYVE28 | 4 | 2315630 | 2.08E-06 | 9.40E-03 |
| 30 | VITILIGO | rs4974650 | ZFYVE28 | 4 | 2311719 | 1.53E-06 | 1.48E-02 |
| 31 | SWB | rs762837 | ZFYVE28 | 4 | 2316480 | 1.17E-05 | 2.52E-02 |
| 32 | CBD | rs4974674 | ZFYVE28 | 4 | 2311361 | 1.19E-05 | 4.83E-02 |
| 33 | SCZ | rs34384833 | AK056485 | 5 | 91238155 | 1.65E-05 | 2.50E-02 |
| 34 | CRP | rs7735726 | ERGIC1 | 5 | 172347823 | 1.91E-05 | 7.56E-03 |
| 35 | FTD | rs538622 | ERGIC1 | 5 | 172347679 | 1.40E-05 | 3.66E-02 |
| 36 | CHRONOTYPE | rs10050775 | GDNF | 5 | 38008042 | 1.00E-05 | 3.61E-02 |
| 37 | MENOPAUSE | rs10059028 | GDNF | 5 | 38015610 | 4.17E-05 | 4.63E-02 |
| 38 | CeD | rs3828599 | GPX3 | 5 | 150401796 | 2.45E-06 | 5.85E-04 |
| 39 | MEMORY | rs4958873 | GPX3 | 5 | 150404096 | 2.49E-06 | 9.34E-04 |
| 40 | PROSTATE | rs4958872 | GPX3 | 5 | 150402334 | 1.46E-06 | 5.00E-03 |
| 41 | PSOR | rs8177427 | GPX3 | 5 | 150403011 | 2.79E-06 | 2.52E-02 |
| 42 | T2D | rs4958434 | GPX3 | 5 | 150402570 | 2.58E-06 | 2.77E-02 |
| 43 | RT | rs35318094 | MAPK9 | 5 | 179672984 | 1.07E-05 | 3.53E-02 |
| 44 | WHR | rs17326496 | MCC | 5 | 112676579 | 5.79E-05 | 1.70E-02 |
| 45 | VNR | rs10463311 | NAF1 | 5 | 150410835 | 7.85E-07 | 6.47E-03 |
| 46 | SBP | rs11747926 | NAF1 | 5 | 150414388 | 4.98E-06 | 1.30E-02 |
| 47 | FTD | rs17111695 | NAF1 | 5 | 150432446 | 2.03E-05 | 3.77E-02 |
| 48 | LUNG | rs150949995 | NDST1 | 5 | 149898499 | 4.13E-06 | 1.01E-02 |
| 49 | PD | rs7728741 | SAR1B | 5 | 133942492 | 2.55E-04 | 4.58E-02 |
| 50 | VITILIGO | rs4958888 | TNIP1 | 5 | 150472842 | 1.45E-05 | 9.80E-03 |
| 51 | UC | rs3762996 | TNIP1 | 5 | 150469626 | 2.13E-05 | 3.65E-02 |
| 52 | WHR | rs1543705 | AK127472 | 6 | 127197108 | 1.92E-04 | 4.43E-02 |
| 53 | MDD | rs145705679 | C6ORF15 | 6 | 31069749 | 1.34E-04 | 4.30E-02 |
| 54 | TG | rs4333390 | PARKIN | 6 | 162330919 | 4.22E-05 | 1.45E-02 |

| | | | | | | | |
|---|---|---|---|---|---|---|---|
| 55 | CHRONOTYPE | rs651001 | TMEM170B | 6 | 11569402 | 3.57E-05 | 1.48E-02 |
| 56 | BREAST | rs566237 | TMEM170B | 6 | 11543342 | 9.75E-05 | 4.35E-02 |
| 57 | HV | rs6919910 | TMEM170B | 6 | 11546496 | 9.33E-05 | 4.41E-02 |
| 58 | PUT | rs141730255 | AKR1D1 | 7 | 137938550 | 5.24E-05 | 1.63E-02 |
| 59 | SMOKEONSET | rs17171046 | ELMO1 | 7 | 37477863 | 3.90E-05 | 3.88E-02 |
| 60 | COG | rs141347161 | GLI3 | 7 | 42417313 | 3.03E-05 | 2.69E-02 |
| 61 | HV | rs12536071 | GLI3 | 7 | 42428460 | 4.06E-05 | 2.92E-02 |
| 62 | MENARCHE | rs80136991 | GLI3 | 7 | 42429558 | 3.69E-05 | 3.24E-02 |
| 63 | PUT | rs10488631 | TNPO3 | 7 | 128594183 | 2.60E-05 | 7.66E-03 |
| 64 | NEUR | rs8180839 | WIPI2 | 7 | 5239970 | 6.14E-05 | 2.41E-02 |
| 65 | PROSTATE | rs6996532 | BC045738 | 8 | 2417678 | 2.52E-06 | 2.82E-03 |
| 66 | RA | rs6997565 | BC045738 | 8 | 2417597 | 1.58E-06 | 5.08E-03 |
| 67 | WHR | rs6997369 | BC045738 | 8 | 2417440 | 2.80E-06 | 1.46E-02 |
| 68 | SMOKEONSET | rs11786174 | ELP3 | 8 | 28038789 | 1.84E-04 | 3.75E-02 |
| 69 | CRP | rs11250002 | MSRA | 8 | 10257041 | 5.60E-05 | 1.42E-02 |
| 70 | NEUR | rs4292737 | MSRA | 8 | 10259115 | 4.85E-05 | 1.92E-02 |
| 71 | DBP | rs13254942 | MSRA | 8 | 10257678 | 8.50E-05 | 2.75E-02 |
| 72 | PD | rs12541491 | MSRA | 8 | 10250048 | 8.66E-05 | 3.69E-02 |
| 73 | COLLEGE | rs2001338 | MSRA | 8 | 10251217 | 7.64E-05 | 4.78E-02 |
| 74 | AD | rs79496463 | TG | 8 | 133917088 | 7.02E-05 | 3.69E-02 |
| 75 | MDD | rs117204439 | C9ORF72 | 9 | 27607973 | 2.67E-10 | 2.78E-07 |
| 76 | CIGPERDAY | rs2453556 | C9ORF72 | 9 | 27586162 | 5.77E-11 | 3.56E-07 |
| 77 | CBD | rs3849942 | C9ORF72 | 9 | 27543281 | 5.42E-19 | 6.07E-07 |
| 78 | DEPRESSIVE | rs73440960 | C9ORF72 | 9 | 27578647 | 6.58E-17 | 8.46E-07 |
| 79 | MDD | rs773720 | C9ORF72 | 9 | 27586615 | 6.78E-11 | 1.27E-06 |
| 80 | ALC | rs774359 | C9ORF72 | 9 | 27561049 | 1.62E-16 | 1.43E-06 |
| 81 | MENOPAUSE | rs2453565 | C9ORF72 | 9 | 27551040 | 1.25E-17 | 1.96E-06 |
| 82 | PD | rs700791 | C9ORF72 | 9 | 27545960 | 9.53E-18 | 2.55E-06 |
| 83 | BIP_2016 | rs11789520 | C9ORF72 | 9 | 27574515 | 1.08E-17 | 2.61E-06 |

| | | | | | | | |
|---|---|---|---|---|---|---|---|
| 84 | GCOG | rs10967984 | C9ORF72 | 9 | 27555836 | 8.46E-09 | 6.62E-05 |
| 85 | FTD | rs13302855 | C9ORF72 | 9 | 27595995 | 4.15E-06 | 2.22E-04 |
| 86 | MS | rs2282241 | C9ORF72 | 9 | 27572255 | 1.49E-07 | 2.54E-03 |
| 87 | MEMORY | rs4879554 | C9ORF72 | 9 | 27546275 | 2.17E-06 | 1.43E-02 |
| 88 | HDL | rs3849944 | C9ORF72 | 9 | 27560594 | 7.99E-06 | 2.90E-02 |
| 89 | ASD | rs34460171 | C9ORF72 | 9 | 27594491 | 3.60E-06 | 4.16E-02 |
| 90 | BPD | rs147599399 | C9ORF72 | 9 | 27543629 | 8.93E-06 | 4.81E-02 |
| 91 | FTD | rs895021 | MOB3B | 9 | 27484911 | 6.02E-17 | 7.38E-09 |
| 92 | COG | rs10511816 | MOB3B | 9 | 27468461 | 4.61E-11 | 3.76E-08 |
| 93 | BPD | rs7041409 | MOB3B | 9 | 27487221 | 6.91E-14 | 5.73E-08 |
| 94 | WHR | rs10812605 | MOB3B | 9 | 27510360 | 1.08E-10 | 1.51E-07 |
| 95 | PSOR | rs1537712 | MOB3B | 9 | 27508489 | 1.12E-13 | 2.15E-07 |
| 96 | RT | rs10738774 | MOB3B | 9 | 27506051 | 5.01E-13 | 2.20E-07 |
| 97 | SMOKEONSET | rs7046653 | MOB3B | 9 | 27490967 | 1.51E-13 | 2.90E-07 |
| 98 | ASD | rs774352 | MOB3B | 9 | 27516590 | 1.29E-13 | 3.04E-07 |
| 99 | VITILIGO | rs4609281 | MOB3B | 9 | 27514962 | 9.13E-13 | 3.06E-07 |
| 100 | HEIGHT | rs2783009 | MOB3B | 9 | 27514584 | 5.96E-16 | 7.06E-07 |
| 101 | TG | rs700782 | MOB3B | 9 | 27526047 | 1.82E-16 | 1.06E-06 |
| 102 | SLEEP | rs2814707 | MOB3B | 9 | 27536397 | 2.38E-18 | 2.72E-06 |
| 103 | SWB | rs1319236 | MOB3B | 9 | 27507209 | 7.19E-13 | 4.82E-06 |
| 104 | MENARCHE | rs868856 | MOB3B | 9 | 27489251 | 8.25E-14 | 5.32E-06 |
| 105 | LUNG | rs17779457 | MOB3B | 9 | 27488092 | 2.56E-17 | 5.75E-06 |
| 106 | T2D | rs10812604 | MOB3B | 9 | 27496661 | 7.75E-14 | 6.34E-06 |
| 107 | HV | rs774355 | MOB3B | 9 | 27515794 | 1.14E-13 | 1.20E-05 |
| 108 | AGREE | rs2484314 | MOB3B | 9 | 27511284 | 7.29E-14 | 1.24E-05 |
| 109 | UC | rs7019847 | MOBKL2B | 9 | 27491397 | 1.24E-13 | 7.80E-06 |
| 110 | ALC | rs78752429 | MOBKL2B | 9 | 27466680 | 2.14E-09 | 4.12E-05 |
| 111 | T2D | rs139185008 | MOBKL2B | 9 | 27491942 | 8.10E-08 | 7.51E-04 |
| 112 | ALC | rs7024326 | SMARCA2 | 9 | 1903397 | 1.86E-05 | 3.28E-02 |

| | | | | | | | |
|---|---|---|---|---|---|---|---|
| 113 | PSOR | rs7041171 | UGCG | 9 | 114701630 | 2.78E-05 | 4.72E-02 |
| 114 | CBD | rs7944397 | CAT | 11 | 34455309 | 1.14E-05 | 8.81E-03 |
| 115 | RA | rs1962369 | CAT | 11 | 34456941 | 1.66E-06 | 1.01E-02 |
| 116 | CRP | rs7118388 | CAT | 11 | 34454147 | 2.54E-06 | 1.10E-02 |
| 117 | FTD | rs12803540 | CAT | 11 | 34492747 | 9.96E-06 | 2.06E-02 |
| 118 | INTELLIGENCE | rs71472777 | ENDOGENOUS | 11 | 24142935 | 1.34E-05 | 2.54E-02 |
| 119 | BPD | rs140988250 | HCCA2 | 11 | 1489420 | 1.27E-05 | 3.51E-02 |
| 120 | CeD | rs11599981 | HCCA2 | 11 | 1520063 | 1.59E-04 | 4.28E-02 |
| 121 | CRP | rs12369156 | ACADS | 12 | 121167675 | 7.70E-06 | 1.98E-03 |
| 122 | VNR | rs116900480 | AK093124 | 12 | 58656105 | 6.60E-06 | 2.40E-02 |
| 123 | SCZ | rs117704471 | APOLD1 | 12 | 12904565 | 7.20E-05 | 3.91E-02 |
| 124 | DBP | rs593226 | ATXN2 | 12 | 111993886 | 7.43E-05 | 1.94E-02 |
| 125 | RA | rs11065961 | ATXN2 | 12 | 112023067 | 5.93E-05 | 2.45E-02 |
| 126 | ASB | rs75209514 | BC042855 | 12 | 64757407 | 1.07E-06 | 3.04E-03 |
| 127 | SMOKEONSET | rs12308116 | BC042855 | 12 | 64710719 | 3.28E-05 | 1.83E-02 |
| 128 | CHRONOTYPE | rs79031632 | BC042855 | 12 | 64768021 | 1.24E-06 | 3.74E-02 |
| 129 | ADHD | rs11171998 | HSD17B6 | 12 | 57240444 | 1.04E-04 | 3.63E-02 |
| 130 | PUT | rs113247976 | KIF5A | 12 | 57975700 | 1.17E-05 | 2.45E-02 |
| 131 | BIP_2016 | rs144387708 | LOC144742 | 12 | 119702200 | 4.58E-06 | 3.27E-02 |
| 132 | MDD | rs74654358 | TBK1 | 12 | 64881967 | 7.07E-07 | 2.68E-03 |
| 133 | DEPRESSIVE | rs73124200 | TBK1 | 12 | 64949248 | 7.95E-07 | 3.52E-03 |
| 134 | CBD | rs1578303 | HTR2A | 13 | 47963146 | 9.55E-05 | 3.63E-02 |
| 135 | EXTRA | rs10492593 | PCDH9 | 13 | 67494117 | 2.73E-05 | 2.64E-02 |
| 136 | SCZ | rs6420358 | SLITRK1 | 13 | 85289468 | 5.84E-05 | 3.35E-02 |
| 137 | SWB | rs17446243 | TTL/TEL | 13 | 40748931 | 1.17E-05 | 3.41E-03 |
| 138 | BPD | rs17446278 | TTL/TEL | 13 | 40756022 | 1.26E-05 | 4.91E-03 |
| 139 | BREAST | rs61954176 | TTL/TEL | 13 | 40772172 | 4.90E-06 | 1.28E-02 |
| 140 | MENOPAUSE | rs9532508 | TTL/TEL | 13 | 40750048 | 7.92E-06 | 1.61E-02 |
| 141 | ADHD | rs447614 | G2E3 | 14 | 31080799 | 1.97E-05 | 5.47E-03 |

| | | | | | | | |
|---|---|---|---|---|---|---|---|
| 142 | UC | rs447853 | G2E3 | 14 | 31080800 | 1.92E-05 | 2.34E-02 |
| 143 | IBD | rs10139154 | KIAA0917 | 14 | 31147498 | 1.81E-05 | 2.06E-02 |
| 144 | PD | rs151123 | KIAA0917 | 14 | 31093979 | 3.70E-05 | 2.15E-02 |
| 145 | PD | rs12886280 | NUBPL | 14 | 32298659 | 3.31E-06 | 2.18E-02 |
| 146 | PUT | rs2378971 | NUBPL | 14 | 32370206 | 3.68E-05 | 3.21E-02 |
| 147 | GCOG | rs12900374 | BC020234 | 15 | 83410013 | 6.62E-06 | 4.75E-03 |
| 148 | MENOPAUSE | rs2381030 | BTBD1 | 15 | 83741876 | 2.35E-05 | 1.64E-02 |
| 149 | FTD | rs13329195 | BTBD1 | 15 | 83708085 | 1.28E-05 | 1.76E-02 |
| 150 | PSOR | rs6603044 | BTBD1 | 15 | 83683811 | 1.12E-05 | 3.47E-02 |
| 151 | PROSTATE | rs1877240 | FSD2 | 15 | 83438878 | 1.04E-05 | 4.37E-02 |
| 152 | CHRONOTYPE | rs3098553 | GABRG3 | 15 | 27876256 | 2.85E-05 | 3.82E-02 |
| 153 | SBP | rs3097439 | GABRG3 | 15 | 27871229 | 9.83E-05 | 4.52E-02 |
| 154 | VNR | rs7193729 | A2BP1 | 16 | 7145869 | 1.19E-04 | 4.76E-02 |
| 155 | ADHD | rs56024498 | AK057218 | 16 | 76927135 | 8.24E-05 | 3.46E-02 |
| 156 | ICV | rs62068675 | CPNE7 | 16 | 89648291 | 1.51E-04 | 4.81E-02 |
| 157 | AGREE | rs192688752 | PKD1L2 | 16 | 81152628 | 2.43E-05 | 3.63E-02 |
| 158 | PALL | rs2285640 | DHRS11 | 17 | 34951204 | 6.34E-06 | 4.72E-02 |
| 159 | VNR | rs9903355 | GGNBP2 | 17 | 34937221 | 4.93E-06 | 1.99E-03 |
| 160 | MS | rs9906189 | GGNBP2 | 17 | 34905408 | 2.19E-05 | 8.05E-03 |
| 161 | CAD | rs35714695 | KIAA0524 | 17 | 26719788 | 1.16E-08 | 5.75E-06 |
| 162 | BMI | rs739439 | KIAA0524 | 17 | 26723822 | 1.98E-08 | 1.88E-05 |
| 163 | UC | rs34660379 | KIAA0524 | 17 | 26710570 | 3.00E-08 | 3.13E-04 |
| 164 | SCZ | rs2240601 | MSI2 | 17 | 55751112 | 4.82E-05 | 2.89E-02 |
| 165 | PSP | rs7224296 | NSF | 17 | 44800046 | 5.78E-04 | 4.94E-02 |
| 166 | ASD | rs9894834 | NTN1 | 17 | 9129028 | 8.16E-05 | 2.83E-02 |
| 167 | WHR | rs11263770 | PIGW | 17 | 34896877 | 1.00E-05 | 2.74E-03 |
| 168 | HDL | rs11652752 | PITPNC1 | 17 | 65375892 | 7.01E-05 | 2.93E-02 |
| 169 | ALC | rs7209200 | ZFP3 | 17 | 4969940 | 6.44E-05 | 3.75E-02 |
| 170 | HV | rs17187386 | DOK6 | 18 | 67196957 | 6.19E-05 | 2.76E-02 |

| | | | | | | | |
|---|---|---|---|---|---|---|---|
| 171 | BREAST | rs12967284 | SPIRE1 | 18 | 12532098 | 1.62E-05 | 3.31E-02 |
| 172 | OPEN | rs112183647 | GIPC1 | 19 | 14612725 | 2.25E-05 | 4.26E-02 |
| 173 | BPD | rs7258235 | GNG7 | 19 | 2612118 | 1.38E-05 | 2.70E-02 |
| 174 | FTD | rs12608932 | UNC13A | 19 | 17752689 | 1.91E-08 | 1.28E-06 |
| 175 | SLEEP | rs78549703 | UNC13A | 19 | 17749542 | 1.16E-08 | 4.45E-05 |
| 176 | PD | rs6015322 | APCDD1L | 20 | 57206540 | 6.30E-05 | 4.09E-02 |
| 177 | FTD | rs2425220 | DLGAP4 | 20 | 34999440 | 1.30E-04 | 3.06E-02 |
| 178 | NEUR | rs9653747 | BC028044 | 21 | 20041418 | 4.34E-06 | 4.33E-03 |
| 179 | VNR | rs75087725 | C21ORF2 | 21 | 45753117 | 7.52E-11 | 1.65E-07 |
| 180 | AGREE | rs2176039 | NUP50 | 22 | 45585032 | 8.98E-05 | 3.82E-02 |

Table S2. Functional Effects of ALS pleiotropic SNPs (From Conditional FDR Analysis) in a Human Brain Tissue (UKBEC)

| Gene Symbol | SNP | ExprID | Chr | AVEALL | CRBL | FCTX | HIPP | MEDU | OCTX | PUTM | SNIG | TCTX | THAL | WHMT |
|---|---|---|---|---|---|---|---|---|---|---|---|---|---|---|
| CCDC28B | rs10914464 | 2328756 | 1 | 4.70E-03 | 3.60E-01 | 7.50E-01 | 4.60E-01 | 1.00E-01 | 4.40E-01 | 1.90E-01 | 1.50E-01 | 4.80E-01 | 9.60E-02 | 6.70E-01 |
| CYR61 | rs10443173 | 2344891 | 1 | 7.30E-04 | 5.70E-02 | 1.50E-01 | 2.70E-01 | 1.10E-02 | 1.00E-03 | 7.50E-01 | 7.50E-01 | 7.90E-02 | 2.30E-01 | 9.40E-01 |
| NFASC | rs2068667 | 2376257 | 1 | 2.30E-03 | 1.70E-01 | 5.00E-02 | 3.50E-01 | 3.20E-01 | 9.90E-02 | 6.40E-02 | 8.70E-01 | 1.40E-03 | 9.80E-03 | 2.00E-01 |
| BIRC6 | rs6737916 | 2476246 | 2 | 2.50E-03 | 2.80E-01 | 5.90E-01 | 3.10E-02 | 1.40E-01 | 4.10E-02 | 3.40E-02 | 8.10E-01 | 1.10E-02 | 4.60E-01 | 4.40E-01 |
| LOC285045 | rs28407220 | 2476495 | 2 | 1.30E-02 | 7.50E-01 | 8.30E-01 | 1.90E-01 | 1.10E-01 | 3.10E-01 | 7.60E-01 | 7.00E-01 | 1.40E-01 | 5.30E-01 | 9.70E-02 |
| LOC100271832 | rs28407220 | 2476495 | 2 | 1.30E-02 | 7.50E-01 | 8.30E-01 | 1.90E-01 | 1.10E-01 | 3.10E-01 | 7.60E-01 | 7.00E-01 | 1.40E-01 | 5.30E-01 | 9.70E-02 |
| LOC285045 | rs13417671 | 2476495 | 2 | 1.10E-02 | 8.80E-01 | 8.30E-01 | 1.90E-01 | 8.20E-02 | 3.30E-01 | 8.00E-01 | 5.50E-01 | 1.40E-01 | 5.10E-01 | 1.10E-01 |
| LOC100271832 | rs13417671 | 2476495 | 2 | 1.10E-02 | 8.80E-01 | 8.30E-01 | 1.90E-01 | 8.20E-02 | 3.30E-01 | 8.00E-01 | 5.50E-01 | 1.40E-01 | 5.10E-01 | 1.10E-01 |
| MTX2 | rs11695294 | 2517022 | 2 | 1.00E-02 | 6.30E-01 | 2.90E-01 | 4.80E-01 | 1.10E-02 | 1.90E-01 | 1.40E-01 | 3.40E-01 | 1.90E-01 | 2.10E-01 | 1.20E-01 |
| NOSTRIN | rs13387347 | 2514255 | 2 | 3.50E-04 | 2.30E-01 | 5.40E-02 | 5.40E-03 | 2.50E-01 | 3.70E-01 | 3.10E-02 | 1.90E-01 | 4.30E-02 | 1.10E-04 | 2.00E-02 |
| PAX8 | rs7602576 | 2571634 | 2 | 1.00E-03 | 2.60E-01 | 3.70E-01 | 9.60E-01 | 1.90E-01 | 8.30E-04 | 6.00E-01 | 5.20E-02 | 5.60E-01 | 2.20E-01 | 6.30E-01 |
| POU3F3 | rs896444 | 2497825 | 2 | 3.20E-02 | 4.70E-01 | 5.00E-01 | 1.30E-01 | 4.00E-01 | 2.80E-03 | 7.70E-01 | 8.90E-01 | 2.80E-01 | 2.60E-01 | 6.50E-01 |
| SCLY | rs515342 | 2534652 | 2 | 4.50E-03 | 1.00E-01 | 5.00E-01 | 5.40E-01 | 4.50E-03 | 6.80E-01 | 5.70E-01 | 7.40E-01 | 8.40E-01 | 4.90E-03 | 6.90E-02 |
| UBE2F | rs515342 | 2534652 | 2 | 4.50E-03 | 1.00E-01 | 5.00E-01 | 5.40E-01 | 4.50E-03 | 6.80E-01 | 5.70E-01 | 7.40E-01 | 8.40E-01 | 4.90E-03 | 6.90E-02 |
| CNTN6 | rs17042645 | 2607836 | 3 | 3.60E-02 | 1.30E-01 | 3.20E-01 | 6.10E-01 | 5.70E-01 | 6.40E-01 | 5.40E-01 | 2.90E-01 | 7.30E-01 | 4.80E-01 | 4.30E-02 |
| SCN11A | rs6765697 | 2669856 | 3 | 4.40E-03 | 2.60E-01 | 3.60E-01 | 8.80E-01 | 1.00E-01 | 1.00E-01 | 6.80E-01 | 3.60E-01 | 7.00E-01 | 1.00E-03 | 8.00E-01 |
| SCN11A | rs9820623 | 2669856 | 3 | 4.40E-03 | 2.60E-01 | 3.60E-01 | 8.80E-01 | 1.00E-01 | 1.00E-01 | 6.80E-01 | 3.60E-01 | 7.00E-01 | 1.00E-03 | 8.00E-01 |
| SCN11A | rs562545 | 2669856 | 3 | 3.40E-03 | 7.00E-01 | 6.60E-01 | 2.70E-01 | 3.50E-01 | 4.60E-02 | 1.70E-01 | 1.00E-01 | 8.10E-02 | 1.80E-01 | 8.00E-01 |
| SCN5A | rs1464047 | 2669671 | 3 | 9.60E-05 | 9.30E-01 | 7.70E-01 | 5.50E-02 | 2.10E-02 | 4.40E-02 | 7.50E-01 | 3.30E-01 | 9.20E-01 | 2.00E-02 | 5.80E-02 |
| WDR48 | rs1768208 | 2617970 | 3 | 2.30E-03 | 2.20E-01 | 1.60E-02 | 2.20E-01 | 8.00E-01 | 1.10E-01 | 9.10E-01 | 5.20E-01 | 8.40E-01 | 8.40E-03 | 3.10E-01 |
| C4orf10 | rs62637887 | 2715724 | 4 | 9.40E-04 | 2.40E-03 | 2.30E-01 | 5.10E-02 | 5.20E-01 | 1.80E-02 | 7.40E-01 | 1.50E-01 | 3.90E-02 | 7.30E-01 | 3.60E-01 |
| LOC646358 | rs1159918 | 2779457 | 4 | 5.10E-03 | 3.90E-01 | 7.40E-02 | 2.80E-01 | 4.10E-01 | 3.70E-02 | 3.30E-01 | 3.10E-02 | 2.10E-01 | 6.50E-01 | 7.00E-01 |
| DNAJB14 | rs1159918 | 2779457 | 4 | 5.10E-03 | 3.90E-01 | 7.40E-02 | 2.80E-01 | 4.10E-01 | 3.70E-02 | 3.30E-01 | 3.10E-02 | 2.10E-01 | 6.50E-01 | 7.00E-01 |
| HTRA3 | rs10938692 | 2717705 | 4 | 6.90E-03 | 3.00E-01 | 9.50E-01 | 9.20E-04 | 5.00E-02 | 3.20E-01 | 2.70E-01 | 8.40E-01 | 1.50E-01 | 6.80E-01 | 4.30E-02 |
| LOC345051 | rs10011222 | 2746338 | 4 | 1.70E-02 | 2.50E-01 | 1.80E-01 | 1.80E-01 | 2.70E-01 | 5.90E-01 | 9.80E-01 | 3.20E-01 | 6.10E-01 | 3.30E-01 | 8.60E-02 |
| PCGF3 | rs12646225 | 2714265 | 4 | 1.10E-03 | 5.30E-02 | 2.40E-01 | 2.20E-02 | 8.80E-02 | 4.90E-01 | 4.60E-02 | 8.50E-03 | 2.10E-01 | 9.20E-02 | 1.70E-01 |

| | | | | | | | | | | | | | | |
|---|---|---|---|---|---|---|---|---|---|---|---|---|---|---|
| RGS12 | rs61789437 | 2716051 | 4 | 4.10E-05 | 4.30E-01 | 4.30E-01 | 4.60E-01 | 2.30E-01 | 2.40E-01 | 8.40E-04 | 2.80E-01 | 4.00E-02 | 3.90E-02 | 2.10E-02 |
| RGS12 | rs4974674 | 2716051 | 4 | 2.20E-04 | 3.80E-01 | 9.90E-01 | 7.90E-01 | 6.10E-01 | 7.30E-01 | 3.40E-05 | 5.50E-02 | 1.00E-01 | 1.20E-02 | 8.30E-02 |
| RGS12 | rs762837 | 2716051 | 4 | 3.70E-04 | 4.30E-01 | 9.40E-01 | 7.80E-01 | 5.70E-01 | 7.40E-01 | 4.10E-05 | 5.80E-02 | 1.20E-01 | 1.70E-02 | 1.10E-01 |
| RGS12 | rs4974650 | 2716051 | 4 | 1.40E-05 | 3.00E-01 | 3.90E-01 | 4.10E-01 | 2.40E-01 | 1.30E-01 | 1.40E-03 | 3.10E-01 | 2.00E-02 | 2.40E-02 | 2.80E-02 |
| SH3BP2 | rs430979 | 2715620 | 4 | 5.40E-03 | 1.40E-01 | 1.80E-01 | 4.30E-01 | 8.60E-02 | 2.50E-02 | 4.30E-03 | 1.80E-02 | 2.30E-01 | 6.20E-02 | 2.60E-02 |
| APC | rs17326496 | 2824220 | 5 | 1.20E-02 | 8.40E-01 | 1.90E-01 | 9.70E-01 | 9.90E-01 | 7.60E-02 | 1.80E-01 | 7.80E-01 | 3.20E-01 | 2.10E-03 | 4.50E-01 |
| BNIP1 | rs538622 | t2841528 | 5 | 1.10E-03 | 1.50E-01 | 2.40E-01 | 3.00E-01 | 7.50E-03 | 2.40E-02 | 3.80E-01 | 1.60E-01 | 6.40E-01 | 9.40E-02 | 5.90E-01 |
| BNIP1 | rs7735726 | t2841528 | 5 | 1.10E-03 | 1.50E-01 | 2.50E-01 | 3.00E-01 | 7.50E-03 | 2.40E-02 | 3.80E-01 | 1.60E-01 | 6.50E-01 | 9.40E-02 | 5.90E-01 |
| DCTN4 | rs10463311 | 2881559 | 5 | 3.60E-05 | 1.70E-03 | 6.20E-02 | 1.00E-02 | 9.20E-01 | 1.10E-01 | 1.20E-02 | 8.30E-02 | 3.90E-01 | 2.30E-02 | 7.80E-01 |
| DCTN4 | rs3828599 | 2881559 | 5 | 5.80E-04 | 1.80E-03 | 7.80E-02 | 5.10E-02 | 7.80E-01 | 1.60E-01 | 5.10E-02 | 1.40E-01 | 5.10E-01 | 3.90E-02 | 8.50E-01 |
| DCTN4 | rs4958872 | 2881559 | 5 | 3.60E-05 | 1.70E-03 | 6.20E-02 | 1.00E-02 | 9.20E-01 | 1.10E-01 | 1.20E-02 | 8.30E-02 | 3.90E-01 | 2.30E-02 | 7.80E-01 |
| EGFLAM | rs10050775 | 2807228 | 5 | 3.90E-02 | 5.00E-01 | 4.10E-01 | 5.60E-01 | 1.60E-02 | 2.20E-01 | 1.60E-01 | 3.80E-04 | 2.00E-01 | 8.30E-01 | 8.70E-01 |
| EGFLAM | rs10059028 | 2807228 | 5 | 3.10E-02 | 2.40E-01 | 7.50E-01 | 6.80E-01 | 2.60E-02 | 2.20E-01 | 5.80E-01 | 4.10E-06 | 3.00E-01 | 8.30E-01 | 2.90E-01 |
| LOC134466 | rs4958888 | 2881638 | 5 | 3.30E-03 | 2.60E-02 | 8.00E-01 | 9.10E-01 | 1.50E-01 | 6.90E-02 | 2.70E-01 | 7.50E-02 | 1.20E-01 | 6.70E-01 | 8.70E-01 |
| LOC134466 | rs8177427 | 2881614 | 5 | 6.70E-06 | 5.70E-01 | 7.10E-04 | 7.50E-01 | 1.80E-01 | 6.00E-01 | 7.80E-02 | 4.30E-01 | 8.80E-02 | 4.10E-01 | 3.90E-03 |
| LOC134466 | rs3762996 | 2881638 | 5 | 3.30E-03 | 2.60E-02 | 8.00E-01 | 9.10E-01 | 1.50E-01 | 6.90E-02 | 2.70E-01 | 7.50E-02 | 1.20E-01 | 6.70E-01 | 8.70E-01 |
| LOC134466 | rs11747926 | 2881614 | 5 | 1.30E-06 | 6.50E-01 | 2.10E-03 | 8.50E-01 | 1.90E-01 | 2.00E-01 | 6.70E-02 | 1.60E-01 | 1.10E-01 | 1.70E-01 | 4.40E-03 |
| LOC134466 | rs4958434 | 2881614 | 5 | 6.70E-06 | 5.70E-01 | 7.10E-04 | 7.50E-01 | 1.80E-01 | 6.00E-01 | 7.80E-02 | 4.30E-01 | 8.80E-02 | 4.10E-01 | 3.90E-03 |
| TNIP1 | rs17111695 | 2881696 | 5 | 7.30E-04 | 4.20E-02 | 2.20E-01 | 3.30E-04 | 5.10E-01 | 6.70E-02 | 9.70E-01 | 4.30E-02 | 4.00E-02 | 1.50E-01 | 1.90E-01 |
| TXNDC15 | rs7728741 | 2829575 | 5 | 2.60E-14 | 6.80E-07 | 1.30E-06 | 7.50E-06 | 1.50E-04 | 2.60E-05 | 5.40E-04 | 4.50E-02 | 3.90E-05 | 2.30E-08 | 1.30E-03 |
| ZNF300 | rs4958888 | 2881638 | 5 | 3.30E-03 | 2.60E-02 | 8.00E-01 | 9.10E-01 | 1.50E-01 | 6.90E-02 | 2.70E-01 | 7.50E-02 | 1.20E-01 | 6.70E-01 | 8.70E-01 |
| ZNF300 | rs8177427 | 2881614 | 5 | 6.70E-06 | 5.70E-01 | 7.10E-04 | 7.50E-01 | 1.80E-01 | 6.00E-01 | 7.80E-02 | 4.30E-01 | 8.80E-02 | 4.10E-01 | 3.90E-03 |
| ZNF300 | rs3762996 | 2881638 | 5 | 3.30E-03 | 2.60E-02 | 8.00E-01 | 9.10E-01 | 1.50E-01 | 6.90E-02 | 2.70E-01 | 7.50E-02 | 1.20E-01 | 6.70E-01 | 8.70E-01 |
| ZNF300 | rs11747926 | 2881614 | 5 | 1.30E-06 | 6.50E-01 | 2.10E-03 | 8.50E-01 | 1.90E-01 | 2.00E-01 | 6.70E-02 | 1.60E-01 | 1.10E-01 | 1.70E-01 | 4.40E-03 |
| ZNF300 | rs4958434 | 2881614 | 5 | 6.70E-06 | 5.70E-01 | 7.10E-04 | 7.50E-01 | 1.80E-01 | 6.00E-01 | 7.80E-02 | 4.30E-01 | 8.80E-02 | 4.10E-01 | 3.90E-03 |
| LOC134466 | rs4958888 | 2881638 | 5 | 3.30E-03 | 2.60E-02 | 8.00E-01 | 9.10E-01 | 1.50E-01 | 6.90E-02 | 2.70E-01 | 7.50E-02 | 1.20E-01 | 6.70E-01 | 8.70E-01 |
| ZNF300 | rs4958888 | 2881638 | 5 | 3.30E-03 | 2.60E-02 | 8.00E-01 | 9.10E-01 | 1.50E-01 | 6.90E-02 | 2.70E-01 | 7.50E-02 | 1.20E-01 | 6.70E-01 | 8.70E-01 |
| LOC134466 | rs8177427 | 2881614 | 5 | 6.70E-06 | 5.70E-01 | 7.10E-04 | 7.50E-01 | 1.80E-01 | 6.00E-01 | 7.80E-02 | 4.30E-01 | 8.80E-02 | 4.10E-01 | 3.90E-03 |
| ZNF300 | rs8177427 | 2881614 | 5 | 6.70E-06 | 5.70E-01 | 7.10E-04 | 7.50E-01 | 1.80E-01 | 6.00E-01 | 7.80E-02 | 4.30E-01 | 8.80E-02 | 4.10E-01 | 3.90E-03 |

| | | | | | | | | | | | | | | |
|---|---|---|---|---|---|---|---|---|---|---|---|---|---|---|
| LOC134466 | rs11747926 | 2881614 | 5 | 1.30E-06 | 6.50E-01 | 2.10E-03 | 8.50E-01 | 1.90E-01 | 2.00E-01 | 6.70E-02 | 1.60E-01 | 1.10E-01 | 1.70E-01 | 4.40E-03 |
| ZNF300 | rs11747926 | 2881614 | 5 | 1.30E-06 | 6.50E-01 | 2.10E-03 | 8.50E-01 | 1.90E-01 | 2.00E-01 | 6.70E-02 | 1.60E-01 | 1.10E-01 | 1.70E-01 | 4.40E-03 |
| LOC134466 | rs3762996 | 2881638 | 5 | 3.30E-03 | 2.60E-02 | 8.00E-01 | 9.10E-01 | 1.50E-01 | 6.90E-02 | 2.70E-01 | 7.50E-02 | 1.20E-01 | 6.70E-01 | 8.70E-01 |
| ZNF300 | rs3762996 | 2881638 | 5 | 3.30E-03 | 2.60E-02 | 8.00E-01 | 9.10E-01 | 1.50E-01 | 6.90E-02 | 2.70E-01 | 7.50E-02 | 1.20E-01 | 6.70E-01 | 8.70E-01 |
| LOC134466 | rs4958434 | 2881614 | 5 | 6.70E-06 | 5.70E-01 | 7.10E-04 | 7.50E-01 | 1.80E-01 | 6.00E-01 | 7.80E-02 | 4.30E-01 | 8.80E-02 | 4.10E-01 | 3.90E-03 |
| ZNF300 | rs4958434 | 2881614 | 5 | 6.70E-06 | 5.70E-01 | 7.10E-04 | 7.50E-01 | 1.80E-01 | 6.00E-01 | 7.80E-02 | 4.30E-01 | 8.80E-02 | 4.10E-01 | 3.90E-03 |
| AGPAT4 | rs4333390 | 2983045 | 6 | 2.40E-03 | 1.90E-01 | 7.60E-01 | 3.50E-02 | 9.00E-01 | 2.20E-01 | 6.80E-01 | 9.50E-02 | 4.70E-03 | 9.90E-01 | 6.90E-02 |
| ELOVL2 | rs6919910 | 2941737 | 6 | 2.00E-02 | 5.90E-01 | 7.00E-01 | 8.10E-01 | 2.10E-01 | 9.80E-01 | 7.80E-02 | 8.40E-02 | 1.50E-02 | 9.80E-01 | 1.00E+00 |
| GCNT6 | rs566237 | 2894647 | 6 | 2.20E-02 | 1.30E-01 | 3.90E-01 | 4.90E-01 | 5.40E-01 | 1.50E-02 | 2.80E-01 | 4.20E-02 | 7.00E-02 | 6.90E-01 | 4.40E-01 |
| HINT3 | rs1543705 | 2924607 | 6 | 4.90E-03 | 6.50E-01 | 2.40E-01 | 3.70E-01 | 1.10E-01 | 2.60E-01 | 6.50E-01 | 1.80E-01 | 2.30E-01 | 2.10E-01 | 5.70E-01 |
| TMEM170B | rs651001 | t2895031 | 6 | 1.80E-02 | 2.70E-01 | 1.60E-02 | 5.70E-01 | 8.80E-01 | 1.80E-01 | 6.70E-01 | 4.40E-01 | 1.10E-01 | 2.50E-01 | 6.20E-01 |
| VARS2 | rs145705679 | 2902084 | 6 | 9.50E-06 | 6.40E-01 | 1.60E-02 | 4.10E-02 | 7.50E-02 | 7.40E-02 | 2.80E-05 | 2.60E-02 | 1.60E-01 | 6.80E-01 | 1.80E-01 |
| AHCYL2 | rs10488631 | 3023388 | 7 | 7.60E-04 | 8.60E-01 | 8.80E-02 | 9.50E-01 | 2.40E-02 | 1.10E-02 | 1.90E-01 | 8.20E-01 | 2.00E-02 | 1.30E-02 | 2.90E-01 |
| AMPH | rs17171046 | 3046747 | 7 | 3.50E-02 | 7.90E-01 | 4.70E-01 | 7.90E-01 | 9.90E-03 | 1.80E-01 | 2.60E-01 | 3.30E-01 | 5.10E-01 | 5.00E-02 | 3.30E-01 |
| CREB3L2 | rs141730255 | 3075148 | 7 | 1.10E-02 | 4.00E-01 | 2.30E-02 | 4.80E-01 | 1.20E-02 | 1.40E-02 | 5.20E-01 | 3.40E-01 | 5.30E-01 | 5.80E-01 | 2.90E-01 |
| HECW1 | rs141347161 | 2999375 | 7 | 9.70E-03 | 6.30E-01 | 9.50E-01 | 7.90E-01 | 5.50E-01 | 4.90E-02 | 1.00E+00 | 7.40E-02 | 9.20E-01 | 1.20E-01 | 6.40E-02 |
| HECW1 | rs12536071 | 2999375 | 7 | 8.70E-03 | 6.80E-01 | 9.40E-01 | 7.40E-01 | 5.00E-01 | 6.30E-02 | 9.50E-01 | 6.80E-02 | 9.20E-01 | 1.30E-01 | 5.00E-02 |
| HECW1 | rs80136991 | 2999375 | 7 | 8.70E-03 | 6.80E-01 | 9.50E-01 | 7.40E-01 | 5.00E-01 | 6.40E-02 | 9.50E-01 | 6.70E-02 | 9.10E-01 | 1.40E-01 | 4.90E-02 |
| ARHGEF10 | rs6997369 | 3082902 | 8 | 1.60E-02 | 7.40E-01 | 6.50E-01 | 1.20E-01 | 7.50E-01 | 3.60E-01 | 6.40E-01 | 3.40E-01 | 8.50E-02 | 6.00E-02 | 1.30E-01 |
| ARHGEF10 | rs6997565 | 3082902 | 8 | 1.60E-02 | 7.40E-01 | 6.50E-01 | 1.20E-01 | 7.50E-01 | 3.60E-01 | 6.40E-01 | 3.40E-01 | 8.50E-02 | 6.00E-02 | 1.30E-01 |
| ARHGEF10 | rs6996532 | 3082902 | 8 | 1.60E-02 | 7.40E-01 | 6.50E-01 | 1.20E-01 | 7.50E-01 | 3.60E-01 | 6.40E-01 | 3.40E-01 | 8.50E-02 | 6.00E-02 | 1.30E-01 |
| LOC100129129 | rs2001338 | t3124380 | 8 | 1.90E-04 | 9.30E-03 | 2.10E-05 | 3.60E-01 | 2.40E-01 | 4.40E-01 | 2.50E-01 | 1.10E-02 | 2.50E-01 | 8.80E-01 | 6.80E-03 |
| LOC100129129 | rs4292737 | t3124380 | 8 | 7.30E-05 | 2.60E-02 | 6.40E-04 | 4.40E-01 | 5.80E-02 | 4.70E-01 | 2.00E-01 | 3.50E-03 | 1.70E-01 | 4.30E-01 | 1.00E-02 |
| LOC100129129 | rs11250002 | t3124380 | 8 | 8.10E-05 | 9.30E-03 | 1.50E-04 | 2.50E-01 | 2.00E-01 | 3.90E-01 | 1.60E-01 | 8.90E-03 | 2.70E-01 | 6.50E-01 | 2.30E-03 |
| LOC100129129 | rs12541491 | t3124380 | 8 | 2.30E-04 | 9.90E-03 | 1.30E-05 | 4.90E-01 | 2.50E-01 | 4.40E-01 | 2.10E-01 | 1.10E-02 | 2.50E-01 | 9.90E-01 | 6.30E-03 |
| LOC100129129 | rs13254942 | t3124380 | 8 | 1.00E-04 | 9.40E-03 | 8.30E-05 | 3.70E-01 | 2.10E-01 | 4.00E-01 | 1.40E-01 | 8.70E-03 | 2.60E-01 | 7.50E-01 | 2.50E-03 |
| OC90 | rs79496463 | 3153948 | 8 | 1.10E-02 | 9.30E-01 | 9.70E-02 | 6.80E-01 | 4.40E-01 | 5.90E-01 | 2.90E-01 | 9.40E-01 | 1.70E-02 | 6.90E-02 | 1.20E-01 |
| TRIM35 | rs11786174 | 3128958 | 8 | 1.50E-02 | 1.20E-01 | 7.30E-01 | 9.30E-01 | 7.20E-03 | 1.00E-01 | 2.20E-01 | 3.60E-01 | 4.10E-01 | 4.80E-01 | 6.50E-01 |
| C9orf11 | rs700782 | 3202306 | 9 | 5.50E-03 | 9.60E-01 | 6.80E-01 | 1.20E-02 | 3.60E-02 | 9.30E-01 | 1.20E-01 | 5.60E-02 | 7.80E-01 | 1.70E-02 | 5.40E-01 |

| | | | | | | | | | | | | | | |
|---|---|---|---|---|---|---|---|---|---|---|---|---|---|---|
| C9orf11 | rs895021 | 3202306 | 9 | 3.50E-03 | 9.20E-01 | 7.30E-01 | 2.20E-02 | 3.70E-02 | 8.90E-01 | 1.20E-01 | 7.30E-02 | 7.00E-01 | 9.50E-03 | 4.70E-01 |
| C9orf11 | rs17779457 | 3202306 | 9 | 5.50E-03 | 9.60E-01 | 6.80E-01 | 1.20E-02 | 3.60E-02 | 9.30E-01 | 1.20E-01 | 5.60E-02 | 7.80E-01 | 1.70E-02 | 5.40E-01 |
| C9orf11 | rs2453556 | 3202306 | 9 | 2.00E-03 | 7.80E-01 | 4.70E-01 | 1.10E-01 | 4.60E-01 | 5.60E-03 | 3.60E-01 | 7.40E-02 | 9.60E-01 | 1.30E-02 | 1.10E-01 |
| C9orf11 | rs773720 | 3202306 | 9 | 2.10E-03 | 7.60E-01 | 4.50E-01 | 1.20E-01 | 4.60E-01 | 5.20E-03 | 3.40E-01 | 7.30E-02 | 9.60E-01 | 1.40E-02 | 1.10E-01 |
| C9orf84 | rs7041171 | 3220767 | 9 | 1.90E-02 | 4.40E-01 | 8.80E-01 | 8.70E-01 | 7.00E-02 | 9.00E-01 | 3.00E-02 | 6.70E-01 | 1.90E-01 | 6.80E-02 | 8.40E-01 |
| LRRC19 | rs3849944 | 3202230 | 9 | 3.80E-02 | 4.50E-01 | 2.40E-01 | 3.90E-01 | 7.70E-01 | 8.80E-01 | 7.20E-01 | 3.80E-01 | 9.00E-01 | 1.50E-01 | 2.10E-01 |
| LRRC19 | rs13302855 | 3202231 | 9 | 1.30E-02 | 3.60E-01 | 4.40E-01 | 2.80E-02 | 9.80E-01 | 2.70E-02 | 1.70E-01 | 5.20E-01 | 4.70E-01 | 7.50E-01 | 6.60E-01 |
| LRRC19 | rs34460171 | 3202231 | 9 | 1.20E-02 | 3.80E-01 | 4.50E-01 | 2.90E-02 | 9.70E-01 | 2.40E-02 | 1.50E-01 | 4.70E-01 | 4.20E-01 | 7.70E-01 | 7.10E-01 |
| LRRC19 | rs4879554 | 3202230 | 9 | 5.20E-02 | 5.00E-01 | 2.00E-01 | 1.90E-01 | 9.70E-01 | 9.20E-01 | 9.20E-01 | 5.10E-01 | 9.00E-01 | 9.70E-02 | 5.60E-01 |
| MOBKL2B | rs3849942 | 3202320 | 9 | 4.80E-03 | 8.80E-02 | 1.10E-01 | 1.80E-01 | 5.00E-01 | 9.70E-02 | 3.70E-03 | 8.20E-01 | 9.00E-01 | 1.00E+00 | 7.50E-01 |
| MOBKL2B | rs2814707 | 3202320 | 9 | 5.90E-03 | 1.20E-01 | 1.10E-01 | 2.50E-01 | 4.20E-01 | 8.30E-02 | 5.80E-03 | 8.20E-01 | 9.10E-01 | 9.30E-01 | 7.30E-01 |
| PLAA | rs2282241 | 3202202 | 9 | 6.70E-03 | 9.30E-02 | 1.00E-01 | 7.30E-02 | 1.80E-02 | 4.40E-02 | 7.20E-01 | 9.30E-03 | 2.40E-01 | 6.40E-01 | 9.70E-01 |
| PLAA | rs10967984 | 3202202 | 9 | 4.00E-03 | 1.90E-01 | 7.40E-02 | 2.60E-02 | 4.60E-02 | 8.30E-02 | 3.20E-01 | 1.30E-02 | 5.30E-01 | 7.60E-01 | 4.50E-01 |
| PLAA | rs2783009 | 3202175 | 9 | 3.70E-03 | 6.00E-02 | 1.10E-01 | 8.80E-01 | 3.60E-01 | 2.10E-01 | 9.10E-02 | 4.00E-01 | 2.40E-01 | 4.30E-01 | 2.40E-01 |
| PLAA | rs11789520 | 3202175 | 9 | 6.70E-03 | 1.60E-01 | 6.50E-02 | 9.30E-01 | 5.20E-01 | 1.80E-01 | 1.50E-01 | 3.20E-01 | 2.00E-01 | 2.90E-01 | 2.40E-01 |
| SMARCA2 | rs7024326 | 3160073 | 9 | 6.60E-04 | 8.90E-02 | 6.90E-02 | 2.10E-01 | 8.30E-02 | 5.50E-01 | 3.40E-02 | 2.30E-01 | 7.60E-02 | 5.30E-02 | 3.50E-01 |
| TEK | rs10812605 | 3165831 | 9 | 8.20E-04 | 4.10E-01 | 6.50E-02 | 3.90E-01 | 2.50E-01 | 4.60E-01 | 4.40E-01 | 8.20E-02 | 6.00E-02 | 1.00E+00 | 1.20E-01 |
| TEK | rs10511816 | 3165865 | 9 | 2.00E-04 | 1.30E-01 | 1.40E-01 | 1.30E-02 | 6.20E-01 | 5.50E-02 | 1.40E-02 | 6.60E-01 | 4.30E-01 | 3.80E-01 | 2.50E-01 |
| TEK | rs10738774 | 3165859 | 9 | 3.90E-05 | 1.50E-01 | 2.80E-01 | 6.70E-02 | 4.00E-01 | 4.70E-02 | 2.70E-01 | 1.50E-03 | 9.40E-03 | 8.50E-01 | 9.00E-03 |
| TEK | rs774355 | 3165859 | 9 | 3.90E-05 | 1.50E-01 | 2.80E-01 | 6.70E-02 | 4.00E-01 | 4.70E-02 | 2.70E-01 | 1.50E-03 | 9.40E-03 | 8.50E-01 | 9.00E-03 |
| TEK | rs774359 | 3165859 | 9 | 1.80E-03 | 5.30E-01 | 4.70E-01 | 1.00E-01 | 6.30E-01 | 6.30E-02 | 8.90E-01 | 2.50E-03 | 1.30E-02 | 8.40E-01 | 3.60E-02 |
| TEK | rs7041409 | 3165831 | 9 | 1.30E-04 | 6.50E-01 | 2.80E-02 | 1.90E-01 | 1.90E-01 | 3.60E-01 | 2.50E-01 | 5.00E-02 | 2.00E-01 | 7.20E-01 | 3.60E-02 |
| TEK | rs78752429 | 3165865 | 9 | 2.80E-05 | 1.20E-01 | 2.60E-01 | 3.10E-03 | 4.40E-01 | 1.20E-01 | 2.40E-02 | 2.20E-01 | 9.70E-01 | 5.80E-01 | 2.20E-02 |
| TEK | rs2484314 | 3165859 | 9 | 3.90E-05 | 1.50E-01 | 2.80E-01 | 6.70E-02 | 4.00E-01 | 4.70E-02 | 2.70E-01 | 1.50E-03 | 9.40E-03 | 8.50E-01 | 9.00E-03 |
| TEK | rs73440960 | 3165859 | 9 | 7.70E-03 | 6.50E-01 | 7.90E-01 | 2.80E-01 | 4.50E-01 | 4.60E-02 | 8.60E-01 | 7.30E-03 | 9.90E-02 | 9.20E-01 | 3.50E-03 |
| TEK | rs10812604 | 3165859 | 9 | 9.10E-05 | 1.80E-01 | 2.50E-01 | 5.50E-02 | 4.10E-01 | 5.30E-02 | 4.20E-01 | 1.50E-03 | 1.70E-02 | 9.30E-01 | 8.60E-03 |
| TEK | rs700791 | 3165831 | 9 | 4.90E-03 | 6.60E-01 | 7.80E-02 | 2.70E-01 | 1.50E-01 | 3.90E-01 | 7.40E-01 | 4.40E-02 | 3.00E-01 | 9.20E-01 | 1.30E-01 |
| TEK | rs1319236 | 3165859 | 9 | 4.00E-05 | 1.50E-01 | 3.00E-01 | 6.50E-02 | 3.50E-01 | 6.10E-02 | 2.90E-01 | 1.50E-03 | 7.50E-03 | 8.60E-01 | 8.50E-03 |
| TEK | rs868856 | 3165859 | 9 | 9.20E-05 | 1.80E-01 | 2.50E-01 | 5.50E-02 | 4.10E-01 | 5.40E-02 | 4.20E-01 | 1.50E-03 | 1.70E-02 | 9.40E-01 | 8.80E-03 |

| | | | | | | | | | | | | | | |
|---|---|---|---|---|---|---|---|---|---|---|---|---|---|---|
| TEK | rs7019847 | 3165859 | 9 | 8.80E-05 | 2.00E-01 | 2.50E-01 | 5.00E-02 | 3.80E-01 | 4.90E-02 | 4.20E-01 | 1.60E-03 | 1.70E-02 | 9.60E-01 | 9.50E-03 |
| TEK | rs774352 | 3165859 | 9 | 3.90E-05 | 1.50E-01 | 2.80E-01 | 6.70E-02 | 4.00E-01 | 4.70E-02 | 2.70E-01 | 1.50E-03 | 9.40E-03 | 8.50E-01 | 9.00E-03 |
| TEK | rs7046653 | 3165859 | 9 | 9.20E-05 | 1.80E-01 | 2.50E-01 | 5.50E-02 | 4.10E-01 | 5.40E-02 | 4.20E-01 | 1.50E-03 | 1.70E-02 | 9.40E-01 | 8.80E-03 |
| TEK | rs2453565 | 3165831 | 9 | 5.00E-03 | 6.60E-01 | 7.80E-02 | 2.70E-01 | 1.50E-01 | 3.90E-01 | 7.50E-01 | 4.40E-02 | 3.10E-01 | 9.20E-01 | 1.30E-01 |
| TEK | rs1537712 | 3165859 | 9 | 3.90E-05 | 1.50E-01 | 2.80E-01 | 6.70E-02 | 4.00E-01 | 4.70E-02 | 2.70E-01 | 1.50E-03 | 9.40E-03 | 8.50E-01 | 9.00E-03 |
| TEK | rs4609281 | 3165859 | 9 | 3.90E-05 | 1.50E-01 | 2.80E-01 | 6.70E-02 | 4.00E-01 | 4.70E-02 | 2.70E-01 | 1.50E-03 | 9.40E-03 | 8.50E-01 | 9.00E-03 |
| CAT | rs12803540 | 3326416 | 11 | 3.10E-03 | 1.20E-01 | 8.50E-03 | 6.50E-02 | 7.40E-04 | 2.40E-01 | 1.50E-01 | 6.90E-01 | 2.10E-01 | 6.20E-01 | 1.90E-01 |
| C11orf91 | rs7944397 | 3368745 | 11 | 1.20E-02 | 2.50E-01 | 8.60E-01 | 5.30E-01 | 5.90E-01 | 2.20E-02 | 5.90E-01 | 5.90E-01 | 3.20E-01 | 5.80E-01 | 1.80E-01 |
| CD59 | rs7944397 | 3368745 | 11 | 1.20E-02 | 2.50E-01 | 8.60E-01 | 5.30E-01 | 5.90E-01 | 2.20E-02 | 5.90E-01 | 5.90E-01 | 3.20E-01 | 5.80E-01 | 1.80E-01 |
| C11orf91 | rs7118388 | 3368742 | 11 | 1.40E-02 | 9.40E-01 | 4.70E-01 | 1.20E-01 | 6.40E-01 | 7.40E-01 | 1.20E-01 | 3.20E-01 | 5.00E-01 | 2.70E-01 | 1.50E-03 |
| CD59 | rs7118388 | 3368742 | 11 | 1.40E-02 | 9.40E-01 | 4.70E-01 | 1.20E-01 | 6.40E-01 | 7.40E-01 | 1.20E-01 | 3.20E-01 | 5.00E-01 | 2.70E-01 | 1.50E-03 |
| FBXO3 | rs1962369 | 3368750 | 11 | 1.40E-02 | 5.90E-01 | 2.70E-02 | 1.10E-02 | 7.90E-02 | 1.00E-02 | 9.60E-01 | 7.10E-01 | 5.90E-01 | 4.40E-01 | 7.50E-01 |
| TSPAN32 | rs11599981 | 3317293 | 11 | 1.20E-04 | 1.70E-01 | 8.50E-01 | 2.10E-02 | 1.00E-01 | 4.40E-01 | 2.60E-01 | 4.50E-01 | 5.60E-02 | 8.90E-01 | 5.40E-01 |
| C12orf56 | rs12308116 | 3460013 | 12 | 4.30E-02 | 7.40E-02 | 2.20E-01 | 7.10E-02 | 9.40E-01 | 4.40E-01 | 1.70E-01 | 6.60E-01 | 7.20E-01 | 2.90E-01 | 4.90E-01 |
| C12orf56 | rs74654358 | 3460006 | 12 | 5.50E-02 | 2.70E-01 | 6.10E-01 | 1.30E-01 | 4.70E-01 | 5.20E-01 | 4.20E-01 | 1.20E-01 | 1.30E-01 | 5.10E-02 | 6.80E-03 |
| CUX2 | rs593226 | 3431843 | 12 | 3.70E-03 | 5.10E-01 | 1.50E-03 | 7.20E-01 | 1.40E-01 | 3.70E-01 | 1.40E-01 | 8.50E-01 | 5.90E-01 | 8.20E-01 | 5.80E-02 |
| CUX2 | rs11065961 | 3431843 | 12 | 3.50E-03 | 5.20E-01 | 1.70E-03 | 6.70E-01 | 1.30E-01 | 3.80E-01 | 1.50E-01 | 8.10E-01 | 6.50E-01 | 8.70E-01 | 5.80E-02 |
| TBK1 | rs73124200 | 3460071 | 12 | 1.60E-02 | 3.10E-01 | 5.10E-01 | 3.80E-01 | 1.20E-02 | 1.00E-01 | 1.70E-01 | 6.40E-01 | 9.80E-02 | 7.10E-01 | 5.50E-01 |
| TMEM5 | rs79031632 | 3419606 | 12 | 6.90E-02 | 2.10E-01 | 8.00E-01 | 5.30E-01 | 4.50E-01 | 6.60E-01 | 2.20E-01 | 6.30E-01 | 6.00E-01 | 6.30E-01 | 7.20E-03 |
| WIBG | rs11171998 | 3457316 | 12 | 6.80E-04 | 2.70E-01 | 8.50E-02 | 1.80E-01 | 2.30E-01 | 2.20E-01 | 8.80E-04 | 4.10E-02 | 4.80E-01 | 2.20E-02 | 1.50E-01 |
| COG6 | rs61954176 | 3486436 | 13 | 6.80E-02 | 2.30E-01 | 8.50E-01 | 3.90E-01 | 5.70E-01 | 4.60E-01 | 6.60E-01 | 4.40E-01 | 1.60E-01 | 3.60E-01 | 5.00E-02 |
| PCDH9 | rs10492593 | 3516677 | 13 | 1.80E-01 | 7.10E-01 | 5.20E-01 | 6.10E-01 | 4.10E-01 | 3.30E-01 | 5.20E-01 | 5.90E-01 | 9.00E-01 | 4.80E-01 | 3.20E-02 |
| SLITRK1 | rs6420358 | 3519632 | 13 | 3.20E-01 | 3.40E-01 | 7.50E-01 | 6.40E-01 | 5.80E-01 | 5.50E-01 | 6.50E-01 | 6.70E-01 | 3.80E-01 | 7.40E-01 | 5.80E-01 |
| SUCLA2 | rs1578303 | 3513335 | 13 | 1.30E-02 | 5.00E-01 | 5.60E-01 | 2.00E-01 | 1.50E-02 | 1.80E-01 | 7.20E-01 | 1.80E-01 | 2.90E-01 | 8.00E-01 | 3.50E-01 |
| SUGT1L1 | rs17446243 | 3510968 | 13 | 1.90E-03 | 3.60E-01 | 9.40E-01 | 1.10E-01 | 7.00E-01 | 7.90E-01 | 2.00E-01 | 5.40E-02 | 4.60E-01 | 4.10E-02 | 7.40E-01 |
| SUGT1L1 | rs17446278 | 3510968 | 13 | 1.90E-03 | 3.60E-01 | 9.40E-01 | 1.10E-01 | 7.00E-01 | 7.80E-01 | 2.00E-01 | 5.40E-02 | 4.60E-01 | 4.10E-02 | 7.40E-01 |
| SUGT1L1 | rs9532508 | 3510968 | 13 | 1.40E-03 | 4.10E-01 | 9.00E-01 | 8.70E-02 | 6.40E-01 | 8.20E-01 | 1.70E-01 | 5.80E-02 | 4.20E-01 | 3.60E-02 | 7.20E-01 |
| AP4S1 | rs2378971 | 3531231 | 14 | 1.50E-02 | 5.60E-01 | 6.60E-01 | 4.30E-02 | 8.40E-01 | 3.90E-01 | 7.00E-01 | 2.40E-02 | 5.00E-01 | 1.00E+00 | 2.10E-01 |
| AP4S1 | rs12886280 | 3531251 | 14 | 5.30E-03 | 8.30E-01 | 8.40E-01 | 3.20E-01 | 9.50E-01 | 3.80E-01 | 8.40E-02 | 5.40E-01 | 6.40E-01 | 4.40E-02 | 8.90E-02 |

| | | | | | | | | | | | | | | |
|---|---|---|---|---|---|---|---|---|---|---|---|---|---|---|
| G2E3 | rs10139154 | 3530983 | 14 | 9.90E-03 | 9.20E-01 | 5.80E-01 | 4.20E-01 | 1.40E-01 | 1.20E-01 | 3.30E-02 | 5.70E-01 | 1.50E-01 | 7.00E-02 | 1.80E-01 |
| G2E3 | rs447853 | 3531004 | 14 | 5.40E-03 | 5.80E-01 | 9.80E-02 | 7.10E-01 | 1.20E-01 | 6.30E-03 | 7.70E-02 | 2.90E-01 | 6.60E-01 | 4.90E-01 | 5.70E-01 |
| SCFD1 | rs151123 | 3531039 | 14 | 9.90E-03 | 1.40E-02 | 5.00E-01 | 2.40E-02 | 5.90E-01 | 2.80E-01 | 6.50E-02 | 2.70E-01 | 4.40E-01 | 1.60E-04 | 3.50E-02 |
| SCFD1 | rs447614 | 3531039 | 14 | 9.80E-03 | 1.30E-02 | 4.90E-01 | 2.60E-02 | 5.60E-01 | 3.00E-01 | 6.30E-02 | 2.60E-01 | 4.40E-01 | 1.60E-04 | 3.00E-02 |
| C15orf40 | rs6603044 | 3636468 | 15 | 7.20E-06 | 1.40E-04 | 1.70E-01 | 1.20E-01 | 3.40E-01 | 2.50E-03 | 3.00E-01 | 2.10E-02 | 4.80E-02 | 7.70E-01 | 4.10E-01 |
| WHAMM | rs2381030 | 3605193 | 15 | 9.80E-06 | 1.70E-05 | 4.10E-04 | 4.60E-03 | 6.30E-06 | 1.80E-02 | 4.00E-03 | 4.80E-02 | 1.40E-01 | 2.20E-02 | 8.50E-02 |
| WHAMM | rs1877240 | 3605193 | 15 | 1.70E-04 | 3.80E-04 | 2.70E-04 | 3.10E-02 | 2.00E-04 | 3.90E-02 | 3.10E-01 | 4.80E-01 | 1.90E-02 | 6.30E-02 | 1.80E-02 |
| WHAMM | rs13329195 | 3605193 | 15 | 7.40E-06 | 1.40E-05 | 3.80E-04 | 3.70E-03 | 4.20E-06 | 1.60E-02 | 4.40E-03 | 5.30E-02 | 1.20E-01 | 1.80E-02 | 6.90E-02 |
| WHAMM | rs12900374 | 3605193 | 15 | 1.70E-04 | 3.80E-04 | 2.70E-04 | 3.10E-02 | 2.10E-04 | 3.90E-02 | 3.10E-01 | 4.80E-01 | 1.90E-02 | 6.30E-02 | 1.80E-02 |
| WHAMML1 | rs2381030 | 3605193 | 15 | 9.80E-06 | 1.70E-05 | 4.10E-04 | 4.60E-03 | 6.30E-06 | 1.80E-02 | 4.00E-03 | 4.80E-02 | 1.40E-01 | 2.20E-02 | 8.50E-02 |
| WHAMML2 | rs2381030 | 3605193 | 15 | 9.80E-06 | 1.70E-05 | 4.10E-04 | 4.60E-03 | 6.30E-06 | 1.80E-02 | 4.00E-03 | 4.80E-02 | 1.40E-01 | 2.20E-02 | 8.50E-02 |
| WHAMM | rs2381030 | 3605193 | 15 | 9.80E-06 | 1.70E-05 | 4.10E-04 | 4.60E-03 | 6.30E-06 | 1.80E-02 | 4.00E-03 | 4.80E-02 | 1.40E-01 | 2.20E-02 | 8.50E-02 |
| WHAMML1 | rs1877240 | 3605193 | 15 | 1.70E-04 | 3.80E-04 | 2.70E-04 | 3.10E-02 | 2.00E-04 | 3.90E-02 | 3.10E-01 | 4.80E-01 | 1.90E-02 | 6.30E-02 | 1.80E-02 |
| WHAMML2 | rs1877240 | 3605193 | 15 | 1.70E-04 | 3.80E-04 | 2.70E-04 | 3.10E-02 | 2.00E-04 | 3.90E-02 | 3.10E-01 | 4.80E-01 | 1.90E-02 | 6.30E-02 | 1.80E-02 |
| WHAMM | rs1877240 | 3605193 | 15 | 1.70E-04 | 3.80E-04 | 2.70E-04 | 3.10E-02 | 2.00E-04 | 3.90E-02 | 3.10E-01 | 4.80E-01 | 1.90E-02 | 6.30E-02 | 1.80E-02 |
| WHAMML1 | rs12900374 | 3605193 | 15 | 1.70E-04 | 3.80E-04 | 2.70E-04 | 3.10E-02 | 2.10E-04 | 3.90E-02 | 3.10E-01 | 4.80E-01 | 1.90E-02 | 6.30E-02 | 1.80E-02 |
| WHAMML2 | rs12900374 | 3605193 | 15 | 1.70E-04 | 3.80E-04 | 2.70E-04 | 3.10E-02 | 2.10E-04 | 3.90E-02 | 3.10E-01 | 4.80E-01 | 1.90E-02 | 6.30E-02 | 1.80E-02 |
| WHAMM | rs12900374 | 3605193 | 15 | 1.70E-04 | 3.80E-04 | 2.70E-04 | 3.10E-02 | 2.10E-04 | 3.90E-02 | 3.10E-01 | 4.80E-01 | 1.90E-02 | 6.30E-02 | 1.80E-02 |
| WHAMML1 | rs13329195 | 3605193 | 15 | 7.40E-06 | 1.40E-05 | 3.80E-04 | 3.70E-03 | 4.20E-06 | 1.60E-02 | 4.40E-03 | 5.30E-02 | 1.20E-01 | 1.80E-02 | 6.90E-02 |
| WHAMML2 | rs13329195 | 3605193 | 15 | 7.40E-06 | 1.40E-05 | 3.80E-04 | 3.70E-03 | 4.20E-06 | 1.60E-02 | 4.40E-03 | 5.30E-02 | 1.20E-01 | 1.80E-02 | 6.90E-02 |
| WHAMM | rs13329195 | 3605193 | 15 | 7.40E-06 | 1.40E-05 | 3.80E-04 | 3.70E-03 | 4.20E-06 | 1.60E-02 | 4.40E-03 | 5.30E-02 | 1.20E-01 | 1.80E-02 | 6.90E-02 |
| WHAMML1 | rs13329195 | 3605193 | 15 | 7.40E-06 | 1.40E-05 | 3.80E-04 | 3.70E-03 | 4.20E-06 | 1.60E-02 | 4.40E-03 | 5.30E-02 | 1.20E-01 | 1.80E-02 | 6.90E-02 |
| WHAMML1 | rs12900374 | 3605193 | 15 | 1.70E-04 | 3.80E-04 | 2.70E-04 | 3.10E-02 | 2.10E-04 | 3.90E-02 | 3.10E-01 | 4.80E-01 | 1.90E-02 | 6.30E-02 | 1.80E-02 |
| WHAMML1 | rs2381030 | 3605193 | 15 | 9.80E-06 | 1.70E-05 | 4.10E-04 | 4.60E-03 | 6.30E-06 | 1.80E-02 | 4.00E-03 | 4.80E-02 | 1.40E-01 | 2.20E-02 | 8.50E-02 |
| WHAMML1 | rs1877240 | 3605193 | 15 | 1.70E-04 | 3.80E-04 | 2.70E-04 | 3.10E-02 | 2.00E-04 | 3.90E-02 | 3.10E-01 | 4.80E-01 | 1.90E-02 | 6.30E-02 | 1.80E-02 |
| WHAMML2 | rs1877240 | 3605193 | 15 | 1.70E-04 | 3.80E-04 | 2.70E-04 | 3.10E-02 | 2.00E-04 | 3.90E-02 | 3.10E-01 | 4.80E-01 | 1.90E-02 | 6.30E-02 | 1.80E-02 |
| WHAMML2 | rs13329195 | 3605193 | 15 | 7.40E-06 | 1.40E-05 | 3.80E-04 | 3.70E-03 | 4.20E-06 | 1.60E-02 | 4.40E-03 | 5.30E-02 | 1.20E-01 | 1.80E-02 | 6.90E-02 |
| WHAMML2 | rs12900374 | 3605193 | 15 | 1.70E-04 | 3.80E-04 | 2.70E-04 | 3.10E-02 | 2.10E-04 | 3.90E-02 | 3.10E-01 | 4.80E-01 | 1.90E-02 | 6.30E-02 | 1.80E-02 |
| WHAMML2 | rs2381030 | 3605193 | 15 | 9.80E-06 | 1.70E-05 | 4.10E-04 | 4.60E-03 | 6.30E-06 | 1.80E-02 | 4.00E-03 | 4.80E-02 | 1.40E-01 | 2.20E-02 | 8.50E-02 |

| | | | | | | | | | | | | | | |
|---|---|---|---|---|---|---|---|---|---|---|---|---|---|---|
| SYCE1L | rs56024498 | 3669418 | 16 | 1.20E-04 | 1.00E-03 | 7.60E-01 | 4.50E-01 | 4.10E-01 | 3.10E-01 | 1.40E-02 | 1.90E-01 | 2.60E-01 | 7.10E-01 | 1.90E-01 |
| LOC100130958 | rs56024498 | 3669418 | 16 | 1.20E-04 | 1.00E-03 | 7.60E-01 | 4.50E-01 | 4.10E-01 | 3.10E-01 | 1.40E-02 | 1.90E-01 | 2.60E-01 | 7.10E-01 | 1.90E-01 |
| CUEDC1 | rs2240601 | 3764043 | 17 | 3.00E-03 | 8.80E-01 | 1.50E-02 | 6.90E-01 | 9.30E-01 | 3.90E-03 | 1.30E-02 | 2.60E-01 | 4.60E-01 | 7.40E-01 | 5.20E-01 |
| GGNBP2 | rs2285640 | 3719164 | 17 | 7.60E-12 | 1.80E-04 | 2.20E-02 | 7.70E-07 | 2.80E-03 | 1.80E-01 | 2.50E-02 | 1.10E-01 | 8.50E-04 | 8.70E-03 | 2.80E-01 |
| GGNBP2 | rs11263770 | 3719164 | 17 | 7.80E-12 | 3.60E-05 | 2.10E-02 | 7.50E-07 | 3.50E-03 | 1.80E-01 | 7.10E-02 | 9.80E-02 | 1.00E-03 | 8.50E-03 | 2.50E-01 |
| GGNBP2 | rs9906189 | 3719164 | 17 | 7.20E-12 | 2.20E-04 | 2.10E-02 | 8.00E-07 | 2.70E-03 | 1.70E-01 | 2.70E-02 | 9.60E-02 | 1.10E-03 | 7.20E-03 | 2.70E-01 |
| GGNBP2 | rs9903355 | 3719164 | 17 | 8.40E-12 | 2.00E-04 | 2.20E-02 | 7.90E-07 | 3.00E-03 | 1.80E-01 | 2.50E-02 | 1.00E-01 | 9.90E-04 | 8.00E-03 | 2.80E-01 |
| LOC100294337 | rs7224296 | 3760211 | 17 | 3.30E-18 | 6.30E-10 | 1.80E-11 | 3.40E-13 | 3.80E-13 | 5.30E-09 | 8.00E-07 | 3.50E-07 | 1.00E-11 | 6.60E-14 | 1.60E-15 |
| KIAA1267 | rs7224296 | 3760211 | 17 | 3.30E-18 | 6.30E-10 | 1.80E-11 | 3.40E-13 | 3.80E-13 | 5.30E-09 | 8.00E-07 | 3.50E-07 | 1.00E-11 | 6.60E-14 | 1.60E-15 |
| LOC100130950 | rs7209200 | 3707601 | 17 | 2.70E-08 | 5.90E-03 | 6.90E-03 | 1.10E-02 | 1.60E-02 | 6.80E-06 | 1.60E-03 | 1.60E-02 | 1.10E-03 | 3.60E-02 | 1.30E-03 |
| TNFAIP1 | rs739439 | 3750631 | 17 | 5.00E-04 | 6.00E-01 | 9.50E-03 | 4.40E-02 | 2.40E-02 | 2.60E-03 | 9.20E-02 | 2.50E-02 | 3.80E-01 | 3.90E-01 | 6.80E-01 |
| POLDIP2 | rs739439 | 3750631 | 17 | 5.00E-04 | 6.00E-01 | 9.50E-03 | 4.40E-02 | 2.40E-02 | 2.60E-03 | 9.20E-02 | 2.50E-02 | 3.80E-01 | 3.90E-01 | 6.80E-01 |
| TNFAIP1 | rs34660379 | 3750631 | 17 | 5.60E-05 | 9.30E-01 | 6.40E-03 | 1.30E-03 | 4.20E-02 | 8.80E-05 | 5.70E-02 | 4.60E-03 | 4.80E-01 | 8.40E-01 | 3.00E-01 |
| POLDIP2 | rs34660379 | 3750631 | 17 | 5.60E-05 | 9.30E-01 | 6.40E-03 | 1.30E-03 | 4.20E-02 | 8.80E-05 | 5.70E-02 | 4.60E-03 | 4.80E-01 | 8.40E-01 | 3.00E-01 |
| TNFAIP1 | rs35714695 | 3750631 | 17 | 9.10E-05 | 6.60E-01 | 8.10E-03 | 2.00E-03 | 3.00E-02 | 2.50E-04 | 8.20E-02 | 1.10E-02 | 3.90E-01 | 5.30E-01 | 5.10E-01 |
| POLDIP2 | rs35714695 | 3750631 | 17 | 9.10E-05 | 6.60E-01 | 8.10E-03 | 2.00E-03 | 3.00E-02 | 2.50E-04 | 8.20E-02 | 1.10E-02 | 3.90E-01 | 5.30E-01 | 5.10E-01 |
| STX8 | rs9894834 | 3744806 | 17 | 5.90E-07 | 7.50E-02 | 1.30E-04 | 1.30E-01 | 6.10E-03 | 3.90E-04 | 1.40E-07 | 3.10E-01 | 1.60E-02 | 7.80E-06 | 1.50E-04 |
| DOK6 | rs17187386 | 3792875 | 18 | 3.80E-02 | 1.60E-01 | 9.20E-03 | 6.10E-01 | 7.80E-01 | 1.20E-01 | 1.20E-01 | 9.50E-01 | 4.70E-01 | 3.60E-01 | 3.40E-01 |
| SLMO1 | rs12967284 | 3779625 | 18 | 3.60E-04 | 1.40E-02 | 1.20E-01 | 2.30E-01 | 7.90E-02 | 7.30E-01 | 3.60E-01 | 6.10E-01 | 2.30E-02 | 3.70E-01 | 3.70E-02 |
| HMGB3 | rs12967284 | 3779625 | 18 | 3.60E-04 | 1.40E-02 | 1.20E-01 | 2.30E-01 | 7.90E-02 | 7.30E-01 | 3.60E-01 | 6.10E-01 | 2.30E-02 | 3.70E-01 | 3.70E-02 |
| AP3D1 | rs7258235 | 3845711 | 19 | 1.30E-03 | 4.90E-02 | 3.10E-01 | 4.30E-01 | 9.70E-03 | 8.70E-02 | 2.70E-01 | 3.00E-01 | 3.00E-01 | 6.00E-02 | 3.10E-01 |
| C19orf57 | rs112183647 | 3852360 | 19 | 2.80E-04 | 3.80E-01 | 4.30E-02 | 7.30E-01 | 7.00E-01 | 4.10E-01 | 1.10E-01 | 8.60E-01 | 3.30E-02 | 5.70E-01 | 3.30E-01 |
| HAUS8 | rs78549703 | 3854230 | 19 | 2.40E-03 | 3.70E-02 | 3.20E-01 | 4.00E-01 | 5.60E-01 | 6.30E-01 | 3.60E-01 | 1.20E-02 | 9.20E-01 | 1.20E-01 | 1.40E-02 |
| HAUS8 | rs12608932 | 3854230 | 19 | 1.80E-03 | 2.70E-02 | 3.20E-01 | 4.50E-01 | 5.50E-01 | 5.60E-01 | 3.90E-01 | 9.40E-03 | 8.90E-01 | 9.70E-02 | 9.40E-03 |
| C20orf24 | rs2425220 | 3883980 | 20 | 1.20E-04 | 7.50E-01 | 1.50E-02 | 2.50E-01 | 3.40E-02 | 8.90E-02 | 9.40E-01 | 8.20E-01 | 4.90E-02 | 6.20E-03 | 4.10E-01 |
| PPP4R1L | rs6015322 | 3911396 | 20 | 1.20E-03 | 7.00E-01 | 5.20E-02 | 3.10E-02 | 2.80E-01 | 7.00E-02 | 6.40E-01 | 7.20E-01 | 7.00E-01 | 3.60E-01 | 6.30E-02 |
| PRSS7 | rs9653747 | 3926306 | 21 | 5.50E-02 | 6.60E-01 | 3.80E-01 | 9.20E-01 | 5.30E-01 | 5.70E-01 | 1.40E-01 | 4.50E-03 | 2.90E-01 | 1.10E-01 | 7.50E-01 |
| TMPRSS15 | rs9653747 | 3926306 | 21 | 5.50E-02 | 6.60E-01 | 3.80E-01 | 9.20E-01 | 5.30E-01 | 5.70E-01 | 1.40E-01 | 4.50E-03 | 2.90E-01 | 1.10E-01 | 7.50E-01 |
| NUP50 | rs2176039 | 3948484 | 22 | 7.40E-05 | 8.00E-02 | 8.80E-01 | 6.30E-02 | 1.50E-02 | 3.20E-01 | 5.50E-01 | 3.20E-01 | 4.20E-04 | 6.20E-01 | 1.60E-01 |

Table S3. Gene network showing genetic interactions, physical interactions, pathways, co-expression, co-localization and protein domain similarity associated with the ALS genetic risk genes

| Gene 1 | Gene 2 | Weight | Network group | Network |
|---|---|---|---|---|
| KIAA0524 | GABRG3 | 0.007286813 | Co-expression | Dobbin-Giordano-2005 |
| CPNE7 | PITPNC1 | 0.005566633 | Co-expression | Dobbin-Giordano-2005 |
| DCTN4 | NUBPL | 0.01384219 | Co-expression | Dobbin-Giordano-2005 |
| DLGAP4 | CPNE7 | 0.016254801 | Co-expression | Dobbin-Giordano-2005 |
| G6PC2 | TSPAN32 | 0.003912206 | Co-expression | Dobbin-Giordano-2005 |
| AP3D1 | TNIP1 | 0.017540012 | Co-expression | Dobbin-Giordano-2005 |
| KIF5A | SYCE1L | 0.01251856 | Co-expression | Dobbin-Giordano-2005 |
| KIF5A | G6PC2 | 0.003414679 | Co-expression | Dobbin-Giordano-2005 |
| HSD17B6 | AKR1D1 | 0.010065821 | Co-expression | Dobbin-Giordano-2005 |
| ADH1A | HSD17B6 | 0.004890242 | Co-expression | Dobbin-Giordano-2005 |
| GNG7 | SCN5A | 0.006443857 | Co-expression | Dobbin-Giordano-2005 |
| GNG7 | NFASC | 0.005163134 | Co-expression | Dobbin-Giordano-2005 |
| SAR1B | AKR1D1 | 0.011008999 | Co-expression | Dobbin-Giordano-2005 |
| parkin | G6PC2 | 0.005404248 | Co-expression | Dobbin-Giordano-2005 |
| CAT | ACADS | 0.011595692 | Co-expression | Dobbin-Giordano-2005 |
| KIAA0917 | NUBPL | 0.013109388 | Co-expression | Dobbin-Giordano-2005 |
| MAPK9 | HMGB3 | 0.011988649 | Co-expression | Dobbin-Giordano-2005 |
| SMARCA2 | ELMO1 | 0.016750768 | Co-expression | Dobbin-Giordano-2005 |
| GLI3 | KIAA0524 | 0.019991884 | Co-expression | Dobbin-Giordano-2005 |
| GLI3 | NFASC | 0.009700749 | Co-expression | Dobbin-Giordano-2005 |
| GLI3 | COL16A1 | 0.010357793 | Co-expression | Dobbin-Giordano-2005 |
| A2BP1 | UNC13A | 0.012120464 | Co-expression | Dobbin-Giordano-2005 |
| A2BP1 | SCN5A | 0.009124341 | Co-expression | Dobbin-Giordano-2005 |
| A2BP1 | NFASC | 0.007936502 | Co-expression | Dobbin-Giordano-2005 |
| STX8 | NUBPL | 0.013928944 | Co-expression | Dobbin-Giordano-2005 |
| STX8 | KIAA0917 | 0.012243342 | Co-expression | Dobbin-Giordano-2005 |
| DOCK2 | ELMO1 | 0.0079722 | Co-expression | Dobbin-Giordano-2005 |
| DOCK2 | SMARCA2 | 0.008537297 | Co-expression | Dobbin-Giordano-2005 |
| STX18 | DCTN4 | 0.01687169 | Co-expression | Dobbin-Giordano-2005 |
| STX12 | TXNDC15 | 0.022064833 | Co-expression | Dobbin-Giordano-2005 |
| PFKP | PITPNC1 | 0.006107026 | Co-expression | Dobbin-Giordano-2005 |
| TDRD7 | NTN1 | 0.007828274 | Co-expression | Dobbin-Giordano-2005 |

| | | | | |
|---|---|---|---|---|
| STX5 | GABRG3 | 0.006993514 | Co-expression | Dobbin-Giordano-2005 |
| ARSB | G6PC2 | 0.005757078 | Co-expression | Dobbin-Giordano-2005 |
| G2E3 | WHAMM | 0.018878302 | Co-expression | Perou-Botstein-2000 |
| ASB1 | BIRC6 | 0.013104028 | Co-expression | Perou-Botstein-2000 |
| SH3BP2 | MOBP | 0.022695266 | Co-expression | Perou-Botstein-2000 |
| COL16A1 | KIAA0524 | 0.005582513 | Co-expression | Perou-Botstein-2000 |
| COL16A1 | HSD17B6 | 0.005594579 | Co-expression | Perou-Botstein-2000 |
| GIPC1 | HTR2A | 0.01107967 | Co-expression | Perou-Botstein-2000 |
| SMARCA2 | HSPBAP1 | 0.010614253 | Co-expression | Perou-Botstein-2000 |
| ATXN2 | NDST1 | 0.016952286 | Co-expression | Perou-Botstein-2000 |
| ATXN2 | UGCG | 0.018782971 | Co-expression | Perou-Botstein-2000 |
| NSF | SAR1B | 0.013871493 | Co-expression | Perou-Botstein-2000 |
| CIC | SETD2 | 0.014090467 | Co-expression | Perou-Botstein-2000 |
| DOCK2 | ELMO1 | 0.013441643 | Co-expression | Perou-Botstein-2000 |
| STX5 | TNPO3 | 0.015387598 | Co-expression | Perou-Botstein-2000 |
| STX5 | WIPI2 | 0.021385381 | Co-expression | Perou-Botstein-2000 |
| C15orf40 | ZFP3 | 0.0225099 | Co-expression | Noble-Diehl-2008 |
| C21orf2 | WHAMM | 0.006715288 | Co-expression | Noble-Diehl-2008 |
| DHRS11 | ABLIM2 | 0.005199276 | Co-expression | Noble-Diehl-2008 |
| BIRC6 | ZFP3 | 0.008724484 | Co-expression | Noble-Diehl-2008 |
| MOBKL2B | DHRS11 | 0.007618249 | Co-expression | Noble-Diehl-2008 |
| BTBD1 | DCTN4 | 0.007831062 | Co-expression | Noble-Diehl-2008 |
| TNPO3 | BIRC6 | 0.008434472 | Co-expression | Noble-Diehl-2008 |
| CYR61 | APOLD1 | 0.026415637 | Co-expression | Noble-Diehl-2008 |
| NUP50 | C20orf24 | 0.0088147 | Co-expression | Noble-Diehl-2008 |
| UGCG | TBK1 | 0.017199613 | Co-expression | Noble-Diehl-2008 |
| WIPI2 | ABLIM2 | 0.014663695 | Co-expression | Noble-Diehl-2008 |
| ELMO1 | TSPAN32 | 0.034189574 | Co-expression | Noble-Diehl-2008 |
| ELMO1 | HTR2A | 0.022509124 | Co-expression | Noble-Diehl-2008 |
| ELMO1 | TG | 0.010006554 | Co-expression | Noble-Diehl-2008 |
| KIAA0917 | TNFAIP1 | 0.008729959 | Co-expression | Noble-Diehl-2008 |
| GIPC1 | ZFYVE28 | 0.007181022 | Co-expression | Noble-Diehl-2008 |
| MAPK9 | BIRC6 | 0.004630518 | Co-expression | Noble-Diehl-2008 |
| TEK | TXNDC15 | 0.005144256 | Co-expression | Noble-Diehl-2008 |
| GLI3 | HSD17B6 | 0.004288347 | Co-expression | Noble-Diehl-2008 |
| GLI3 | COL16A1 | 0.004755667 | Co-expression | Noble-Diehl-2008 |
| GLI3 | PCDH9 | 0.007106588 | Co-expression | Noble-Diehl-2008 |
| ATXN2 | WHAMM | 0.006358025 | Co-expression | Noble-Diehl-2008 |

| | | | | |
|---|---|---|---|---|
| STX8 | DCTN4 | 0.008762078 | Co-expression | Noble-Diehl-2008 |
| STX8 | CAT | 0.02213303 | Co-expression | Noble-Diehl-2008 |
| DNM2 | AHCYL2 | 0.004029511 | Co-expression | Noble-Diehl-2008 |
| DNM2 | ZFYVE28 | 0.004743413 | Co-expression | Noble-Diehl-2008 |
| DNM2 | ACADS | 0.004011209 | Co-expression | Noble-Diehl-2008 |
| DNM2 | GIPC1 | 0.002682871 | Co-expression | Noble-Diehl-2008 |
| NUDC | ASB1 | 0.009698252 | Co-expression | Noble-Diehl-2008 |
| TDRD7 | MOBKL2B | 0.010720439 | Co-expression | Noble-Diehl-2008 |
| NAPG | C20orf24 | 0.010859227 | Co-expression | Noble-Diehl-2008 |
| IDH3B | C19orf57 | 0.008553861 | Co-expression | Noble-Diehl-2008 |
| IDH3B | NUBPL | 0.009272989 | Co-expression | Noble-Diehl-2008 |
| PAX8 | IL37 | 0.011846123 | Co-expression | Rosenwald-Staudt-2001 |
| SETD2 | BIRC6 | 0.01396872 | Co-expression | Rosenwald-Staudt-2001 |
| SETD2 | RNF3A | 0.010957726 | Co-expression | Rosenwald-Staudt-2001 |
| SMARCA2 | BNIP1 | 0.018510392 | Co-expression | Rosenwald-Staudt-2001 |
| A2BP1 | MOBKL2B | 0.021273388 | Co-expression | Rosenwald-Staudt-2001 |
| BNIP1 | IL37 | 0.011099311 | Co-expression | Boldrick-Relman-2002 |
| PAX8 | IL37 | 0.011854169 | Co-expression | Boldrick-Relman-2002 |
| UGCG | CYR61 | 0.02464532 | Co-expression | Boldrick-Relman-2002 |
| MCC | SH3BP2 | 0.017953536 | Co-expression | Boldrick-Relman-2002 |
| DOCK2 | TSPAN32 | 0.011124335 | Co-expression | Boldrick-Relman-2002 |
| DOCK2 | SMARCA2 | 0.00803542 | Co-expression | Boldrick-Relman-2002 |
| DOCK2 | ERGIC3 | 0.011830392 | Co-expression | Boldrick-Relman-2002 |
| HSD17B6 | AKR1D1 | 0.011326293 | Co-expression | Wu-Garvey-2007 |
| NFASC | KIAA0524 | 0.016778303 | Co-expression | Wu-Garvey-2007 |
| TG | MOBP | 0.002599769 | Co-expression | Wu-Garvey-2007 |
| TG | HTR2A | 0.006554249 | Co-expression | Wu-Garvey-2007 |
| SETD2 | UGCG | 0.016692262 | Co-expression | Wu-Garvey-2007 |
| GIPC1 | NDST1 | 0.008794934 | Co-expression | Wu-Garvey-2007 |
| GIPC1 | TNFAIP1 | 0.012098746 | Co-expression | Wu-Garvey-2007 |
| GIPC1 | AP3D1 | 0.010529459 | Co-expression | Wu-Garvey-2007 |
| SMARCA2 | KIAA0917 | 0.007989998 | Co-expression | Wu-Garvey-2007 |
| ATXN2 | ASB1 | 0.022113938 | Co-expression | Wu-Garvey-2007 |
| NUDC | TNIP1 | 0.013101539 | Co-expression | Wu-Garvey-2007 |
| ERGIC3 | GIPC1 | 0.009182973 | Co-expression | Wu-Garvey-2007 |
| STX12 | UGCG | 0.013601212 | Co-expression | Wu-Garvey-2007 |
| STX12 | SETD2 | 0.010989543 | Co-expression | Wu-Garvey-2007 |
| STX7 | SETD2 | 0.014803617 | Co-expression | Wu-Garvey-2007 |

| | | | | |
|---|---|---|---|---|
| TNIP1 | PITPNC1 | 0.011393133 | Co-expression | Wang-Cheung-2015 |
| TNPO3 | DLGAP4 | 0.01952188 | Co-expression | Wang-Cheung-2015 |
| CYR61 | RNF3A | 0.016282717 | Co-expression | Wang-Cheung-2015 |
| GNG7 | ADH1A | 0.018714052 | Co-expression | Wang-Cheung-2015 |
| COL16A1 | BNIP1 | 0.006059661 | Co-expression | Wang-Cheung-2015 |
| UGCG | CYR61 | 0.007422074 | Co-expression | Wang-Cheung-2015 |
| NTN1 | CNTN6 | 0.013292592 | Co-expression | Wang-Cheung-2015 |
| MCC | DCTN4 | 0.01830397 | Co-expression | Wang-Cheung-2015 |
| WIPI2 | UGCG | 0.011388299 | Co-expression | Wang-Cheung-2015 |
| KIAA0917 | COL16A1 | 0.006676947 | Co-expression | Wang-Cheung-2015 |
| ATXN2 | GGNBP2 | 0.008713255 | Co-expression | Wang-Cheung-2015 |
| ATXN2 | WIPI2 | 0.013816046 | Co-expression | Wang-Cheung-2015 |
| DOCK2 | MSRA | 0.011054218 | Co-expression | Wang-Cheung-2015 |
| STX18 | PITPNC1 | 0.01405296 | Co-expression | Wang-Cheung-2015 |
| PFKP | HTR2A | 0.010799343 | Co-expression | Wang-Cheung-2015 |
| NAPG | UGCG | 0.009980558 | Co-expression | Wang-Cheung-2015 |
| NUP50 | G2E3 | 0.006732316 | Co-expression | Alizadeh-Staudt-2000 |
| GPX3 | CYR61 | 0.01808541 | Co-expression | Alizadeh-Staudt-2000 |
| PCDH9 | GPX3 | 0.010569473 | Co-expression | Alizadeh-Staudt-2000 |
| SMARCA2 | BNIP1 | 0.015188683 | Co-expression | Alizadeh-Staudt-2000 |
| TNIP1 | ASB1 | 0.016683621 | Co-expression | Bild-Nevins-2006 B |
| NFASC | KIAA0524 | 0.02477285 | Co-expression | Bild-Nevins-2006 B |
| TNPO3 | DLGAP4 | 0.015736448 | Co-expression | Bild-Nevins-2006 B |
| GPX3 | ADH1A | 0.019506322 | Co-expression | Bild-Nevins-2006 B |
| SETD2 | RNF3A | 0.023322186 | Co-expression | Bild-Nevins-2006 B |
| ELMO1 | GPX3 | 0.009601185 | Co-expression | Bild-Nevins-2006 B |
| SMARCA2 | CAT | 0.006690039 | Co-expression | Bild-Nevins-2006 B |
| GOSR1 | TNFAIP1 | 0.015056666 | Co-expression | Bild-Nevins-2006 B |
| ERGIC3 | DLGAP4 | 0.011421819 | Co-expression | Bild-Nevins-2006 B |
| CIC | DLGAP4 | 0.014602494 | Co-expression | Bild-Nevins-2006 B |
| CIC | TNIP1 | 0.012275652 | Co-expression | Bild-Nevins-2006 B |
| DOCK2 | ELMO1 | 0.005410322 | Co-expression | Bild-Nevins-2006 B |
| PFKP | VTI1B | 0.012091057 | Co-expression | Bild-Nevins-2006 B |
| STX5 | TNIP1 | 0.012263699 | Co-expression | Bild-Nevins-2006 B |
| NAPG | GIPC1 | 0.013254213 | Co-expression | Bild-Nevins-2006 B |
| GPX3 | ADH1A | 0.005384604 | Co-localization | Schadt-Shoemaker-2004 |
| GPX3 | CYR61 | 0.005575808 | Co-localization | Schadt-Shoemaker-2004 |
| CAT | ADH1A | 0.005646279 | Co-localization | Schadt-Shoemaker-2004 |

| | | | | |
|---|---|---|---|---|
| CAT | ACADS | 0.008117259 | Co-localization | Schadt-Shoemaker-2004 |
| TEK | ADH1A | 0.00496935 | Co-localization | Schadt-Shoemaker-2004 |
| TEK | CYR61 | 0.004971826 | Co-localization | Schadt-Shoemaker-2004 |
| TEK | GPX3 | 0.005854193 | Co-localization | Schadt-Shoemaker-2004 |
| ARSB | GDNF | 0.034320004 | Co-localization | Schadt-Shoemaker-2004 |
| WIBG | SLC10A7 | 0.016242802 | Co-localization | Johnson-Shoemaker-2003 |
| TNFAIP1 | CRIM1 | 0.006437785 | Co-localization | Johnson-Shoemaker-2003 |
| HSPBAP1 | CRIM1 | 0.005716966 | Co-localization | Johnson-Shoemaker-2003 |
| HSD17B6 | C19orf57 | 0.008758523 | Co-localization | Johnson-Shoemaker-2003 |
| HSD17B6 | AKR1D1 | 0.004655154 | Co-localization | Johnson-Shoemaker-2003 |
| HMGB3 | CRIM1 | 0.006520567 | Co-localization | Johnson-Shoemaker-2003 |
| ADH1A | AKR1D1 | 0.006630094 | Co-localization | Johnson-Shoemaker-2003 |
| ADH1A | HSD17B6 | 0.006267184 | Co-localization | Johnson-Shoemaker-2003 |
| HTR2A | KIF5A | 0.00666198 | Co-localization | Johnson-Shoemaker-2003 |
| TG | GPX3 | 0.007412243 | Co-localization | Johnson-Shoemaker-2003 |
| PAX8 | GPX3 | 0.011734744 | Co-localization | Johnson-Shoemaker-2003 |
| PAX8 | TG | 0.011348739 | Co-localization | Johnson-Shoemaker-2003 |
| CAT | AKR1D1 | 0.007142729 | Co-localization | Johnson-Shoemaker-2003 |
| TEK | TNFAIP1 | 0.008490931 | Co-localization | Johnson-Shoemaker-2003 |
| TEK | CYR61 | 0.006214554 | Co-localization | Johnson-Shoemaker-2003 |
| GLI3 | KIAA0524 | 0.009511952 | Co-localization | Johnson-Shoemaker-2003 |
| ATXN2 | DCTN4 | 0.024206327 | Co-localization | Johnson-Shoemaker-2003 |
| NSF | KIF5A | 0.003043416 | Co-localization | Johnson-Shoemaker-2003 |
| NSF | HTR2A | 0.003519136 | Co-localization | Johnson-Shoemaker-2003 |
| NSF | MAPK9 | 0.004453508 | Co-localization | Johnson-Shoemaker-2003 |
| A2BP1 | KIF5A | 0.004558156 | Co-localization | Johnson-Shoemaker-2003 |
| A2BP1 | HTR2A | 0.005286318 | Co-localization | Johnson-Shoemaker-2003 |
| A2BP1 | NSF | 0.002447994 | Co-localization | Johnson-Shoemaker-2003 |
| DNM2 | MOBKL2B | 0.014613495 | Co-localization | Johnson-Shoemaker-2003 |
| NUDC | HSD17B6 | 0.009637226 | Co-localization | Johnson-Shoemaker-2003 |
| CNTNAP1 | NSF | 0.007018059 | Co-localization | Johnson-Shoemaker-2003 |
| VTI1B | MAPK9 | 0.023011085 | Co-localization | Johnson-Shoemaker-2003 |
| PFKP | HTR2A | 0.013975075 | Co-localization | Johnson-Shoemaker-2003 |
| STX7 | ELMO1 | 0.02195248 | Co-localization | Johnson-Shoemaker-2003 |
| NAPG | MAPK9 | 0.013341601 | Co-localization | Johnson-Shoemaker-2003 |
| NAPG | NSF | 0.003304744 | Co-localization | Johnson-Shoemaker-2003 |
| NAPG | A2BP1 | 0.004839824 | Co-localization | Johnson-Shoemaker-2003 |
| ARSB | MSRA | 0.015247721 | Co-localization | Johnson-Shoemaker-2003 |

| | | | | |
|---|---|---|---|---|
| SPIRE1 | SLC10A7 | 0.001985388 | Genetic Interactions | Lin-Smith-2010 |
| COL24A1 | C15orf40 | 0.001081622 | Genetic Interactions | Lin-Smith-2010 |
| ABLIM2 | COL24A1 | 0.000654556 | Genetic Interactions | Lin-Smith-2010 |
| ABLIM2 | C9orf72 | 0.000432864 | Genetic Interactions | Lin-Smith-2010 |
| CRIM1 | APCDD1L | 0.000631964 | Genetic Interactions | Lin-Smith-2010 |
| CRIM1 | SPIRE1 | 0.000393137 | Genetic Interactions | Lin-Smith-2010 |
| CRIM1 | C9orf72 | 0.000297491 | Genetic Interactions | Lin-Smith-2010 |
| CRIM1 | ABLIM2 | 0.000419709 | Genetic Interactions | Lin-Smith-2010 |
| MSI2 | SLC10A7 | 0.001602157 | Genetic Interactions | Lin-Smith-2010 |
| MSI2 | C9orf72 | 0.000327195 | Genetic Interactions | Lin-Smith-2010 |
| SLITRK1 | COL24A1 | 0.000689306 | Genetic Interactions | Lin-Smith-2010 |
| SLITRK1 | C9orf72 | 0.000455844 | Genetic Interactions | Lin-Smith-2010 |
| SLITRK1 | ZFYVE28 | 0.001143852 | Genetic Interactions | Lin-Smith-2010 |
| SLITRK1 | ABLIM2 | 0.000643118 | Genetic Interactions | Lin-Smith-2010 |
| SLITRK1 | CRIM1 | 0.000441991 | Genetic Interactions | Lin-Smith-2010 |
| C20orf24 | C15orf40 | 0.00102633 | Genetic Interactions | Lin-Smith-2010 |
| C20orf24 | C21orf2 | 0.001384699 | Genetic Interactions | Lin-Smith-2010 |
| C20orf24 | DOK6 | 0.000739284 | Genetic Interactions | Lin-Smith-2010 |
| C20orf24 | CRIM1 | 0.000426856 | Genetic Interactions | Lin-Smith-2010 |
| C20orf24 | MSI2 | 0.000469476 | Genetic Interactions | Lin-Smith-2010 |
| C6orf15 | CRIM1 | 0.001932729 | Genetic Interactions | Lin-Smith-2010 |
| NOSTRIN | ZFYVE28 | 0.001019181 | Genetic Interactions | Lin-Smith-2010 |
| MOBP | SLC10A7 | 0.003136379 | Genetic Interactions | Lin-Smith-2010 |
| MOBP | VARS2 | 0.001209641 | Genetic Interactions | Lin-Smith-2010 |
| NUBPL | C9orf72 | 0.000500781 | Genetic Interactions | Lin-Smith-2010 |
| NUBPL | ABLIM2 | 0.000706515 | Genetic Interactions | Lin-Smith-2010 |
| NUBPL | MSI2 | 0.000534044 | Genetic Interactions | Lin-Smith-2010 |
| NUBPL | NOSTRIN | 0.000662931 | Genetic Interactions | Lin-Smith-2010 |
| NUBPL | NDST1 | 0.001500394 | Genetic Interactions | Lin-Smith-2010 |
| MSRA | SPIRE1 | 0.000624837 | Genetic Interactions | Lin-Smith-2010 |
| MSRA | DOK6 | 0.000794006 | Genetic Interactions | Lin-Smith-2010 |
| MSRA | CRIM1 | 0.000458452 | Genetic Interactions | Lin-Smith-2010 |
| CPNE7 | C9orf72 | 0.002533326 | Genetic Interactions | Lin-Smith-2010 |
| IL37 | DOK6 | 0.00192961 | Genetic Interactions | Lin-Smith-2010 |
| AKR1D1 | SPIRE1 | 0.002974642 | Genetic Interactions | Lin-Smith-2010 |
| AKR1D1 | SLMO1 | 0.005909902 | Genetic Interactions | Lin-Smith-2010 |
| ELP3 | C15orf40 | 0.001554935 | Genetic Interactions | Lin-Smith-2010 |
| ELP3 | CPNE7 | 0.005507103 | Genetic Interactions | Lin-Smith-2010 |

| | | | | |
|---|---|---|---|---|
| RGS12 | SPIRE1 | 0.000707842 | Genetic Interactions | Lin-Smith-2010 |
| RGS12 | C9orf72 | 0.000535633 | Genetic Interactions | Lin-Smith-2010 |
| RGS12 | MSI2 | 0.00057121 | Genetic Interactions | Lin-Smith-2010 |
| RGS12 | NOSTRIN | 0.000709067 | Genetic Interactions | Lin-Smith-2010 |
| RGS12 | NUBPL | 0.000874252 | Genetic Interactions | Lin-Smith-2010 |
| TSPAN32 | VARS2 | 0.001170965 | Genetic Interactions | Lin-Smith-2010 |
| TSPAN32 | COL24A1 | 0.00093759 | Genetic Interactions | Lin-Smith-2010 |
| TSPAN32 | ABLIM2 | 0.000874765 | Genetic Interactions | Lin-Smith-2010 |
| TSPAN32 | SLITRK1 | 0.000921206 | Genetic Interactions | Lin-Smith-2010 |
| TSPAN32 | ELP3 | 0.001347876 | Genetic Interactions | Lin-Smith-2010 |
| SCN5A | C15orf40 | 0.001592784 | Genetic Interactions | Lin-Smith-2010 |
| SCN5A | SPIRE1 | 0.000902868 | Genetic Interactions | Lin-Smith-2010 |
| SCN5A | DOK6 | 0.001147312 | Genetic Interactions | Lin-Smith-2010 |
| SCN5A | MSRA | 0.001052868 | Genetic Interactions | Lin-Smith-2010 |
| G2E3 | APCDD1L | 0.000666918 | Genetic Interactions | Lin-Smith-2010 |
| G2E3 | C15orf40 | 0.000731909 | Genetic Interactions | Lin-Smith-2010 |
| G2E3 | ABLIM2 | 0.000442923 | Genetic Interactions | Lin-Smith-2010 |
| G2E3 | MSI2 | 0.000334799 | Genetic Interactions | Lin-Smith-2010 |
| G2E3 | NDST1 | 0.000940616 | Genetic Interactions | Lin-Smith-2010 |
| BIRC6 | MOBP | 0.001582983 | Genetic Interactions | Lin-Smith-2010 |
| BIRC6 | SCN5A | 0.001688499 | Genetic Interactions | Lin-Smith-2010 |
| TNFAIP1 | SPIRE1 | 0.004515764 | Genetic Interactions | Lin-Smith-2010 |
| ASB1 | C15orf40 | 0.000838291 | Genetic Interactions | Lin-Smith-2010 |
| ASB1 | VARS2 | 0.000679077 | Genetic Interactions | Lin-Smith-2010 |
| ASB1 | ABLIM2 | 0.000507302 | Genetic Interactions | Lin-Smith-2010 |
| ASB1 | MSI2 | 0.000383462 | Genetic Interactions | Lin-Smith-2010 |
| ASB1 | NUBPL | 0.000586898 | Genetic Interactions | Lin-Smith-2010 |
| ASB1 | TSPAN32 | 0.000726662 | Genetic Interactions | Lin-Smith-2010 |
| MOBKL2B | SPIRE1 | 0.00051983 | Genetic Interactions | Lin-Smith-2010 |
| MOBKL2B | ABLIM2 | 0.000554965 | Genetic Interactions | Lin-Smith-2010 |
| MOBKL2B | MSI2 | 0.000419489 | Genetic Interactions | Lin-Smith-2010 |
| MOBKL2B | TSPAN32 | 0.000794935 | Genetic Interactions | Lin-Smith-2010 |
| MOBKL2B | G2E3 | 0.000402503 | Genetic Interactions | Lin-Smith-2010 |
| DLGAP4 | SPIRE1 | 0.000589302 | Genetic Interactions | Lin-Smith-2010 |
| DLGAP4 | DOK6 | 0.00074885 | Genetic Interactions | Lin-Smith-2010 |
| DLGAP4 | ABLIM2 | 0.000629132 | Genetic Interactions | Lin-Smith-2010 |
| DLGAP4 | MSI2 | 0.000475551 | Genetic Interactions | Lin-Smith-2010 |
| DLGAP4 | NUBPL | 0.000727844 | Genetic Interactions | Lin-Smith-2010 |

| | | | | |
|---|---|---|---|---|
| BTBD1 | SLITRK1 | 0.000572462 | Genetic Interactions | Lin-Smith-2010 |
| BTBD1 | C20orf24 | 0.00055286 | Genetic Interactions | Lin-Smith-2010 |
| BTBD1 | NOSTRIN | 0.000510069 | Genetic Interactions | Lin-Smith-2010 |
| BTBD1 | NUBPL | 0.000628895 | Genetic Interactions | Lin-Smith-2010 |
| BTBD1 | MSRA | 0.000593782 | Genetic Interactions | Lin-Smith-2010 |
| BTBD1 | G2E3 | 0.000394262 | Genetic Interactions | Lin-Smith-2010 |
| BTBD1 | BIRC6 | 0.000952257 | Genetic Interactions | Lin-Smith-2010 |
| BTBD1 | ASB1 | 0.000451568 | Genetic Interactions | Lin-Smith-2010 |
| BTBD1 | DLGAP4 | 0.000560014 | Genetic Interactions | Lin-Smith-2010 |
| G6PC2 | CRIM1 | 0.001985808 | Genetic Interactions | Lin-Smith-2010 |
| RNF3A | C9orf72 | 0.000622218 | Genetic Interactions | Lin-Smith-2010 |
| RNF3A | ASB1 | 0.000729217 | Genetic Interactions | Lin-Smith-2010 |
| AP3D1 | MSI2 | 0.001261905 | Genetic Interactions | Lin-Smith-2010 |
| AP3D1 | C6orf15 | 0.007687661 | Genetic Interactions | Lin-Smith-2010 |
| AP3D1 | G2E3 | 0.001210806 | Genetic Interactions | Lin-Smith-2010 |
| HSPBAP1 | SLC10A7 | 0.002361066 | Genetic Interactions | Lin-Smith-2010 |
| HSPBAP1 | C9orf72 | 0.000482181 | Genetic Interactions | Lin-Smith-2010 |
| HSPBAP1 | CRIM1 | 0.000467527 | Genetic Interactions | Lin-Smith-2010 |
| HSPBAP1 | C20orf24 | 0.000691858 | Genetic Interactions | Lin-Smith-2010 |
| HSPBAP1 | MOBP | 0.001006613 | Genetic Interactions | Lin-Smith-2010 |
| HSPBAP1 | NUBPL | 0.000787009 | Genetic Interactions | Lin-Smith-2010 |
| HSPBAP1 | RGS12 | 0.000841781 | Genetic Interactions | Lin-Smith-2010 |
| HSPBAP1 | G2E3 | 0.000493386 | Genetic Interactions | Lin-Smith-2010 |
| HSPBAP1 | DLGAP4 | 0.00070081 | Genetic Interactions | Lin-Smith-2010 |
| KIF5A | ABLIM2 | 0.002124871 | Genetic Interactions | Lin-Smith-2010 |
| KIF5A | HSPBAP1 | 0.002366959 | Genetic Interactions | Lin-Smith-2010 |
| HSD17B6 | SLC10A7 | 0.003486902 | Genetic Interactions | Lin-Smith-2010 |
| HSD17B6 | C9orf72 | 0.000712101 | Genetic Interactions | Lin-Smith-2010 |
| HSD17B6 | MOBKL2B | 0.000912968 | Genetic Interactions | Lin-Smith-2010 |
| HMGB3 | APCDD1L | 0.001253188 | Genetic Interactions | Lin-Smith-2010 |
| HMGB3 | CRIM1 | 0.000571999 | Genetic Interactions | Lin-Smith-2010 |
| HMGB3 | BTBD1 | 0.000740848 | Genetic Interactions | Lin-Smith-2010 |
| NFASC | APCDD1L | 0.000818558 | Genetic Interactions | Lin-Smith-2010 |
| NFASC | SPIRE1 | 0.000509215 | Genetic Interactions | Lin-Smith-2010 |
| NFASC | C9orf72 | 0.000385329 | Genetic Interactions | Lin-Smith-2010 |
| NFASC | MSI2 | 0.000410924 | Genetic Interactions | Lin-Smith-2010 |
| NFASC | SLMO1 | 0.001011689 | Genetic Interactions | Lin-Smith-2010 |
| NFASC | DLGAP4 | 0.000560044 | Genetic Interactions | Lin-Smith-2010 |

| | | | | |
|---|---|---|---|---|
| TNPO3 | VARS2 | 0.001273294 | Genetic Interactions | Lin-Smith-2010 |
| TNPO3 | C9orf72 | 0.000674222 | Genetic Interactions | Lin-Smith-2010 |
| TNPO3 | RGS12 | 0.001177041 | Genetic Interactions | Lin-Smith-2010 |
| TNPO3 | MOBKL2B | 0.000864404 | Genetic Interactions | Lin-Smith-2010 |
| BNIP1 | COL24A1 | 0.00149254 | Genetic Interactions | Lin-Smith-2010 |
| BNIP1 | ASB1 | 0.001156766 | Genetic Interactions | Lin-Smith-2010 |
| BNIP1 | HSD17B6 | 0.002290838 | Genetic Interactions | Lin-Smith-2010 |
| ADH1A | C9orf72 | 0.00122861 | Genetic Interactions | Lin-Smith-2010 |
| CYR61 | ZFYVE28 | 0.003760334 | Genetic Interactions | Lin-Smith-2010 |
| CYR61 | TSPAN32 | 0.003028401 | Genetic Interactions | Lin-Smith-2010 |
| CYR61 | G2E3 | 0.001533382 | Genetic Interactions | Lin-Smith-2010 |
| CYR61 | RNF3A | 0.00303905 | Genetic Interactions | Lin-Smith-2010 |
| NUP50 | C20orf24 | 0.001073208 | Genetic Interactions | Lin-Smith-2010 |
| NUP50 | NDST1 | 0.002240962 | Genetic Interactions | Lin-Smith-2010 |
| NUP50 | MSRA | 0.001152647 | Genetic Interactions | Lin-Smith-2010 |
| NUP50 | TSPAN32 | 0.001511531 | Genetic Interactions | Lin-Smith-2010 |
| NUP50 | ASB1 | 0.000876581 | Genetic Interactions | Lin-Smith-2010 |
| CNTN6 | COL24A1 | 0.000548623 | Genetic Interactions | Lin-Smith-2010 |
| CNTN6 | CRIM1 | 0.000351783 | Genetic Interactions | Lin-Smith-2010 |
| CNTN6 | MSI2 | 0.000386908 | Genetic Interactions | Lin-Smith-2010 |
| CNTN6 | SLITRK1 | 0.000539036 | Genetic Interactions | Lin-Smith-2010 |
| CNTN6 | C20orf24 | 0.000520578 | Genetic Interactions | Lin-Smith-2010 |
| CNTN6 | NOSTRIN | 0.000480285 | Genetic Interactions | Lin-Smith-2010 |
| CNTN6 | MSRA | 0.000559111 | Genetic Interactions | Lin-Smith-2010 |
| CNTN6 | TSPAN32 | 0.000733194 | Genetic Interactions | Lin-Smith-2010 |
| CNTN6 | G2E3 | 0.000371241 | Genetic Interactions | Lin-Smith-2010 |
| CNTN6 | BIRC6 | 0.000896654 | Genetic Interactions | Lin-Smith-2010 |
| CNTN6 | DLGAP4 | 0.000527314 | Genetic Interactions | Lin-Smith-2010 |
| CNTN6 | CYR61 | 0.001772044 | Genetic Interactions | Lin-Smith-2010 |
| GDNF | APCDD1L | 0.001245612 | Genetic Interactions | Lin-Smith-2010 |
| GDNF | MSI2 | 0.000625309 | Genetic Interactions | Lin-Smith-2010 |
| GDNF | G2E3 | 0.000599987 | Genetic Interactions | Lin-Smith-2010 |
| GDNF | MOBKL2B | 0.00075176 | Genetic Interactions | Lin-Smith-2010 |
| GDNF | DLGAP4 | 0.000852228 | Genetic Interactions | Lin-Smith-2010 |
| GDNF | RNF3A | 0.001189131 | Genetic Interactions | Lin-Smith-2010 |
| GNG7 | C15orf40 | 0.001244525 | Genetic Interactions | Lin-Smith-2010 |
| GNG7 | COL24A1 | 0.000807229 | Genetic Interactions | Lin-Smith-2010 |
| GNG7 | BTBD1 | 0.000670397 | Genetic Interactions | Lin-Smith-2010 |

| | | | | |
|---|---|---|---|---|
| GNG7 | HSPBAP1 | 0.000838945 | Genetic Interactions | Lin-Smith-2010 |
| GNG7 | TNPO3 | 0.001173076 | Genetic Interactions | Lin-Smith-2010 |
| GNG7 | CNTN6 | 0.000631252 | Genetic Interactions | Lin-Smith-2010 |
| GPX3 | C15orf40 | 0.002488639 | Genetic Interactions | Lin-Smith-2010 |
| GPX3 | COL24A1 | 0.001614192 | Genetic Interactions | Lin-Smith-2010 |
| GPX3 | SLITRK1 | 0.001585983 | Genetic Interactions | Lin-Smith-2010 |
| GPX3 | CPNE7 | 0.008813995 | Genetic Interactions | Lin-Smith-2010 |
| GPX3 | BTBD1 | 0.001340572 | Genetic Interactions | Lin-Smith-2010 |
| TG | C15orf40 | 0.000844686 | Genetic Interactions | Lin-Smith-2010 |
| TG | DOK6 | 0.000608443 | Genetic Interactions | Lin-Smith-2010 |
| TG | NOSTRIN | 0.000479638 | Genetic Interactions | Lin-Smith-2010 |
| TG | G2E3 | 0.00037074 | Genetic Interactions | Lin-Smith-2010 |
| TG | DLGAP4 | 0.000526603 | Genetic Interactions | Lin-Smith-2010 |
| TG | BTBD1 | 0.000455012 | Genetic Interactions | Lin-Smith-2010 |
| TG | HMGB3 | 0.000696649 | Genetic Interactions | Lin-Smith-2010 |
| TBK1 | SLMO1 | 0.003788822 | Genetic Interactions | Lin-Smith-2010 |
| TBK1 | SCN5A | 0.00321341 | Genetic Interactions | Lin-Smith-2010 |
| PAX8 | DOK6 | 0.001271392 | Genetic Interactions | Lin-Smith-2010 |
| PAX8 | CRIM1 | 0.00073409 | Genetic Interactions | Lin-Smith-2010 |
| PAX8 | C20orf24 | 0.001086324 | Genetic Interactions | Lin-Smith-2010 |
| PAX8 | DLGAP4 | 0.001100381 | Genetic Interactions | Lin-Smith-2010 |
| UGCG | COL24A1 | 0.000797234 | Genetic Interactions | Lin-Smith-2010 |
| UGCG | NOSTRIN | 0.000697928 | Genetic Interactions | Lin-Smith-2010 |
| UGCG | ASB1 | 0.000617881 | Genetic Interactions | Lin-Smith-2010 |
| UGCG | TG | 0.000622594 | Genetic Interactions | Lin-Smith-2010 |
| SETD2 | VARS2 | 0.001192218 | Genetic Interactions | Lin-Smith-2010 |
| SETD2 | TSPAN32 | 0.001275762 | Genetic Interactions | Lin-Smith-2010 |
| SETD2 | BIRC6 | 0.001560183 | Genetic Interactions | Lin-Smith-2010 |
| PCDH9 | SPIRE1 | 0.000439212 | Genetic Interactions | Lin-Smith-2010 |
| PCDH9 | ZFYVE28 | 0.000833983 | Genetic Interactions | Lin-Smith-2010 |
| PCDH9 | GGNBP2 | 0.008580335 | Genetic Interactions | Lin-Smith-2010 |
| PCDH9 | CRIM1 | 0.000322256 | Genetic Interactions | Lin-Smith-2010 |
| PCDH9 | MSI2 | 0.000354433 | Genetic Interactions | Lin-Smith-2010 |
| PCDH9 | SLMO1 | 0.000872608 | Genetic Interactions | Lin-Smith-2010 |
| PCDH9 | TSPAN32 | 0.000671652 | Genetic Interactions | Lin-Smith-2010 |
| PCDH9 | G2E3 | 0.00034008 | Genetic Interactions | Lin-Smith-2010 |
| PCDH9 | HSPBAP1 | 0.00052232 | Genetic Interactions | Lin-Smith-2010 |
| PCDH9 | HMGB3 | 0.000639036 | Genetic Interactions | Lin-Smith-2010 |

| | | | | |
|---|---|---|---|---|
| PCDH9 | GPX3 | 0.001156342 | Genetic Interactions | Lin-Smith-2010 |
| PCDH9 | TG | 0.000392482 | Genetic Interactions | Lin-Smith-2010 |
| ACADS | VARS2 | 0.001351041 | Genetic Interactions | Lin-Smith-2010 |
| ACADS | CRIM1 | 0.000693648 | Genetic Interactions | Lin-Smith-2010 |
| ACADS | C20orf24 | 0.001026477 | Genetic Interactions | Lin-Smith-2010 |
| ACADS | NOSTRIN | 0.000947028 | Genetic Interactions | Lin-Smith-2010 |
| ACADS | ELP3 | 0.001555159 | Genetic Interactions | Lin-Smith-2010 |
| ACADS | SETD2 | 0.001471954 | Genetic Interactions | Lin-Smith-2010 |
| SAR1B | APCDD1L | 0.001038504 | Genetic Interactions | Lin-Smith-2010 |
| SAR1B | C20orf24 | 0.000701451 | Genetic Interactions | Lin-Smith-2010 |
| SAR1B | SLMO1 | 0.001283528 | Genetic Interactions | Lin-Smith-2010 |
| SAR1B | DLGAP4 | 0.000710527 | Genetic Interactions | Lin-Smith-2010 |
| SAR1B | RNF3A | 0.000991413 | Genetic Interactions | Lin-Smith-2010 |
| SAR1B | HMGB3 | 0.000939964 | Genetic Interactions | Lin-Smith-2010 |
| SAR1B | NFASC | 0.000613966 | Genetic Interactions | Lin-Smith-2010 |
| SAR1B | CNTN6 | 0.000578085 | Genetic Interactions | Lin-Smith-2010 |
| NTN1 | NDST1 | 0.002279369 | Genetic Interactions | Lin-Smith-2010 |
| NTN1 | RGS12 | 0.001328147 | Genetic Interactions | Lin-Smith-2010 |
| NTN1 | G2E3 | 0.000778456 | Genetic Interactions | Lin-Smith-2010 |
| NTN1 | TG | 0.000898405 | Genetic Interactions | Lin-Smith-2010 |
| NTN1 | SAR1B | 0.001212187 | Genetic Interactions | Lin-Smith-2010 |
| parkin | SPIRE1 | 0.000304028 | Genetic Interactions | Lin-Smith-2010 |
| parkin | COL24A1 | 0.000347888 | Genetic Interactions | Lin-Smith-2010 |
| parkin | C9orf72 | 0.000230062 | Genetic Interactions | Lin-Smith-2010 |
| parkin | DOK6 | 0.000386342 | Genetic Interactions | Lin-Smith-2010 |
| parkin | ABLIM2 | 0.000324577 | Genetic Interactions | Lin-Smith-2010 |
| parkin | MSI2 | 0.000245343 | Genetic Interactions | Lin-Smith-2010 |
| parkin | RGS12 | 0.000401637 | Genetic Interactions | Lin-Smith-2010 |
| parkin | ASB1 | 0.000269625 | Genetic Interactions | Lin-Smith-2010 |
| parkin | MOBKL2B | 0.000294957 | Genetic Interactions | Lin-Smith-2010 |
| parkin | BTBD1 | 0.000288918 | Genetic Interactions | Lin-Smith-2010 |
| parkin | KIF5A | 0.001129342 | Genetic Interactions | Lin-Smith-2010 |
| parkin | HSD17B6 | 0.000533959 | Genetic Interactions | Lin-Smith-2010 |
| parkin | BNIP1 | 0.000740112 | Genetic Interactions | Lin-Smith-2010 |
| parkin | PCDH9 | 0.000249213 | Genetic Interactions | Lin-Smith-2010 |
| MCC | COL24A1 | 0.000800157 | Genetic Interactions | Lin-Smith-2010 |
| MCC | TSPAN32 | 0.00106935 | Genetic Interactions | Lin-Smith-2010 |
| MCC | BTBD1 | 0.000664524 | Genetic Interactions | Lin-Smith-2010 |

| | | | | |
|---|---|---|---|---|
| MCC | GDNF | 0.001011271 | Genetic Interactions | Lin-Smith-2010 |
| ELMO1 | C15orf40 | 0.001475561 | Genetic Interactions | Lin-Smith-2010 |
| ELMO1 | DOK6 | 0.001062874 | Genetic Interactions | Lin-Smith-2010 |
| ELMO1 | CRIM1 | 0.000613693 | Genetic Interactions | Lin-Smith-2010 |
| ELMO1 | NUP50 | 0.001542958 | Genetic Interactions | Lin-Smith-2010 |
| CAT | CRIM1 | 0.000682176 | Genetic Interactions | Lin-Smith-2010 |
| CAT | DCTN4 | 0.004419341 | Genetic Interactions | Lin-Smith-2010 |
| CAT | HMGB3 | 0.00135276 | Genetic Interactions | Lin-Smith-2010 |
| CAT | TNPO3 | 0.001546055 | Genetic Interactions | Lin-Smith-2010 |
| CAT | CNTN6 | 0.000831958 | Genetic Interactions | Lin-Smith-2010 |
| CAT | TBK1 | 0.003309112 | Genetic Interactions | Lin-Smith-2010 |
| CAT | PCDH9 | 0.000762126 | Genetic Interactions | Lin-Smith-2010 |
| KIAA0917 | APCDD1L | 0.000689898 | Genetic Interactions | Lin-Smith-2010 |
| KIAA0917 | C15orf40 | 0.000757129 | Genetic Interactions | Lin-Smith-2010 |
| KIAA0917 | DOK6 | 0.000545374 | Genetic Interactions | Lin-Smith-2010 |
| KIAA0917 | MSI2 | 0.000346335 | Genetic Interactions | Lin-Smith-2010 |
| KIAA0917 | SLITRK1 | 0.00048251 | Genetic Interactions | Lin-Smith-2010 |
| KIAA0917 | NDST1 | 0.000973027 | Genetic Interactions | Lin-Smith-2010 |
| KIAA0917 | TSPAN32 | 0.000656308 | Genetic Interactions | Lin-Smith-2010 |
| KIAA0917 | BTBD1 | 0.000407848 | Genetic Interactions | Lin-Smith-2010 |
| KIAA0917 | AP3D1 | 0.001252528 | Genetic Interactions | Lin-Smith-2010 |
| KIAA0917 | HSPBAP1 | 0.000510387 | Genetic Interactions | Lin-Smith-2010 |
| KIAA0917 | BNIP1 | 0.001044769 | Genetic Interactions | Lin-Smith-2010 |
| KIAA0917 | ADH1A | 0.00130048 | Genetic Interactions | Lin-Smith-2010 |
| KIAA0917 | GDNF | 0.000620662 | Genetic Interactions | Lin-Smith-2010 |
| KIAA0917 | SETD2 | 0.00066822 | Genetic Interactions | Lin-Smith-2010 |
| KIAA0917 | MCC | 0.000560105 | Genetic Interactions | Lin-Smith-2010 |
| MAPK9 | MSI2 | 0.000454387 | Genetic Interactions | Lin-Smith-2010 |
| MAPK9 | TSPAN32 | 0.000861067 | Genetic Interactions | Lin-Smith-2010 |
| MAPK9 | HTR2A | 0.001341849 | Genetic Interactions | Lin-Smith-2010 |
| MAPK9 | CNTN6 | 0.000503846 | Genetic Interactions | Lin-Smith-2010 |
| MAPK9 | ACADS | 0.000993486 | Genetic Interactions | Lin-Smith-2010 |
| MAPK9 | parkin | 0.000319495 | Genetic Interactions | Lin-Smith-2010 |
| MAPK9 | ELMO1 | 0.00087897 | Genetic Interactions | Lin-Smith-2010 |
| TEK | SLC10A7 | 0.002701482 | Genetic Interactions | Lin-Smith-2010 |
| TEK | C6orf15 | 0.003584268 | Genetic Interactions | Lin-Smith-2010 |
| TEK | MOBP | 0.001151745 | Genetic Interactions | Lin-Smith-2010 |
| TEK | NUBPL | 0.000900479 | Genetic Interactions | Lin-Smith-2010 |

| | | | | |
|---|---|---|---|---|
| TEK | KIF5A | 0.002708225 | Genetic Interactions | Lin-Smith-2010 |
| TEK | HMGB3 | 0.001060778 | Genetic Interactions | Lin-Smith-2010 |
| TEK | PCDH9 | 0.000597627 | Genetic Interactions | Lin-Smith-2010 |
| TEK | parkin | 0.000413686 | Genetic Interactions | Lin-Smith-2010 |
| SMARCA2 | SPIRE1 | 0.000321281 | Genetic Interactions | Lin-Smith-2010 |
| SMARCA2 | ABLIM2 | 0.000342997 | Genetic Interactions | Lin-Smith-2010 |
| SMARCA2 | CRIM1 | 0.000235729 | Genetic Interactions | Lin-Smith-2010 |
| SMARCA2 | MSI2 | 0.000259266 | Genetic Interactions | Lin-Smith-2010 |
| SMARCA2 | C19orf57 | 0.001609237 | Genetic Interactions | Lin-Smith-2010 |
| SMARCA2 | NUBPL | 0.000396813 | Genetic Interactions | Lin-Smith-2010 |
| SMARCA2 | TSPAN32 | 0.00049131 | Genetic Interactions | Lin-Smith-2010 |
| SMARCA2 | G2E3 | 0.000248767 | Genetic Interactions | Lin-Smith-2010 |
| SMARCA2 | BIRC6 | 0.000600844 | Genetic Interactions | Lin-Smith-2010 |
| SMARCA2 | HSPBAP1 | 0.000382074 | Genetic Interactions | Lin-Smith-2010 |
| SMARCA2 | HSD17B6 | 0.00056426 | Genetic Interactions | Lin-Smith-2010 |
| SMARCA2 | NFASC | 0.00030533 | Genetic Interactions | Lin-Smith-2010 |
| SMARCA2 | BNIP1 | 0.000782112 | Genetic Interactions | Lin-Smith-2010 |
| SMARCA2 | CYR61 | 0.00118744 | Genetic Interactions | Lin-Smith-2010 |
| SMARCA2 | GDNF | 0.000464626 | Genetic Interactions | Lin-Smith-2010 |
| SMARCA2 | GNG7 | 0.000422999 | Genetic Interactions | Lin-Smith-2010 |
| SMARCA2 | SETD2 | 0.000500228 | Genetic Interactions | Lin-Smith-2010 |
| SMARCA2 | PCDH9 | 0.000263356 | Genetic Interactions | Lin-Smith-2010 |
| SMARCA2 | parkin | 0.000182298 | Genetic Interactions | Lin-Smith-2010 |
| SMARCA2 | MCC | 0.000419294 | Genetic Interactions | Lin-Smith-2010 |
| SMARCA2 | CAT | 0.000557491 | Genetic Interactions | Lin-Smith-2010 |
| SMARCA2 | KIAA0917 | 0.000257339 | Genetic Interactions | Lin-Smith-2010 |
| SMARCA2 | MAPK9 | 0.000337625 | Genetic Interactions | Lin-Smith-2010 |
| GLI3 | SPIRE1 | 0.000455204 | Genetic Interactions | Lin-Smith-2010 |
| GLI3 | ABLIM2 | 0.000485971 | Genetic Interactions | Lin-Smith-2010 |
| GLI3 | NOSTRIN | 0.000455992 | Genetic Interactions | Lin-Smith-2010 |
| GLI3 | MOBP | 0.0007191 | Genetic Interactions | Lin-Smith-2010 |
| GLI3 | ELP3 | 0.000748806 | Genetic Interactions | Lin-Smith-2010 |
| GLI3 | RGS12 | 0.000601348 | Genetic Interactions | Lin-Smith-2010 |
| GLI3 | MOBKL2B | 0.000441622 | Genetic Interactions | Lin-Smith-2010 |
| GLI3 | BTBD1 | 0.000432581 | Genetic Interactions | Lin-Smith-2010 |
| GLI3 | HSPBAP1 | 0.000541339 | Genetic Interactions | Lin-Smith-2010 |
| GLI3 | KIF5A | 0.0016909 | Genetic Interactions | Lin-Smith-2010 |
| GLI3 | NUP50 | 0.000839723 | Genetic Interactions | Lin-Smith-2010 |

| | | | | |
|---|---|---|---|---|
| GLI3 | parkin | 0.000258288 | Genetic Interactions | Lin-Smith-2010 |
| GLI3 | GIPC1 | 0.003251437 | Genetic Interactions | Lin-Smith-2010 |
| GLI3 | MAPK9 | 0.000478361 | Genetic Interactions | Lin-Smith-2010 |
| ATXN2 | UGCG | 0.00235099 | Genetic Interactions | Lin-Smith-2010 |
| ATXN2 | MCC | 0.002359613 | Genetic Interactions | Lin-Smith-2010 |
| NSF | RNF3A | 0.001577133 | Genetic Interactions | Lin-Smith-2010 |
| NSF | CYR61 | 0.003798389 | Genetic Interactions | Lin-Smith-2010 |
| NSF | GDNF | 0.001486248 | Genetic Interactions | Lin-Smith-2010 |
| NSF | SETD2 | 0.001600132 | Genetic Interactions | Lin-Smith-2010 |
| NSF | parkin | 0.000583137 | Genetic Interactions | Lin-Smith-2010 |
| A2BP1 | APCDD1L | 0.000398546 | Genetic Interactions | Lin-Smith-2010 |
| A2BP1 | SPIRE1 | 0.000247931 | Genetic Interactions | Lin-Smith-2010 |
| A2BP1 | COL24A1 | 0.000283698 | Genetic Interactions | Lin-Smith-2010 |
| A2BP1 | DOK6 | 0.000315056 | Genetic Interactions | Lin-Smith-2010 |
| A2BP1 | ABLIM2 | 0.000264688 | Genetic Interactions | Lin-Smith-2010 |
| A2BP1 | CRIM1 | 0.00018191 | Genetic Interactions | Lin-Smith-2010 |
| A2BP1 | SLITRK1 | 0.00027874 | Genetic Interactions | Lin-Smith-2010 |
| A2BP1 | NDST1 | 0.000562106 | Genetic Interactions | Lin-Smith-2010 |
| A2BP1 | MOBP | 0.000391664 | Genetic Interactions | Lin-Smith-2010 |
| A2BP1 | NUBPL | 0.000306218 | Genetic Interactions | Lin-Smith-2010 |
| A2BP1 | IL37 | 0.000702629 | Genetic Interactions | Lin-Smith-2010 |
| A2BP1 | DHRS11 | 0.001075725 | Genetic Interactions | Lin-Smith-2010 |
| A2BP1 | G2E3 | 0.000191972 | Genetic Interactions | Lin-Smith-2010 |
| A2BP1 | MOBKL2B | 0.000240533 | Genetic Interactions | Lin-Smith-2010 |
| A2BP1 | BTBD1 | 0.000235609 | Genetic Interactions | Lin-Smith-2010 |
| A2BP1 | HSPBAP1 | 0.000294845 | Genetic Interactions | Lin-Smith-2010 |
| A2BP1 | KIF5A | 0.000920963 | Genetic Interactions | Lin-Smith-2010 |
| A2BP1 | NFASC | 0.000235622 | Genetic Interactions | Lin-Smith-2010 |
| A2BP1 | CNTN6 | 0.000221851 | Genetic Interactions | Lin-Smith-2010 |
| A2BP1 | COL16A1 | 0.001099903 | Genetic Interactions | Lin-Smith-2010 |
| A2BP1 | TG | 0.000221552 | Genetic Interactions | Lin-Smith-2010 |
| A2BP1 | PCDH9 | 0.00020323 | Genetic Interactions | Lin-Smith-2010 |
| A2BP1 | ACADS | 0.000437447 | Genetic Interactions | Lin-Smith-2010 |
| A2BP1 | parkin | 0.000140679 | Genetic Interactions | Lin-Smith-2010 |
| A2BP1 | MCC | 0.000323566 | Genetic Interactions | Lin-Smith-2010 |
| A2BP1 | ELMO1 | 0.000387024 | Genetic Interactions | Lin-Smith-2010 |
| A2BP1 | KIAA0917 | 0.000198587 | Genetic Interactions | Lin-Smith-2010 |
| A2BP1 | SMARCA2 | 0.000148662 | Genetic Interactions | Lin-Smith-2010 |

| | | | | |
|---|---|---|---|---|
| A2BP1 | GLI3 | 0.00021063 | Genetic Interactions | Lin-Smith-2010 |
| STX8 | SLMO1 | 0.001263258 | Genetic Interactions | Lin-Smith-2010 |
| STX8 | G2E3 | 0.000492327 | Genetic Interactions | Lin-Smith-2010 |
| STX8 | BTBD1 | 0.000604237 | Genetic Interactions | Lin-Smith-2010 |
| STX8 | CNTN6 | 0.000568955 | Genetic Interactions | Lin-Smith-2010 |
| STX8 | SAR1B | 0.000766637 | Genetic Interactions | Lin-Smith-2010 |
| STX8 | GLI3 | 0.000540177 | Genetic Interactions | Lin-Smith-2010 |
| BTBD8 | MOBKL2B | 0.002969787 | Genetic Interactions | Lin-Smith-2010 |
| DNM2 | VARS2 | 0.00219467 | Genetic Interactions | Lin-Smith-2010 |
| DNM2 | NOSTRIN | 0.001538379 | Genetic Interactions | Lin-Smith-2010 |
| DNM2 | CNTN6 | 0.001374182 | Genetic Interactions | Lin-Smith-2010 |
| DNM2 | TG | 0.001372329 | Genetic Interactions | Lin-Smith-2010 |
| DNM2 | ATXN2 | 0.005182076 | Genetic Interactions | Lin-Smith-2010 |
| DNM2 | A2BP1 | 0.000710602 | Genetic Interactions | Lin-Smith-2010 |
| NUDC | ABLIM2 | 0.001067475 | Genetic Interactions | Lin-Smith-2010 |
| NUDC | TG | 0.000893509 | Genetic Interactions | Lin-Smith-2010 |
| NUDC | SMARCA2 | 0.000599545 | Genetic Interactions | Lin-Smith-2010 |
| NUDC | A2BP1 | 0.000462665 | Genetic Interactions | Lin-Smith-2010 |
| GOSR1 | NUBPL | 0.000798308 | Genetic Interactions | Lin-Smith-2010 |
| GOSR1 | TSPAN32 | 0.000988418 | Genetic Interactions | Lin-Smith-2010 |
| GOSR1 | G2E3 | 0.000500469 | Genetic Interactions | Lin-Smith-2010 |
| GOSR1 | BTBD1 | 0.00061423 | Genetic Interactions | Lin-Smith-2010 |
| GOSR1 | ADH1A | 0.001958559 | Genetic Interactions | Lin-Smith-2010 |
| GOSR1 | GNG7 | 0.000850989 | Genetic Interactions | Lin-Smith-2010 |
| GOSR1 | SETD2 | 0.001006358 | Genetic Interactions | Lin-Smith-2010 |
| GOSR1 | parkin | 0.000366748 | Genetic Interactions | Lin-Smith-2010 |
| GOSR1 | MAPK9 | 0.000679234 | Genetic Interactions | Lin-Smith-2010 |
| GOSR1 | GLI3 | 0.00054911 | Genetic Interactions | Lin-Smith-2010 |
| GOSR1 | A2BP1 | 0.000299078 | Genetic Interactions | Lin-Smith-2010 |
| CNTNAP1 | DLGAP4 | 0.001216501 | Genetic Interactions | Lin-Smith-2010 |
| CNTNAP1 | BTBD1 | 0.00105112 | Genetic Interactions | Lin-Smith-2010 |
| CNTNAP1 | HSD17B6 | 0.00194261 | Genetic Interactions | Lin-Smith-2010 |
| CNTNAP1 | parkin | 0.000627608 | Genetic Interactions | Lin-Smith-2010 |
| CNTNAP1 | MCC | 0.001443524 | Genetic Interactions | Lin-Smith-2010 |
| CNTNAP1 | A2BP1 | 0.000511806 | Genetic Interactions | Lin-Smith-2010 |
| DOCK2 | MOBP | 0.000808019 | Genetic Interactions | Lin-Smith-2010 |
| DOCK2 | MSRA | 0.000596469 | Genetic Interactions | Lin-Smith-2010 |
| DOCK2 | SCN5A | 0.000861878 | Genetic Interactions | Lin-Smith-2010 |

| | | | | |
|---|---|---|---|---|
| DOCK2 | BIRC6 | 0.000956566 | Genetic Interactions | Lin-Smith-2010 |
| DOCK2 | ASB1 | 0.000453611 | Genetic Interactions | Lin-Smith-2010 |
| DOCK2 | MOBKL2B | 0.00049623 | Genetic Interactions | Lin-Smith-2010 |
| DOCK2 | HMGB3 | 0.0007442 | Genetic Interactions | Lin-Smith-2010 |
| DOCK2 | NFASC | 0.000486097 | Genetic Interactions | Lin-Smith-2010 |
| DOCK2 | parkin | 0.000290225 | Genetic Interactions | Lin-Smith-2010 |
| DOCK2 | CAT | 0.000887546 | Genetic Interactions | Lin-Smith-2010 |
| DOCK2 | KIAA0917 | 0.000409693 | Genetic Interactions | Lin-Smith-2010 |
| DOCK2 | A2BP1 | 0.000236675 | Genetic Interactions | Lin-Smith-2010 |
| STX12 | C9orf72 | 0.000327897 | Genetic Interactions | Lin-Smith-2010 |
| STX12 | CRIM1 | 0.000317931 | Genetic Interactions | Lin-Smith-2010 |
| STX12 | SLITRK1 | 0.000487164 | Genetic Interactions | Lin-Smith-2010 |
| STX12 | C20orf24 | 0.000470483 | Genetic Interactions | Lin-Smith-2010 |
| STX12 | NOSTRIN | 0.000434067 | Genetic Interactions | Lin-Smith-2010 |
| STX12 | NUBPL | 0.000535188 | Genetic Interactions | Lin-Smith-2010 |
| STX12 | MSRA | 0.000505308 | Genetic Interactions | Lin-Smith-2010 |
| STX12 | HMGB3 | 0.00063046 | Genetic Interactions | Lin-Smith-2010 |
| STX12 | BNIP1 | 0.001054847 | Genetic Interactions | Lin-Smith-2010 |
| STX12 | ADH1A | 0.001313024 | Genetic Interactions | Lin-Smith-2010 |
| STX12 | NUP50 | 0.000799348 | Genetic Interactions | Lin-Smith-2010 |
| STX12 | TG | 0.000387215 | Genetic Interactions | Lin-Smith-2010 |
| STX12 | PAX8 | 0.000809117 | Genetic Interactions | Lin-Smith-2010 |
| STX12 | SETD2 | 0.000674666 | Genetic Interactions | Lin-Smith-2010 |
| STX12 | PCDH9 | 0.000355192 | Genetic Interactions | Lin-Smith-2010 |
| STX12 | MCC | 0.000565508 | Genetic Interactions | Lin-Smith-2010 |
| STX12 | TEK | 0.000589607 | Genetic Interactions | Lin-Smith-2010 |
| STX12 | A2BP1 | 0.000200503 | Genetic Interactions | Lin-Smith-2010 |
| STX12 | GOSR1 | 0.000522708 | Genetic Interactions | Lin-Smith-2010 |
| STX12 | DOCK2 | 0.000413645 | Genetic Interactions | Lin-Smith-2010 |
| PFKP | SPIRE1 | 0.000987192 | Genetic Interactions | Lin-Smith-2010 |
| PFKP | MOBP | 0.001559497 | Genetic Interactions | Lin-Smith-2010 |
| PFKP | RGS12 | 0.001304131 | Genetic Interactions | Lin-Smith-2010 |
| PFKP | HMGB3 | 0.001436325 | Genetic Interactions | Lin-Smith-2010 |
| PFKP | TNPO3 | 0.00164156 | Genetic Interactions | Lin-Smith-2010 |
| PFKP | SH3BP2 | 0.002485168 | Genetic Interactions | Lin-Smith-2010 |
| PFKP | ATXN2 | 0.003331139 | Genetic Interactions | Lin-Smith-2010 |
| PFKP | A2BP1 | 0.000456789 | Genetic Interactions | Lin-Smith-2010 |
| TDRD7 | NOSTRIN | 0.001830375 | Genetic Interactions | Lin-Smith-2010 |

| | | | | |
|---|---|---|---|---|
| TDRD7 | MOBP | 0.002886506 | Genetic Interactions | Lin-Smith-2010 |
| TDRD7 | MSRA | 0.002130782 | Genetic Interactions | Lin-Smith-2010 |
| TDRD7 | SCN5A | 0.003078909 | Genetic Interactions | Lin-Smith-2010 |
| STX7 | C9orf72 | 0.000477899 | Genetic Interactions | Lin-Smith-2010 |
| STX7 | ZFYVE28 | 0.001199193 | Genetic Interactions | Lin-Smith-2010 |
| STX7 | G2E3 | 0.000489005 | Genetic Interactions | Lin-Smith-2010 |
| STX7 | KIF5A | 0.002345941 | Genetic Interactions | Lin-Smith-2010 |
| STX7 | HSD17B6 | 0.001109175 | Genetic Interactions | Lin-Smith-2010 |
| STX7 | CNTN6 | 0.000565116 | Genetic Interactions | Lin-Smith-2010 |
| STX7 | TG | 0.000564354 | Genetic Interactions | Lin-Smith-2010 |
| STX7 | SAR1B | 0.000761463 | Genetic Interactions | Lin-Smith-2010 |
| STX7 | KIAA0917 | 0.000505855 | Genetic Interactions | Lin-Smith-2010 |
| STX7 | PFKP | 0.001163564 | Genetic Interactions | Lin-Smith-2010 |
| STX5 | APCDD1L | 0.001515319 | Genetic Interactions | Lin-Smith-2010 |
| STX5 | MOBKL2B | 0.000914535 | Genetic Interactions | Lin-Smith-2010 |
| STX5 | TG | 0.000842367 | Genetic Interactions | Lin-Smith-2010 |
| STX5 | A2BP1 | 0.000436184 | Genetic Interactions | Lin-Smith-2010 |
| STX5 | GOSR1 | 0.001137127 | Genetic Interactions | Lin-Smith-2010 |
| RBM24 | C15orf40 | 0.001164211 | Genetic Interactions | Lin-Smith-2010 |
| RBM24 | NOSTRIN | 0.000661074 | Genetic Interactions | Lin-Smith-2010 |
| RBM24 | RGS12 | 0.000871804 | Genetic Interactions | Lin-Smith-2010 |
| RBM24 | BTBD1 | 0.000627133 | Genetic Interactions | Lin-Smith-2010 |
| RBM24 | SH3BP2 | 0.001661319 | Genetic Interactions | Lin-Smith-2010 |
| RBM24 | MCC | 0.000861255 | Genetic Interactions | Lin-Smith-2010 |
| RBM24 | TEK | 0.000897957 | Genetic Interactions | Lin-Smith-2010 |
| RBM24 | A2BP1 | 0.00030536 | Genetic Interactions | Lin-Smith-2010 |
| RBM24 | GOSR1 | 0.000796072 | Genetic Interactions | Lin-Smith-2010 |
| RBM24 | STX12 | 0.000533689 | Genetic Interactions | Lin-Smith-2010 |
| NAPG | MSI2 | 0.000402643 | Genetic Interactions | Lin-Smith-2010 |
| NAPG | NUBPL | 0.000616256 | Genetic Interactions | Lin-Smith-2010 |
| NAPG | MSRA | 0.00058185 | Genetic Interactions | Lin-Smith-2010 |
| NAPG | G2E3 | 0.000386339 | Genetic Interactions | Lin-Smith-2010 |
| NAPG | MOBKL2B | 0.000484067 | Genetic Interactions | Lin-Smith-2010 |
| NAPG | TNPO3 | 0.000829691 | Genetic Interactions | Lin-Smith-2010 |
| NAPG | ADH1A | 0.001511916 | Genetic Interactions | Lin-Smith-2010 |
| NAPG | UGCG | 0.00064879 | Genetic Interactions | Lin-Smith-2010 |
| NAPG | TEK | 0.000678918 | Genetic Interactions | Lin-Smith-2010 |
| NAPG | GOSR1 | 0.000601886 | Genetic Interactions | Lin-Smith-2010 |

| | | | | |
|---|---|---|---|---|
| NAPG | TDRD7 | 0.001701506 | Genetic Interactions | Lin-Smith-2010 |
| IDH3B | NAPG | 0.002836541 | Genetic Interactions | Lin-Smith-2010 |
| ARSB | COL24A1 | 0.000660154 | Genetic Interactions | Lin-Smith-2010 |
| ARSB | C20orf24 | 0.000626407 | Genetic Interactions | Lin-Smith-2010 |
| ARSB | NOSTRIN | 0.000577924 | Genetic Interactions | Lin-Smith-2010 |
| ARSB | SLMO1 | 0.001146213 | Genetic Interactions | Lin-Smith-2010 |
| ARSB | BTBD1 | 0.000548252 | Genetic Interactions | Lin-Smith-2010 |
| ARSB | G6PC2 | 0.002914157 | Genetic Interactions | Lin-Smith-2010 |
| ARSB | HSPBAP1 | 0.000686092 | Genetic Interactions | Lin-Smith-2010 |
| ARSB | NFASC | 0.000548282 | Genetic Interactions | Lin-Smith-2010 |
| ARSB | CNTN6 | 0.000516239 | Genetic Interactions | Lin-Smith-2010 |
| ARSB | SETD2 | 0.00089826 | Genetic Interactions | Lin-Smith-2010 |
| ARSB | A2BP1 | 0.000266952 | Genetic Interactions | Lin-Smith-2010 |
| ARSB | STX8 | 0.00068462 | Genetic Interactions | Lin-Smith-2010 |
| ARSB | STX12 | 0.000466562 | Genetic Interactions | Lin-Smith-2010 |
| ARSB | RBM24 | 0.000710562 | Genetic Interactions | Lin-Smith-2010 |
| GDNF | DOK6 | 0.13123737 | Pathway | Wu-Stein-2010 |
| GNG7 | HTR2A | 0.009824721 | Pathway | Wu-Stein-2010 |
| PAX8 | TG | 0.37898692 | Pathway | Wu-Stein-2010 |
| CNTNAP1 | NFASC | 0.690983 | Pathway | Wu-Stein-2010 |
| VTI1B | STX8 | 0.111152895 | Pathway | Wu-Stein-2010 |
| DOCK2 | ELMO1 | 0.5975598 | Pathway | Wu-Stein-2010 |
| STX18 | BNIP1 | 0.36939806 | Pathway | Wu-Stein-2010 |
| STX12 | STX8 | 0.16513501 | Pathway | Wu-Stein-2010 |
| STX12 | VTI1B | 0.119380414 | Pathway | Wu-Stein-2010 |
| STX7 | STX8 | 0.15194081 | Pathway | Wu-Stein-2010 |
| STX7 | VTI1B | 0.10984199 | Pathway | Wu-Stein-2010 |
| STX5 | GOSR1 | 0.37009686 | Pathway | Wu-Stein-2010 |
| GDNF | DOK6 | 0.17095798 | Pathway | NCI_NATURE |
| ELMO1 | NTN1 | 0.06397213 | Pathway | NCI_NATURE |
| DOCK2 | ELMO1 | 0.31301603 | Pathway | NCI_NATURE |
| NSF | KIAA0917 | 0.14760774 | Physical Interactions | Behrends-Harper-2010 |
| STX8 | NSF | 0.28700674 | Physical Interactions | Behrends-Harper-2010 |
| BTBD8 | WIPI2 | 0.42877966 | Physical Interactions | Behrends-Harper-2010 |
| DNM2 | WIPI2 | 0.42877966 | Physical Interactions | Behrends-Harper-2010 |
| NUDC | WIPI2 | 0.42877966 | Physical Interactions | Behrends-Harper-2010 |
| GOSR1 | KIAA0917 | 0.3685327 | Physical Interactions | Behrends-Harper-2010 |
| GOSR1 | NSF | 0.05496511 | Physical Interactions | Behrends-Harper-2010 |

| | | | | |
|---|---|---|---|---|
| VTI1B | NSF | 0.28700674 | Physical Interactions | Behrends-Harper-2010 |
| STX12 | NSF | 0.28700674 | Physical Interactions | Behrends-Harper-2010 |
| PFKP | WIPI2 | 0.29525253 | Physical Interactions | Behrends-Harper-2010 |
| STX7 | NSF | 0.28700674 | Physical Interactions | Behrends-Harper-2010 |
| STX5 | NSF | 0.28700674 | Physical Interactions | Behrends-Harper-2010 |
| SNAP29 | NSF | 0.28700674 | Physical Interactions | Behrends-Harper-2010 |
| NAPG | NSF | 0.28700674 | Physical Interactions | Behrends-Harper-2010 |
| A2BP1 | TNIP1 | 0.26131737 | Physical Interactions | Lim-Zoghbi-2006 |
| A2BP1 | ATXN2 | 0.025902292 | Physical Interactions | Lim-Zoghbi-2006 |
| CIC | SETD2 | 0.2900025 | Physical Interactions | Lim-Zoghbi-2006 |
| TDRD7 | ATXN2 | 0.39532113 | Physical Interactions | Lim-Zoghbi-2006 |
| RBM24 | A2BP1 | 0.26131737 | Physical Interactions | Lim-Zoghbi-2006 |
| IDH3B | ATXN2 | 0.39532113 | Physical Interactions | Lim-Zoghbi-2006 |
| STX8 | TG | 0.22086208 | Physical Interactions | Dyer-Sobral-2010 |
| ARSB | GIPC1 | 0.46740454 | Physical Interactions | Dyer-Sobral-2010 |
| NSF | KIAA0917 | 0.22967176 | Physical Interactions | Wan-Emili-2015 |
| SNAP29 | NSF | 0.24036162 | Physical Interactions | Wan-Emili-2015 |
| SNAP29 | STX7 | 0.17399624 | Physical Interactions | Wan-Emili-2015 |
| TBK1 | TNIP1 | 0.032397024 | Physical Interactions | IREF-SMALL-SCALE-STUDIES |
| ATXN2 | parkin | 0.010089592 | Physical Interactions | IREF-SMALL-SCALE-STUDIES |
| A2BP1 | ATXN2 | 0.48912728 | Physical Interactions | IREF-SMALL-SCALE-STUDIES |
| STX8 | NSF | 0.040670313 | Physical Interactions | IREF-SMALL-SCALE-STUDIES |
| GOSR1 | NSF | 0.047341432 | Physical Interactions | IREF-SMALL-SCALE-STUDIES |
| ERGIC3 | ERGIC1 | 1 | Physical Interactions | IREF-SMALL-SCALE-STUDIES |
| CNTNAP1 | NFASC | 0.12418162 | Physical Interactions | IREF-SMALL-SCALE-STUDIES |
| VTI1B | STX8 | 0.14363973 | Physical Interactions | IREF-SMALL-SCALE-STUDIES |
| DOCK2 | ELMO1 | 0.22078465 | Physical Interactions | IREF-SMALL-SCALE-STUDIES |
| STX18 | BNIP1 | 0.16178823 | Physical Interactions | IREF-SMALL-SCALE-STUDIES |
| STX18 | KIAA0917 | 0.15913051 | Physical Interactions | IREF-SMALL-SCALE-STUDIES |
| STX7 | STX8 | 0.04876707 | Physical Interactions | IREF-SMALL-SCALE-STUDIES |
| STX7 | VTI1B | 0.13772634 | Physical Interactions | IREF-SMALL-SCALE-STUDIES |
| STX5 | GOSR1 | 0.0438315 | Physical Interactions | IREF-SMALL-SCALE-STUDIES |
| STX5 | VTI1B | 0.10634396 | Physical Interactions | IREF-SMALL-SCALE-STUDIES |
| SNAP29 | STX12 | 0.08377251 | Physical Interactions | IREF-SMALL-SCALE-STUDIES |
| SNAP29 | STX7 | 0.063081205 | Physical Interactions | IREF-SMALL-SCALE-STUDIES |
| NAPG | NSF | 0.12772426 | Physical Interactions | IREF-SMALL-SCALE-STUDIES |
| NAPG | STX8 | 0.15972765 | Physical Interactions | IREF-SMALL-SCALE-STUDIES |
| TNPO3 | C20orf24 | 0.06513457 | Physical Interactions | Ewing-Figeys-2007 |

| | | | | |
|---|---|---|---|---|
| DNM2 | MCC | 0.03867685 | Physical Interactions | Ewing-Figeys-2007 |
| PFKP | MCC | 0.018131975 | Physical Interactions | Ewing-Figeys-2007 |
| KIAA1267 | FSD2 | 0.026384622 | Physical Interactions | IREF-INTACT |
| TNPO3 | C20orf24 | 0.05292571 | Physical Interactions | IREF-INTACT |
| A2BP1 | TNIP1 | 0.018675499 | Physical Interactions | IREF-INTACT |
| A2BP1 | ATXN2 | 0.03540918 | Physical Interactions | IREF-INTACT |
| STX8 | TG | 0.3206155 | Physical Interactions | IREF-INTACT |
| DNM2 | MCC | 0.003328617 | Physical Interactions | IREF-INTACT |
| ERGIC3 | ERGIC1 | 1 | Physical Interactions | IREF-INTACT |
| CIC | SETD2 | 0.19472425 | Physical Interactions | IREF-INTACT |
| VTI1B | STX8 | 0.05390927 | Physical Interactions | IREF-INTACT |
| PFKP | MCC | 0.014808652 | Physical Interactions | IREF-INTACT |
| TDRD7 | ATXN2 | 0.056049086 | Physical Interactions | IREF-INTACT |
| RBM24 | A2BP1 | 0.39862606 | Physical Interactions | IREF-INTACT |
| SNAP29 | STX12 | 0.23442437 | Physical Interactions | IREF-INTACT |
| IDH3B | ATXN2 | 0.14339492 | Physical Interactions | IREF-INTACT |
| ARSB | GIPC1 | 0.07991065 | Physical Interactions | IREF-INTACT |
| ZNF300 | ZFP3 | 0.002762994 | Predicted | Wu-Stein-2010 |
| CYR61 | CRIM1 | 0.35400745 | Predicted | Wu-Stein-2010 |
| A2BP1 | ATXN2 | 0.28901884 | Predicted | Wu-Stein-2010 |
| DNM2 | MCC | 0.061834827 | Predicted | Wu-Stein-2010 |
| GOSR1 | NSF | 0.1359736 | Predicted | Wu-Stein-2010 |
| ERGIC3 | ERGIC1 | 1 | Predicted | Wu-Stein-2010 |
| CNTNAP1 | CNTN6 | 0.41209117 | Predicted | Wu-Stein-2010 |
| CIC | SETD2 | 0.7494166 | Predicted | Wu-Stein-2010 |
| STX18 | KIAA0917 | 0.45952222 | Predicted | Wu-Stein-2010 |
| STX5 | KIAA0917 | 0.1738322 | Predicted | Wu-Stein-2010 |
| STX5 | VTI1B | 0.17662838 | Predicted | Wu-Stein-2010 |
| RBM24 | A2BP1 | 0.56433994 | Predicted | Wu-Stein-2010 |
| SNAP29 | STX12 | 0.117512956 | Predicted | Wu-Stein-2010 |
| SNAP29 | STX7 | 0.13317996 | Predicted | Wu-Stein-2010 |
